# A computational method for Cell type-specific Expression Quantitative Trait Loci mapping using bulk RNA-seq data

**DOI:** 10.1101/2022.03.31.486605

**Authors:** Paul Little, Si Liu, Vasyl Zhabotynsky, Yun Li, Danyu Lin, Wei Sun

## Abstract

Mapping cell type-specific gene expression quantitative trait loci (ct-eQTLs) is a powerful way to investigate the genetic basis of complex traits. A popular method for ct-eQTL mapping is to assess the interaction between the genotype of a genetic locus and the abundance of a specific cell type using a linear model. However, this approach requires transforming RNA-seq count data, which distorts the relation between gene expression and cell type proportions and results in reduced power and/or inflated type I error. To address this issue, we have developed a statistical method called CSeQTL that allows for ct-eQTL mapping using bulk RNA-seq count data while taking advantage of allele-specific expression. We validated the results of CSeQTL through simulations and real data analysis, comparing CSeQTL results to those obtained from purified bulk RNA-seq data or single cell RNA-seq data. Using our ct-eQTL findings, we were able to identify cell types relevant to 21 categories of human traits.

## Introduction

Studying the variation of gene expression is essential for understanding cellular and molecular biology. Gene expression can vary significantly across different cell types, and that the composition of cell types can vary across tissue samples [1]. As a result, variation in gene expression observed in bulk tissue samples can be due to both cell type-specific expression and variations in cell type compositions [2]. Investigating gene expression quantitative trait loci (eQTLs), or genetic variants associated with gene expression, is a powerful approach for studying the genetic basis of complex traits [3, 4]. Several recent studies found that many genetic loci implicated in human diseases are associated with certain cell types [5–8]. By studying cell type-specific eQTLs (ct-eQTLs), we can gain further insights into the genetic basis of complex traits [9–11].

A popular method to study ct-eQTLs using bulk tissue gene expression data is to include an interaction between the genotype at a genetic locus and the abundance of a cell type in a linear model [3, 4, 11–17]. However, linear models require the residual variation in gene expression to be constant across samples. Therefore, it is often necessary to use a log transformation or normal quantile transformation of RNA-seq count data to stabilize variance. These transformations can result in nonlinear relationships between transformed gene expression and cell type proportions, leading to a mis-specified linear model. An alternative and more appropriate modeling approach is to use negative binomial regression to directly model RNA-seq count data. In addition to total read count (TReC), RNA-seq data can also provide information about allele-specific expression (ASE). By incorporating both TReC and ASE, it is possible to increase the power of eQTL mapping by taking advantage of the allelic imbalance of gene expression caused by *cis*-acting eQTLs. We have developed a method called TReCASE that uses this approach [18, 19]. Most local eQTLs are *cis*-eQTLs, and the terms are often used synonymously. In this paper, we use the term “*cis*-eQTL” to refer specifically to *cis*-acting eQTLs that lead to allelic imbalance.

We have previously developed a method called pTReCASE for eQTL mapping using RNA-seq data from tumor samples, where we treated tumor and non-tumor cells as two distinct cell types with known composition [20]. However, this approach is limited to situations where there are only two cell types and where cell type proportions vary significantly across samples. It treats bulk TReC as the sum of TReC from the two cell types. In more general situations with an arbitrary number of cell types, there are several challenges to ct-eQTL mapping. For example, a cell type may have nearly constant proportions across samples, which can make it difficult to accurately estimate the ct-eQTL effect for that cell type. Additionally, a gene’s expression may be zero or very low in some cell types, making it difficult or impossible to estimate eQTL effects in those cell types. In this paper, we have designed a flexible and robust computational framework from scratch to handle these challenges.

Although single cell RNA-seq (scRNA-seq) data have become more widely available and can be used to study ct-eQTLs, there are still some limitations. First, scRNA-seq is expensive for studies with large sample sizes, and it also requires high quality samples. Additionally, scRNA-seq may not provide a representative sampling of all cell types in a tissue sample, and the inherent sparsity of scRNA-seq data can make it difficult to accurately assign cell types to individual cells. Our method allows for a new study design: collecting scRNA-seq data from a subset of samples, along with bulk RNA-seq data from all samples. The scRNA-seq data can be used to create a cell type-specific gene expression reference, and then the bulk RNA-seq data can be used for ct-eQTL mapping after estimating cell type proportions using the reference.

## Results

### A brief introduction of CSeQTL and OLS method

Our method is called **C**ell type-**S**pecific **eQTL** or CSeQTL for short. CSeQTL jointly models total read count (TReC) and allele-specific read count (ASReC) as a function of covariates, cell type composition, and the genotype at a single nucleotide polymorphism (SNP). More specifically, TReC and ASReC are modeled by a negative binomial and a beta-binomial distribution, respectively, with shared parameters for genetic effects [18]. Unlike the TReCASE and pTReCASE methods, CSeQTL is designed to handle challenging situations where cell type-specific gene expression may be zero or very low, or the proportion of one or more cell types may be close to zero or lack variation. These challenges can make it difficult or impossible to accurately estimate eQTL effects. We address these issues by using several computational solutions, including trimming outliers of TReC to increase the robustness of our estimates and iteratively detecting and removing non-expressed cell types.

We compare CSeQTL to a linear model approach that we refer to as the ordinary least squares (OLS) method. To implement the OLS method, we first apply an inverse normal quantile transformation to read-depth normalized TReC for each gene. Next, we define a reference cell type (usually the one with the highest average abundance) and fit a linear model with transformed gene expression as the dependent variable and the following covariates as independent variables: the proportions of all cell types except the reference cell type, the genotype at a SNP, and the interactions between the SNP genotype and the proportion of each non-reference cell type. Other covariates such as age, sex, and batch can also be included. With this model setup, the ct-eQTL effect for the reference cell type is the main effect of the SNP genotype, and the ct-eQTL effect for a non-reference cell type is the sum of the genotype’s main effect and the effect size of the corresponding interaction term. This model is the same as the one used by Aguirre-Gamboa et al. [16].

### A CSeQTL controls type I error and has much higher power than OLS

We conducted simulations to evaluate type I error and power of CSeQTL in a variety of settings. First, we varied the baseline expression (i.e., gene expression of the reference allele) across cell types. Second, we considered three scenarios of cell type composition variation for three cell types, referred to as CT1, CT2, and CT3 (Figure 1(a)). In scenario 1, cell type proportions were generated independently and identically distributed, and then normalized to sum to one. This scenario represents an ideal, but unrealistic, situation. In scenario 2, we created more realistic cell type proportions by setting the average abundance of CT3 to be lower than CT1 and CT2, and by reducing the variance in the proportion of CT3. This scenario represents a more difficult situation for ct-eQTL mapping of CT3. In scenario 3, we added outlier proportions to the simulated proportions of scenario 2 to mimic observations in real data. We also conducted a secondary set of simulations to explore the performance of CSeQTL given noisy estimates of cell type proportions (Supplementary Figure 1).

**Figure 1:**
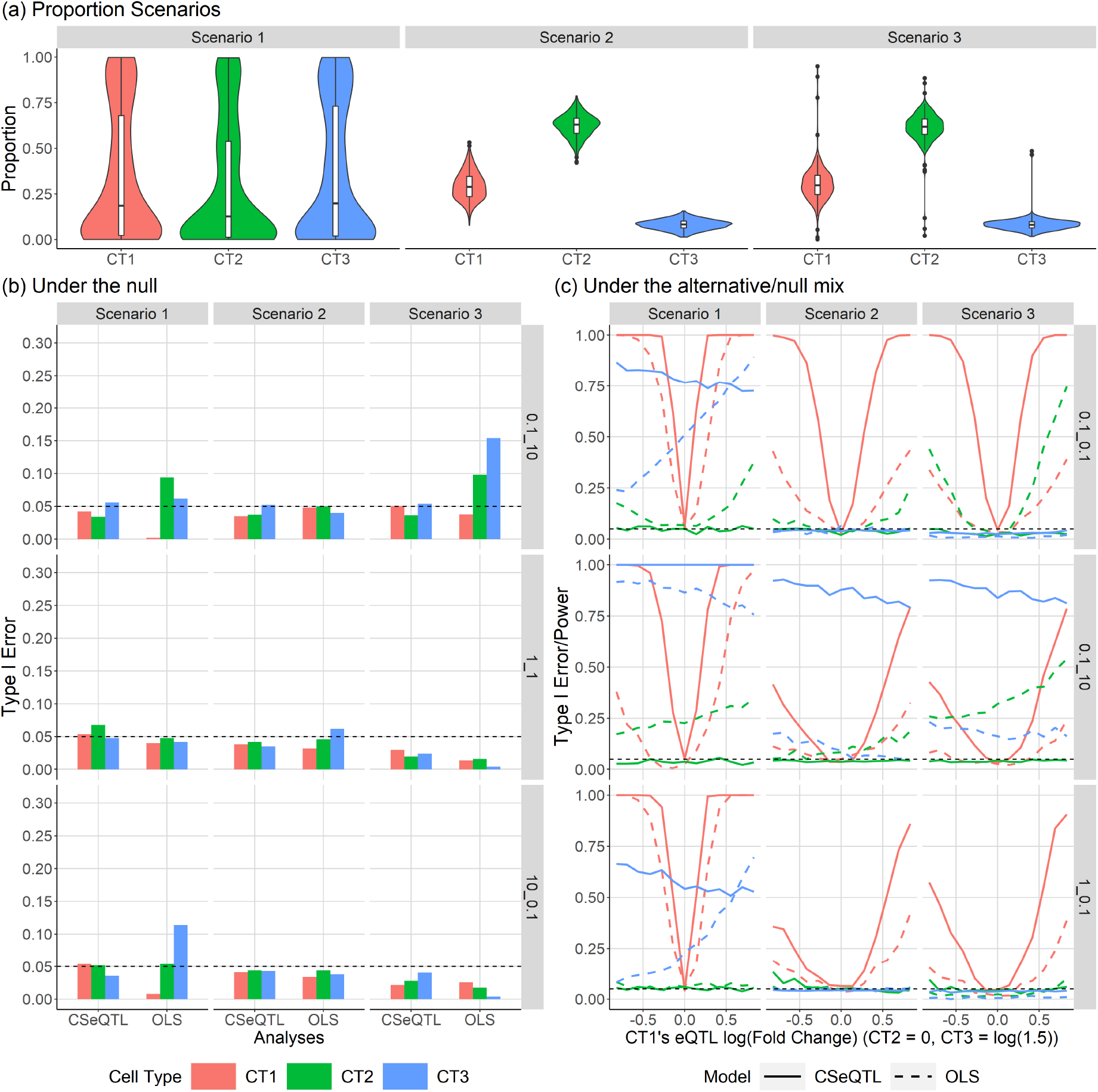
Summary of the results from simulation studies. (a) Simulated cell type proportions for three scenarios. Scenario 1: equally abundant and highly variable cell type proportions. Scenario 2: variable abundance and smaller variance. Scenario 3: modification of scenario 2 by adding outliers of cell type proportions. (b) Simulation results under the global null (i.e., no eQTL for any cell type) for different scenarios and methods (columns of the plots) and reference allele expression configurations (rows of the plots). Each reference allele configuration is denoted by fold change of reference allele expression. For example, “10 0.1” indicates fold changes of 10 in CT2 over CT1 and 0.1 in CT3 over CT1. (c) Simulation under the mixture of null and alternative hypothesis by models, scenarios, and reference allele expression configurations. Results of (b) and (c) are obtained after trimming outliers.

We set the mean expression of the reference allele in CT1 be 500. For example, if the reference/alternative allele is A/T, then the mean expression in an individual with genotype AA is 1000. We set the ASReC to be 5% of the TReC. We also set the fold change for the reference allele gene expression of CT2 or CT3 versus CT1 to be 0.1, 1.0, or 10. Following the TReCASE model, TReC and ASReC were simulated conditional on phased SNP genotypes, cell type proportions, expected expressions per allele and cell type, and other covariates. The sample size was 300. All the eQTLs were set to be *cis*-eQTLs that influenced both TReC and ASReC and we specified eQTL effect size by fold change of alternative allele B versus reference allele A. In a global null situation, all ct-eQTL effects were set to be 1.0 (Figure 1(b)). In another mixed null/alternative situation, we allowed CT1’s eQTL effect to vary from exp(*−*1) to exp(1), set the eQTL effect for CT2 to be 1 (i.e., no eQTL effect), and set the eQTL effect for CT3 to be 1.5. This design allowed us to assess power in CT1 and CT3 and type I error in CT2 simultaneously (Figure 1(c)).

Under both global null and mixed null/alternative situations, CSeQTL controls type I error but OLS has apparent type I error inflation in several configurations (Figure 1(b-c)). Focusing on the mixed null/alternative situation, we found that under scenario 1, when the three cell types have the same distribution of proportions, CSeQTL generally has higher power than OLS. When the baseline expression of CT3 is low (0.1 fold of CT1), OLS’s power in CT3 is positively correlated with CT1’s eQTL effect size even though CT3 has a constant effect size throughout. This “leaking” of eQTL effect from CT1 to CT3 is likely due to the transformation of gene expression. OLS also suffers from inflated type I error (i.e., eQTL findings from CT2) in cases where CT2 has lower baseline expression, highlighting the difficulty in estimating eQTL effects when cell type-specific gene expression levels are low.

In scenario 2, where CT1 has the highest proportion and CT3 has the lowest proportion, power to detect ct-eQTLs is reduced across models and cell types when compared with scenario 1. CSeQTL’s power to detect CT1 eQTLs is much higher than OLS. CT3 eQTLs are detectable by either method if its baseline expression is high and in that case (2nd row and 2nd column of Figure 1(c)) CSeQTL has much higher power than OLS, e.g., *>* 80% power by CSeQTL vs. *<* 20% power by OLS. OLS still has type I error inflation for CT2, to a smaller degree than in scenario 1. Finally in scenario 3, the introduction of outliers in cell type proportions substantially increases the type I error inflation of OLS for CT2 when baseline expression of CT2 is low. Additional simulation results, including results using a noisy version of cell type proportions, are presented in Supplementary Materials (Supplementary Figures 2-5).

Our implementation allows for the trimming of outliers whose Cook’s distance is larger than a threshold, following the approach used by DESeq2 [21]. The Cooks’ distance is calculated based on the null model (no eQTL), and the value of outliers are imputed with the null model’s predicted outcome (See Online Methods Section 3 for more details). This trimming procedure may slightly reduce power, but helps to guard against type I error. The impact of trimming is more apparent in one dataset (GTEx brain samples) in our real data analysis, which we discuss in the next section.

In summary, power to detect ct-eQTLs is driven by the model and positively correlated with eQTL effect size, absolute and relative reference allele expression, and variability in cell type proportions.

### A CSeQTL identifies many more ct-eQTLs than OLS in human brain and blood

We analyzed bulk RNA-seq data from three sources: 670 whole blood samples from the Genotype Tissue Expression (GTEx) project [3], 254 schizophrenia patients and 283 controls from the CommonMind Consortium (CMC) [22, 23], and 175 brain samples from GTEx. Additionally, we studied cell type-purified bulk RNA-seq data from the BLUEPRINT cohort, including purified CD4+ T cells (n=212), monocytes (n=197), and neutrophils (n=205). For the purified bulk RNA-seq data, CSeQTL was equivalent to TReCASE, and the results were used to validate ct-eQTL results from GTEx whole blood samples.

We obtained phased genotypes, TReC, ASReC, observed covariates, and latent batch covariates for each of the four cohorts (GTEx whole blood, CMC brain, GTEx brain, and BLUEPRINT). See Supplementary Materials Section 4 for more information. Using ICeDT [24], we estimated cell type proportions based on TReC and cell type-specific reference data for 5 brain cell types [25] and 22 blood cell types [26]. See Supplementary Materials Section 2 for more details.

We found that the distributions of cell type proportions were similar between schizophrenia patients and healthy controls in CMC and GTEx brain samples (Figure 2(a)). Excitatory neurons (Exc) had the highest proportions, followed by astrocytes (Astro), inhibitory neurons (Inh), oligodendrocytes (Oligo), and oligodendrocyte precursor cells (OPC). Microglia had the lowest proportions and the smallest variation, making it difficult to detect ct-eQTLs in this cell type. For the 22 blood cell types [26], we collapsed them into 7 cell types due to limited prevalence and variability in some cell types (Supplementary Figure 8 and Section 5 of online methods). In GTEx whole blood samples, neutrophils were the dominant cell type with the highest proportions and the largest variance (Figure 2(b)).

**Figure 2:**
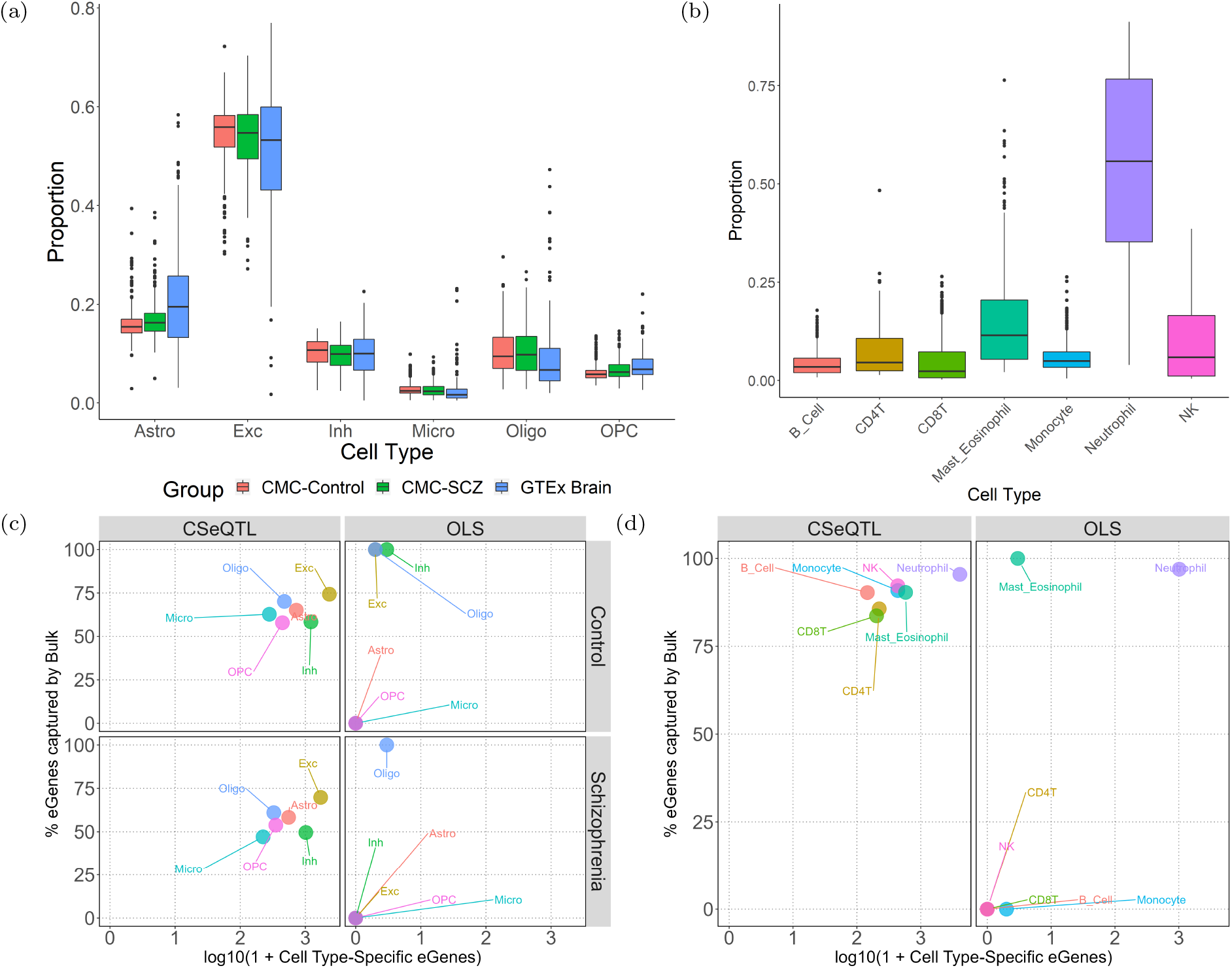
Summary of cell type compositions and the number of eGenes. (a) Cell type proportion estimates of six cell types astrocytes (Astro), excitatory neuron (Exc), inhibitory neuron (Inh), microglia (Micro), oligodendrocytes (Oligo), and oligodendrocyte precursor cells (OPC) from the brain samples of schizophrenia patients and controls from the CommonMind Consortium (CMC) as well as samples from GTEx Brain. (b) Cell type proportion estimates of 7 cell types from whole blood samples of GTEx. Cell type proportions were first estimated for 22 cell types (Supplementary Figure 8) and then collapsed to 7 cell types to avoid individual cell types with very low proportions and variances. (c) A summary of the number of detected eGenes per stratum by method and case/control status for CMC. (d) A summary of detected eGenes by method for GTEx whole blood. For (c) and (d), the X-axis is the (number of cell type-specific eGenes + 1) in log (base 10) scale. The Y-axis denotes the percentage of cell type-specific eGenes that overlap with eGenes from bulk eQTL mapping.

We conducted both ct-eQTL mapping and traditional bulk eQTL mapping, which assesses aggregated eQTL effects in bulk tissue samples. For both ct-eQTL and bulk eQTL mapping, we included a set of covariates: library size, observed covariates (such as age, sex, and known batch effects), and genotype principal components (PCs). We also added latent factors estimated from gene expression data, which were calculated by PCs of residualized gene expression data after accounting for all the aforementioned covariates. We obtained two sets of residualized TReC PCs: the first set was generated by residuals that did not account for cell type proportions and the second set did. The first set was used for bulk eQTL mapping, to mimic the common practice of eQTL mapping. The second set was used in ct-eQTL mapping. When using OLS for ct-eQTL mapping, we included cell type proportions and interaction terms between genotype and cell type proportions. We excluded genes with low expression in most samples (75th percentile of TReC less than 50) and SNPs with minor allele frequencies below 5% from our analysis. We considered SNPs located between 50 kilobases before the transcription start site and 50 kilobases after the transcription end site, including those within the gene body.

We trimmed expression outliers of each gene using Cook’s distance. To determine the appropriate threshold of Cook’s distance for each dataset, we ran TReC-only eQTL mapping using permuted data for all the genes on chromosome 1 with thresholds of 10, 15, 20, or no trimming. We selected the threshold for each dataset to ensure that type I error was controlled per cell type. The selected thresholds were 10 for GTEx brain data and 20 for the other three cohorts. A more aggressive trimming threshold was needed for GTEx brain data, likely due to its smaller sample size.

For eQTL mapping, we need to account for two layers of multiple testing: (1) testing across multiple local SNPs per gene and (2) testing across genes. For each gene, we assessed the significance of its minimum p-value across all local SNPs by calculating the corresponding permutation p-value. A brute-force implementation, which involves permuting the data many times and running CSeQTL on each permuted dataset, is computationally prohibitive. Instead, we used a computationally efficient method called geoP [19, 27] to calculate a permutation p-value by estimating the effective number of independent tests. After this step, each gene has one permutation p-value. To account for multiple testing across genes, we selected a permutation p-value cutoff to control false discovery rate (FDR) quantified by q-value (Supplementary Materials, Section 3). We calculated a q-value [28] for each permutation p-value cutoff and chose a q-value cutoff 0.005 by default. This cutoff was smaller than typical FDR cutoff (e.g., 0.05) because the calculation of q-value accounted for the proportion of nulls, which could lead to a liberal permutation p-value cutoff when the proportion of nulls was small. For bulk eQTL results, a q-value of 0.005 corresponds to permutation p-value around 0.02 while a q-value of 0.05 may correspond to a permutation p-value larger than 0.1 (Supplementary Tables 7-10). We applied this two-step multiple testing correction procedure to both bulk eQTL mapping and ct-eQTL mapping for each cell type. A similar procedure has been used in GTEx studies [3].

When performing bulk eQTL mapping or ct-eQTL mapping using cell type-purified samples from BLUEPRINT, CSeQTL is equivalent to the TReCASE method [29]. Consistent with our previous results [19, 29], CSeQTL has much higher power than OLS. For example, considering the results for CMC schizophrenia samples (n=250), for bulk eQTL mapping after trimming outliers, CSeQTL and OLS identified around 6,900 and 2,900 eGenes (genes with at least one significant eQTL) respectively (Supplementary Table 2). Similar results were observed for BLUEPRINT data from purified blood cell types (Supplementary Table 4 and Supplementary Figures 17-18).

CSeQTL identified many more ct-eQTLs than OLS for different brain cell types. After trimming, CSeQTL identified hundreds to thousands of ct-eQTLs per cell type in CMC schizophrenia data and OLS only identified two eQTLs in oligodendrocytes (Figure 2(c), Supplementary Table 2). The results were similar for CMC control samples (n=275) and trimming outliers did not have a large impact (Supplementary Table 2).

In contrast, trimming outliers had a large effect for GTEx brain data, which had a relatively small sample size of 174. In particular, for microglia, the cell type with the lowest abundance, CSeQTL and OLS identified 885 and 184 eGenes before trimming, but only 96 and 0 eGenes after trimming (Supplementary Figure 14 and Supplementary Table 3). The results from GTEx brain data suggest that CSeQTL still has much higher power than OLS when sample size is small, but should be used with caution.

For blood ct-eQTLs estimated by GTEx whole blood data, OLS identified 1,014 eGenes in neutrophil, and two or zero eGenes in other cell types. In contrast, CSeQTL identified *>*4,000 eGenes in neutrophil, including most findings by OLS (Supplementary Figure 21) and hundreds of eGenes in other cell types (Figure 2(d) and Supplementary Table 5).

CSeQTL results demonstrated limited eGene overlaps across cell types (Supplementary Figures 12, 15, and 20), though the majority of ct-eQTLs overlap with the eQTLs detected by bulk eQTL mapping. These results suggest ct-eQTL signals may be detectable from bulk tissue samples, though without knowing the relevant cell types. Very low consistency between ct-eQTLs and bulk eQTLs may indicate false discoveries in ct-eQTLs. For example, for the GTEx brain study, before trimming, OLS identified 1,332 eQTLs in microglia for 184 eGenes, while only *<*0.01% overlap with bulk eQTLs, and none of these 1,332 eQTLs remained significant after trimming outliers (Supplementary Table 3). In all comparisons hereafter, we focused on the eQTL results after trimming outliers since earlier results demonstrated it could reduce the number of false positives.

### A CSeQTL findings have significant overlaps with ct-eQTLs identified by purified bulk RNA-seq data or scRNA-seq data

We validated the CSeQTL findings from GTEx whole blood using the eQTLs identified from purified bulk RNA-seq data of three cell types - CD4T, monocyte, and neutrophils - from the BLUEPRINT project [30]. A large number of eGenes were identified from BLUEPRINT data and the number was imbalanced across cell types (Supplementary Table 6). In order to make a meaningful comparison, we compared the CSeQTL findings to the top 500 eGenes (*<* 5% of all genes considered by BLUEPRINT) for each of the three BLUEPRINT cell types. At a q-value cutoff of 0.005 for any fold changes, around 35%, 17%, and 12% of CSeQTL eGenes from neutrophil, CD4T, and monocytes overlapped with the top 500 BLUEPRINT eGenes, respectively. These proportions increased to 40%, 30%, and 20% for q-value *<* 0.001 and fold change *≥* 1.5 (Figure 3(a-b)). The numbers of overlaps were 5.7-8.8 times of the numbers expected by chance (Figure 3(c)). Higher overlapping proportion in neutrophil was expected because it was the most abundant cell type and CSeQTL had higher power for the more abundant cell type.

**Figure 3:**
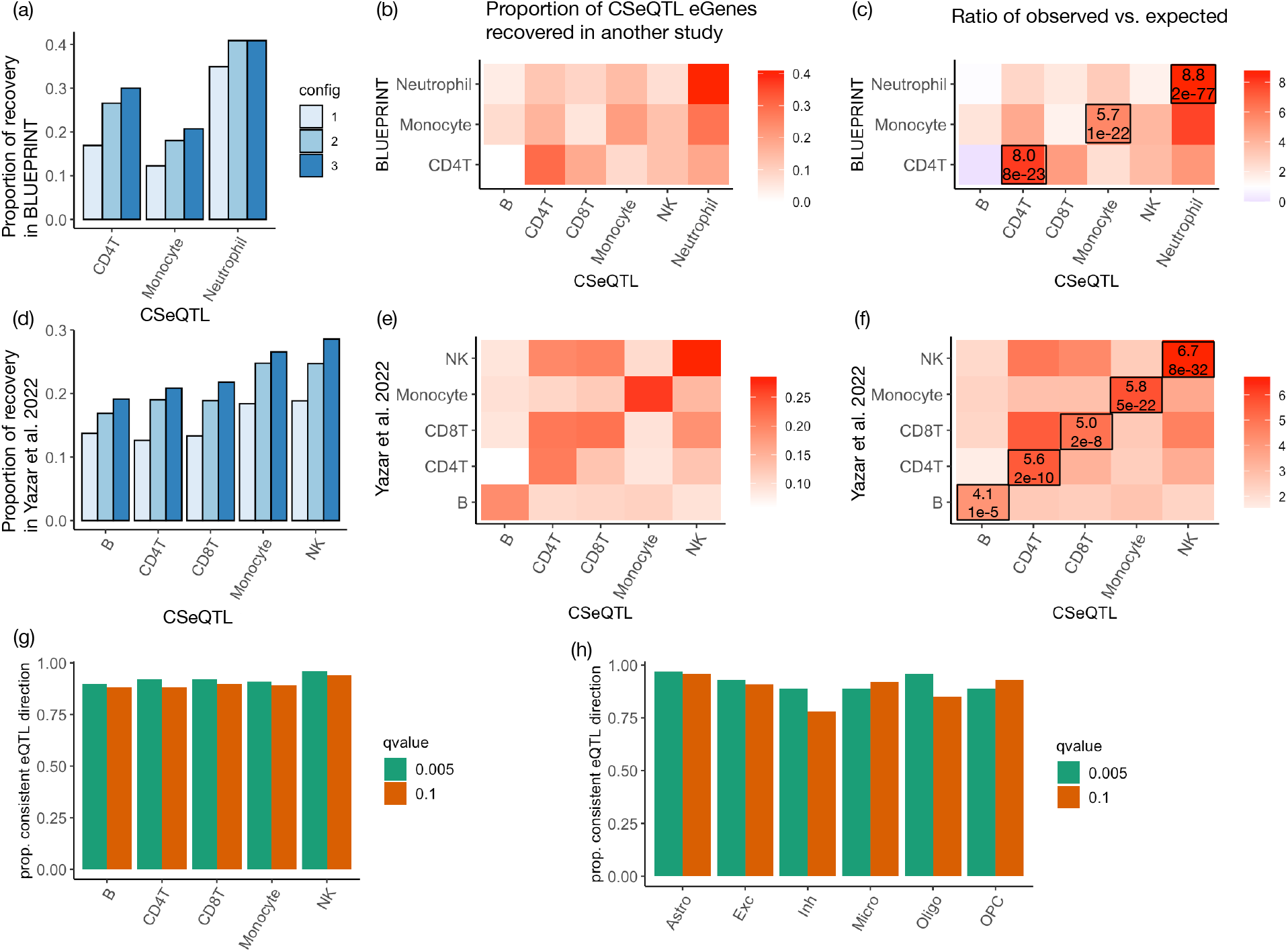
Validation of CSeQTL results. We compare the results by CSeQTL versus the results from cell type purified bulk RNA-seq data (BLUEPRINT) [30] or scRNA-seq data from blood [31] or brain [32]. (a) and (d): the proportion of CSeQTL eGenes recovered from the top 500 (*<* 5%) eGenes of other studies, using three configurations: (1) q-value *<* 0.005; (2) q-value *<* 0.005 and fold change *≥*1.5; and (3) q-value *<* 0.001 and fold change *≥*1.5. (b) and (e): illustration of the recovered eGene proportions at configuration 3 for all cell type pairs. (c) and (f): for each pair of cell types studied by CSeQTL versus BLUEPRINT or Yazar et al., we evaluated the ratio of the observed number of overlapping eGenes versus its expected value. The ratios and corresponding p-values by Fisher’s exact test are labeled for each pair of matched cell types. (g-h) The proportion of eQTL that have consistent directions by comparing CSeQTL results (top eQTL per gene with q-value cutoff 0.1 or 0.005) versus two scRNA eQTL studies with p-value cutoff 0.01.

We also compared the CSeQTL results from GTEx whole blood with the ct-eQTLs identified from a large scRNA-seq dataset [31] from peripheral blood mononuclear cells (PBMCs) of 982 donors, with an average of 1,291 cells per donor. Yazar et al. [31] studied ct-eQTLs in 14 types of immune cells. We removed two cell types with very low proportions and very small number of eGenes. The remaining 12 cell types were collapsed to five categories: B, CD4T, CD8T, Monocyte, and NK (Natural Killer), matching the cell types studied in GTEx whole blood data. This was a challenging comparison because the five cell types had small proportions in whole blood samples, where the most abundant cell type was neutrophil (Figure 2(a-b)). Nevertheless, we found highly significant overlaps between CSeQTL eGenes and Yazar et al. eGenes, with fold change enrichments ranging from 4.1 to 6.7 (Figure 3(d-f)). The fact that more stringent criteria to select ct-eQTLs lead to larger overlap proportions (Figure 3(a) and (d)) suggests that our quantification of ct-eQTL effect sizes and significance levels is useful to select stronger ct-eQTLs. CD4T is the most abundant cell type studied by Yazar et al. [31], though the replication percentage is lower than most other cell types, likely due to two reasons. First, its proportion is low in whole blood samples (Figure 2(b)). Second, similarity between CD4T cells and other cell types, such as CD8T cells, may lead to reduced accuracy of cell type deconvolution in bulk RNA-seq data as well as cell type classification in scRNA-seq data. Another important criterion to evaluate eQTL findings is the consistency of eQTL effect directions. We examined the ct-eQTLs that were identified by both CSeQTL (using a q-value cutoff 0.1 or 0.005) and scRNA-seq data (p-value *<* 0.01), and found the eQTL directions were consistent for more than 90% of ct-eQTL findings across most cell types (Figure 2(g)). In contrast, without applying any q-value/p-value filtering the consistency proportion is 51% (Supplementary Table 11).

For the ct-eQTLs identified from brain samples (GTEx brain, CMC schizophrenia patients or controls), we compared with ct-eQTLs reported by a single nucleus RNA-seq (snRNA-seq) study [32]. Bryois et al. [32] collected snRNA-seq data for 6,940 to 14,595 genes in 144 to 192 individuals for eight major brain cell types: excitatory neurons, inhibitory neurons, astrocytes, microglia, oligodendrocytes, oligodendrocyte precursor cells (OPCs), Endothelial, and Pericytes. Both Pericytes and Endothelial had very small number of cells and ct-eQTLs, and thus we skipped them in our comparison. The remaining six cell types were exactly the same as the cell types considered in our CSeQTL analysis. Overall the results were consistent with the findings for immune cell types. The CSeQTL eGenes had significant overlap with the top eGenes reported by Bryois et al. (Supplementary Figure 22). Though the overlap was low for two cell types: inhibitory neurons and microglia, likely due to low proportions of these two cell types. In addition, cell type-specific expression were similar between excitatory neurons and inhibitory neurons (Supplementary Figure 9), which could further increase the difficulty to map ct-eQTLs for inhibitory neurons. The eQTL effect direction estimates by CSeQTL and snRNA-seq were highly consistent for most cell types except for inhibitory neurons, again suggesting that ct-eQTL mapping was challenging for this cell type (Figure 3(h), Supplementary Table 12-14).

We further compared the ct-eQTLs identified only by scRNA-seq/snRNA-seq data or only by CSeQTL on bulk RNA-seq data. The scRNA-seq-only ct-eQTLs tended to have smaller effect sizes and larger p-values. Therefore CSeQTL may have missed those scRNA-seq-only ct-eQTLs because of their weaker effects (Supplementary Figure 23). CSeQTL combines deconvolution of gene expression and eQTL mapping into one step which accounts for the uncertainty of deconvolution. Therefore, the power of CSeQTL is impacted by both the uncertainty of gene expression deconvolution and the magnitude of the ct-eQTLs. The ct-eQTLs identified solely by CSeQTL tended to have smaller effect sizes and higher expression levels in bulk samples (Supplementary Figure 24). This makes sense because genes with higher expression have smaller uncertainty in gene expression deconvolution. Higher gene expression level should also increase the power of eQTL mapping using scRNA-seq data, though its effect could be more pronounced for CSeQTL as it also improves the accuracy of cell type deconvolution.

### A Characterization of ct-eQTLs

We have analyzed two blood RNA-seq data sets (BLUEPRINT and GTEx whole blood) and three brain RNA-seq data sets (CMC schizophrenia (SCZ), CMC control, and GTEx brain). It is interesting to study the consistency of eQTL results across datasets. Overall, CSeQTL results showed a higher level of consistency than OLS results (Supplementary Figure 25, Supplementary Table 6). For whole blood, a higher consistency was observed for neutrophil, likely due to its higher abundance. For brain datasets, CMC-SCZ and CMC-Control showed higher levels of consistency than between CMC dataset and GTEx brain, likely due to batch effects between the two studies.

We summarized the locations of the minimum p-value SNPs (minP-SNPs) relative to the corresponding eGenes (Supplementary Figures 11, 13, 16, 19). In the brain datasets, The locations of minP-SNPs from bulk eQTL mapping showed enrichment around the transcription start site (TSS) or transcription end site (TES), though such patterns were not as clear in ct-eQTLs. A potential reason was that the eQTLs around TSS and TES were more likely to be shared across cell types. In GTEx brain results, more ct-eQTLs tended to be located further away from the corresponding eGenes, which might be due to the limited sample size hence higher uncertainty to locate the eQTLs. For the three purified cell types from BLUEPRINT (Supplementary Figure 16), the enrichment of eQTLs around TSS was stronger than TES. Similar patterns of eQTL locations were observed for the same three cell types in GTEx whole blood samples (Supplementary Figure 19).

Next we focused on CSeQTL results and evaluated the distribution of ct-eQTLs with respect to functional annotations of genomic regions (e.g., enhancers, promoters, 3’ UTR, 5’ UTR, etc.) by Torus [33] (Supplementary Figures 26-27). For brain tissues, the functional enrichment of eQTLs for excitatory neuron, which was the most abundant cell type, was similar to the functional enrichment of bulk eQTLs. The lack of significant functional enrichment in other cell types could be partially due to smaller number of ct-eQTLs. Comparing brain samples of schizophrenia patients versus controls (either CMC controls and GTEx controls), enrichment of eQTLs at 5’ UTR and non-coding (NC) transcript were observed in both control groups but were absent in schizophrenia patients. Since neutrophil was the dominant cell type in whole blood, as expected, functional enrichment in neutrophil and whole blood was highly consistent. Despite the small proportions of CD4+ T and monocyte in whole blood, CSeQTL results recovered similar functional enrichment as those observed in purified cell types from BLUEPRINT data.

### A CSeQTL helps interpret GWAS findings

EQTLs are often used to study the genetic basis of complex traits by examining their overlap with genetic loci identified from Genome-Wide Association Studies (GWAS). Here we systematically evaluated the overlap between ct-eQTLs and GWAS hits of either all the traits included in the GWAS catalog [34] on 21 categories of traits (Figure 4 and Supplementary Figure 5). We calculated the enrichment of eQTLs among GWAS hits by a log fold change (the proportion of GWAS hits that overlap with eQTLs versus the proportion of genetic loci being eQTLs). See Section C.3.3 of Vasyl et al. [19] for details on the computation of point estimates and their confidence intervals.

**Figure 4:**
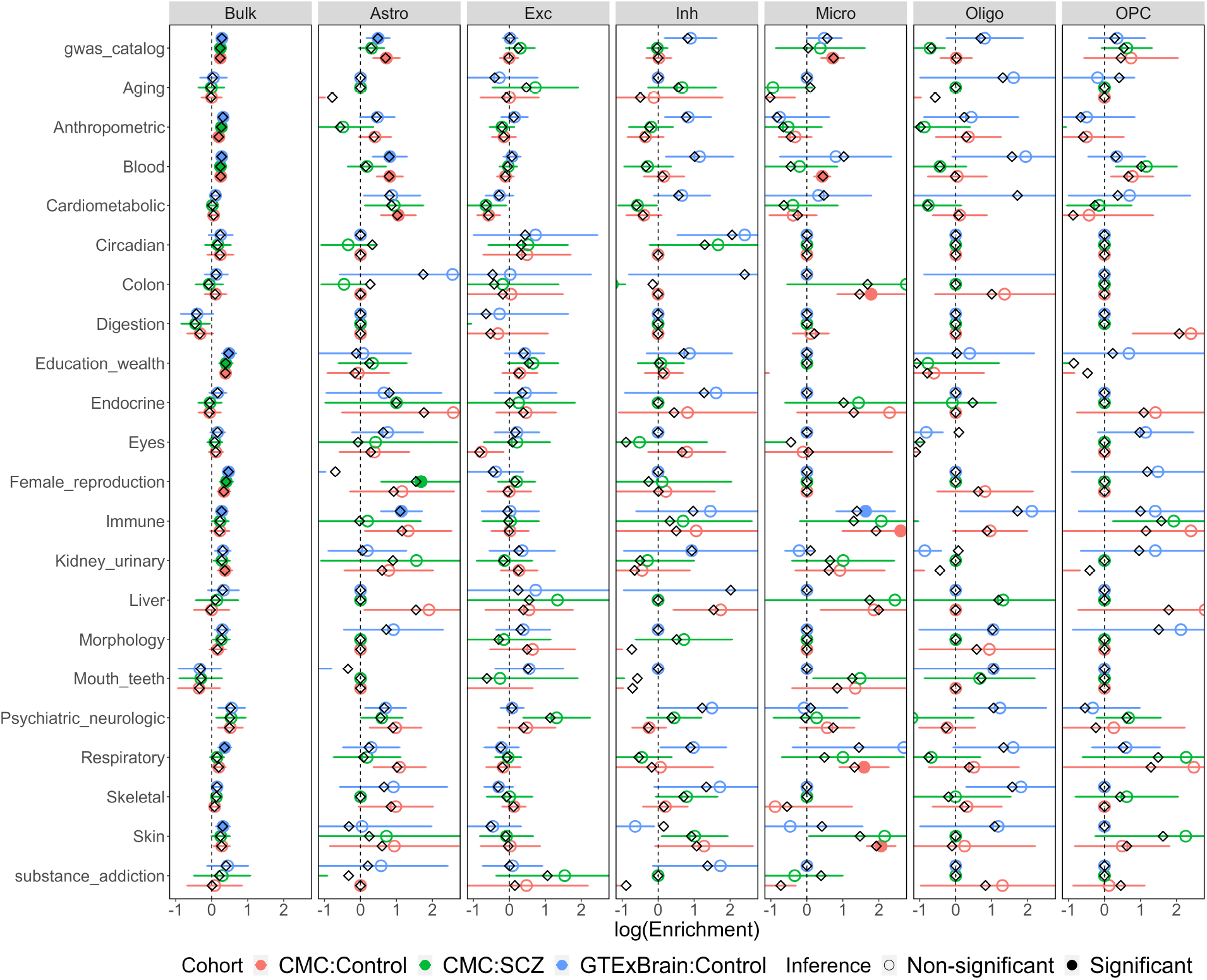
GWAS enrichment for CMC and GTEx Brain. Black diamonds correspond to point estimates of log enrichment of eQTLs among GWAS hits, while open and filled circles correspond to jackknife estimates of log enrichment. Jackknife-based 95% confidence intervals are provided. Intervals are converted to nominal p-values that are then Bonferroni corrected. Filled circles correspond to the ones with lower bound of confidence intervals larger than zero and adjusted p-values *<* 0.05.

GWAS hits of several categories (e.g., education/wealth) were enriched in the bulk eQTLs of all three brain datasets, though the degree of enrichment (measured by log fold change in Figure 4) was small. When considering ct-eQTLs by CSeQTL, due to the smaller number of eQTLs, the confidence to estimate enrichment was often low, which led to wider confidence intervals. Despite such limitation, we observed several interesting findings. For example, the GWAS hits of immune traits were enriched in the ct-eQTLs for microglia in CMC controls and GTEx brain samples, but not in CMC SCZ samples, suggesting potential SCZ-specific and ct-eQTL signals.

Blood is arguably the most accessible tissue and thus molecular biomarkers (e.g., cell type-specific gene expression) in blood can be very valuable to understand the mechanism that connects genetic variants and complex traits. Our ct-eQTL results provided a useful resource for such studies (Figure 5). For example, enrichment of respiratory and skin disease GWAS signals among B cell specific eQTLs, and the association between liver disease GWAS hits and the eQTLs in CD8+ T cells. Earlier studies have reported that CD8+ T cells were associated with liver damage, hepatitis, immunopathology, and liver cancer [35–39].

**Figure 5:**
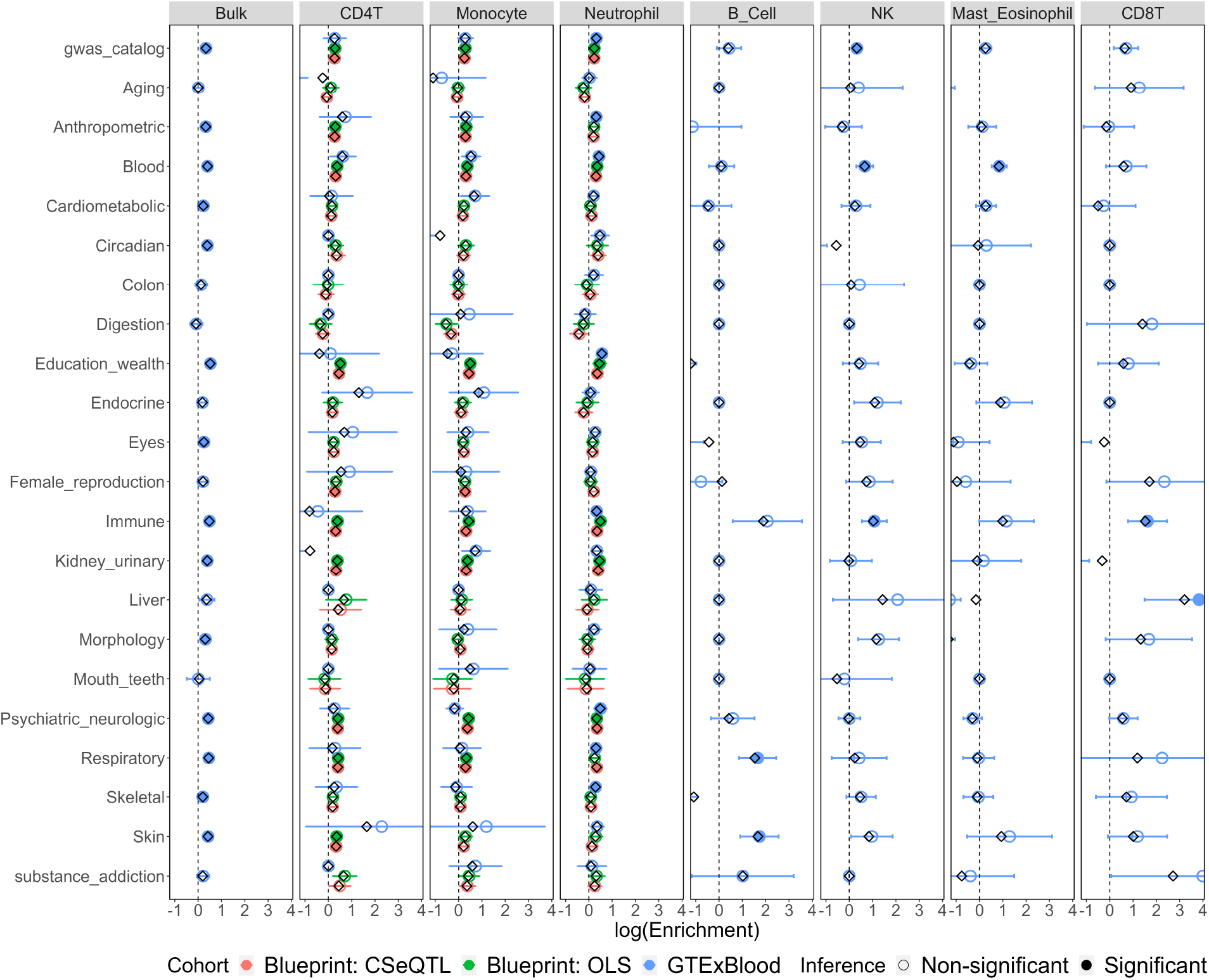
GWAS enrichment for BLUEPRINT and GTEx Whole Blood. Black diamonds correspond to point estimates of log enrichment of eQTLs among GWAS hits, while open and filled circles correspond to jackknife estimates of log enrichment. Jackknife-based 95% confidence intervals are provided. Intervals are converted to nominal p-values that are then Bonferroni corrected. Filled circles correspond to the ones with lower bound of confidence intervals larger than zero and adjusted p-values *<* 0.05.

## Discussion

CSeQTL’s framework allows mapping ct-eQTLs using bulk RNA-seq data, by jointly modeling the effects of cell type composition and ct-eQTLs. We have shown by simulations and real data analyses that CSeQTL can have substantially higher power than a linear regression approach, while still maintaining type I error control. This is due to the underlying statistical model of CSeQTL. Deconvolution of gene expression to individual cell types should be performed using untransformed count data [40], while eQTL mapping is often done using transformed gene expression data (e.g., normal quantile transformation) to avoid the impact of outliers in count data. Such outliers are often due to a strong positive associations between mean value and variance of count data. Our CSeQTL method satisfies these two restrictions by directly modeling count data using a negative binomial distribution that accounts for the strong mean-variance dependence. In contrast, the linear regression approach uses transformed gene expression data and models ct-eQTLs by adding interactions between cell type compositions and genotypes. The transformation of gene expression data distorts their associations with cell type compositions and thus can reduce power and inflate type I error. In addition, we also include allele-specific expression in our model to boost the power to detect ct-eQTLs.

Model optimization of CSeQTL is very challenging because the model may not be identifiable, for example, due to a lack of variation of one cell type’s abundance across individuals or very low expression of one gene in one or more cell types. A naive implementation may result in sub-optimal solutions due to noninvertible observed information matrices, negative variances, or extreme parameter estimates, which can have a profound impact on hypothesis testing. We have developed a comprehensive set of assessments to ensure a robust and optimal solution is obtained. In addition, both linear regression and CSeQTL can be sensitive to outliers, and we addressed this issue by trimming those outliers based on the null model without eQTL effects. As shown in our real data analyses, such trimming can be particularly helpful for studies with smaller sample sizes.

Our applications toward human brain and blood bulk RNA-seq data demonstrate that the linear regression method often identifies none or a few ct-eGenes (with the exception of neutrophil in whole blood) while CSeQTL can identify hundreds to thousands of cell type-specific eGenes. When examining the overlap between these ct-eQTLs and GWAS findings, we have identified several interesting results but with high uncertainty in many cases. Future independent studies and comparisons with larger sample sizes may be needed to reach more definite conclusions.

A limitation when applying our method or any ct-eQTL mapping method on bulk RNA-seq data is accurate estimation of cell type composition, which in turn requires accurate cell type-specific gene expression reference. Here we have applied our method on the bulk RNA-seq data from brain and blood because these two tissues have readily available cell type-specific gene expression reference. We expect that in the near future, with the advance of the human cell atlas [1] or other similar projects, such resources will become available in more tissue types.

Our work also enables a new type of study design to jointly model scRNA-seq and bulk RNA-seq data to study ct-eQTLs. For example, scRNA-seq data can be collected in a small number of individuals, to be used as reference for cell type-specific expression. In addition, scRNA-seq data can also be used for eQTL mapping. After clustering and identification of cell types, scRNA-seq data can be converted to pseudo-bulk data of individual cell types and be used for eQTL mapping, e.g., by applying our TReCASE method [29]. The likelihood function of the TReCASE model can be combined with CSeQTL model in order to combine bulk RNA-seq and scRNA-seq data for ct-eQTL mapping. Adding scRNA-seq data to bulk RNA-seq data can alleviate some challenges when using bulk RNA data, e.g., limited variability in cell type abundance for one cell type. Adding bulk RNA-seq data to scRNA-seq data can reduce the cost, increase sample size, and avoid distortion of gene expression in the process of isolating single cells.

## Author Contributions

WS, DL, YL supervised the project. PL and WS designed the method, acquired and preprocessed the four datasets used in this paper. PL wrote the software package to perform simulation and real data analyses. VZ provided the geoP software. SL and WS provided the validation analyses. PL, SL, and WS wrote the manuscript, with input from DL, YL, and VZ.

## Data Availability

Our work did not generate any new data. We have used publicly available datasets. BLUEPRINT from European Genome Archive with phased SNPs derived from whole genome sequencing (EGAD00001002663) and three purified cell types of RNA-seq bam files (EGAD00001002671, EGAD00001002674, EGAD00001002675). CommonMind data were downloaded from https://www.nimhgenetics.org/resources/commonmind. GTEx data were downloaded from NHGRI AnVIL (Genomic Data Science Analysis, Visualization, and Informatics Lab-space). Brain MTG data were downloaded from Allen Brain Institute Website https://portal.brain-map.org/atlases-and-data/rnaseq/human-mtg-smart-seq. SEA-AD snRNA-seq data were downloaded from cellxgene: https://cellxgene.cziscience.com/collections/1ca90a2d-2943-483d-b678-b809bf464c30.

## Code Availability

The source codes for R package CSeQTL and analysis pipeline are made publicly available at the Github repositories https://github.com/pllittle/CSeQTL and https://github.com/pllittle/CSeQTLworkflow, respectively.

