## Supplementary material for "A computational method for Cell type-specific Expression Quantitative Trait Loci mapping using bulk RNA-seq data": Online Methods

**2 Optimization Scheme and Parameter Assessment** **4**

**3 Trimming Influential Counts** **5**

**4 Simulation Setup** **6**

**5 Grouping 22 blood cell types to 7 cell types** **6**

### 1 Statistical Models

#### 1.1 Notations and the Joint Model of TReC and ASReC

Since our model is the same for any gene-SNP pair, we omit gene and SNP indices to simplify notations. We use $i$ and $q$ as indices for sample and cell type, respectively, where $i = 1, \ldots, n$, $q = 1, \ldots, Q$, and $n$ and $Q$ denotes sample size and the number of cell types, respectively. Let $T_i$ and $N_i$ be the total read count (TReC) and the allele-specific read count (ASReC) mapped in the $i$th sample. Each SNP of interest has two alleles, $A$ and $B$. Each gene has two haplotypes that are arbitrarily defined as haplotype 1 and 2. Let $N_i - N_{i2}$ and $N_{i2}$ denote the ASReC mapped to the first and second haplotypes of sample $i$, respectively.

Let $Z_i$ denote the phased genotype for the SNP in sample $i$, which takes values $AA$, $AB$, $BA$, or $BB$. Let $\boldsymbol{X}_i = (X_{i1}, \ldots, X_{ip})^{\mathrm{T}}$ be a $p$-vector of baseline covariates (excluding the intercept), where T denotes vector or matrix transpose. Among the baseline covariates in our model, we adjust for log-transformed read depth, defined as the log of the 75th percentile of a sample's gene-level TReCs, a more robust measurement of read-depth than summing over all TReC values. Let $\rho_{iq}$ denote the cell type proportion in the $i$th sample and $q$th cell type such that $\sum_{q=1}^{Q} \rho_{iq} = 1$ and $\boldsymbol{\rho}_i = (\rho_{i1}, \ldots, \rho_{iQ})^{\mathrm{T}}$. The cell type corresponding to $q = 1$ is referred to as the reference cell type. Our model is based on the following factorization:

$$P(T_i, N_i, N_{i2} | Z_i, \boldsymbol{X}_i, \boldsymbol{\rho}_i) = P(T_i | Z_i, \boldsymbol{X}_i, \boldsymbol{\rho}_i) \, P(N_i | T_i, Z_i, \boldsymbol{X}_i, \boldsymbol{\rho}_i) \, P(N_{i2} | T_i, N_i, Z_i, \boldsymbol{X}_i, \boldsymbol{\rho}_i)$$

Each factor is defined as follows.

- $P(T_i | Z_i, \boldsymbol{X}_i, \boldsymbol{\rho}_i)$: given $(Z_i, \boldsymbol{X}_i, \boldsymbol{\rho}_i)$, $T_i$ is assumed to follow a negative binomial distribution with mean $\mu_i = E[T_i | Z_i, \boldsymbol{X}_i, \boldsymbol{\rho}_i]$ and dispersion parameter $\phi$ such that $V[T_i | Z_i, \boldsymbol{X}_i, \boldsymbol{\rho}_i] = \mu_i + \phi \mu_i^2$. This likelihood term corresponds to the TReC model.
- $P(N_i | T_i, Z_i, \boldsymbol{X}_i, \boldsymbol{\rho}_i)$: this term describes the total number of allele-specific reads as a function of TReC. It is determined by the number of heterozygous SNPs within the gene and is a constant with respect to the parameters of eQTLs. Thus it is factored out from the likelihood.
- $P(N_{i2} | T_i, N_i, Z_i, \boldsymbol{X}_i, \boldsymbol{\rho}_i)$: given $(N_i, Z_i, \boldsymbol{\rho}_i)$, the read count $N_{i2}$ is assumed to be independent of $(T_i, \boldsymbol{X}_i)$ and follows a beta-binomial distribution with parameter $\pi_i$, which is the expected proportion of ASReC from the haplotype harboring the B allele for heterozygous samples among $N_i$ allele-specific reads, and a dispersion parameter $\psi$. This likelihood term corresponds to the ASReC model.

The above likelihood framework is the same as our TReCASE method that combine TReC and ASE to map *cis*-eQTLs [1, 2, 3]. Similar to TReCASE, we reduce the negative binomial and beta-binomial distribution to Poisson and binomial distribution, respectively, when the data does not support a non-zero overdispersion parameter. Next we describe how to extend each component of the likelihood function for cell type-specific eQTL mapping.

Let $\mu_{i,z,q}$ be the expected TReC for the $i$th sample, $z$th phased genotype, and $q$th cell type. We assume a multiplicative model $\mu_{i,z,q} = \mu_{z,q} \exp\left\{\boldsymbol{X}_i^{\mathrm{T}} \boldsymbol{\beta}\right\}$ and that the effect of baseline covariates $\boldsymbol{\beta}$ are the same for all cell types. We assume that the gene expression for each genotype is the summation of allelic expressions such that $\mu_{AA,q} = \mu_{A,q} + \mu_{A,q} = 2\mu_{A,q}$, $\mu_{AB,q} = \mu_{A,q} + \mu_{B,q} = \mu_{BA,q}$, and $\mu_{BB,q} = \mu_{B,q} + \mu_{B,q} = 2\mu_{B,q}$ where $\mu_{A,q}$ and $\mu_{B,q}$ denote the expected TReC for A and B alleles of the $q$th cell type, respectively. Define $\kappa_q = \mu_{A,q} / \mu_{A,1}$ and $\eta_q = \mu_{B,q} / \mu_{A,q}$ where $\boldsymbol{\kappa} = (\kappa_1, \ldots, \kappa_Q)^{\mathrm{T}}$ and $\boldsymbol{\eta} = (\eta_1, \ldots, \eta_Q)^{\mathrm{T}}$. $\kappa_q$, which is a nuisance parameter, is the fold change of the A allele's expression in the $q$th cell type versus the first cell type. $\eta_q$ is the eQTL effect size: the expression fold change of the B allele vs. A allele for the $q$th cell type.

#### 1.2 Linear Model

To establish a baseline for comparison and mimicking published analyses, we propose fitting a linear model by ordinary least squares (OLS). Let $G_i$ denote the number of B alleles for a given phased SNP for the $i$th sample (i.e.. $G_i = 0$, 1, 1, and 2 for $Z_i = AA$, $AB$, $BA$, and $Z_i = BB$, respectively). For a gene-SNP pair, the cell type-specific linear model is

$$E\left[\bar{T}_i\right] = \zeta_0 + \sum_{j=1}^{p} X_{ij}\zeta_j + G_i\zeta_g + \sum_{q=2}^{Q} \rho_{iq}\gamma_q + G_i \sum_{q=2}^{Q} \rho_{iq}\delta_q$$

where $\bar{T}_i$ is the inverse normal quantile transformation of read-depth adjusted $T_i$. The benefit of the transformation is guarding against outliers on the count scale, and it is a popular choice in eQTL studies [4, 5]. From the above model, we can test $H_0 : \zeta_g + \delta_q = 0$ to assess the strength of cell type-specific eQTL for the $q$th cell type, where $q = 2, \ldots, Q$, and test $H_0 : \zeta_g = 0$ for the reference cell type's eQTL.

#### 1.3 TReC Model

Let $\eta_q^{(\mathrm{T})}$ be the eQTL effect size for the TReC model, where the superscript $^{(\mathrm{T})}$ indicates TReC model. Given the above notations and parameters, let $\mu_{i,AA}$ be the expected TReC for the $i$th sample with $AA$ genotype and it is defined such that

$$\begin{aligned}\log\left(\mu_{i,AA}\right) = \log\left(\sum_{q=1}^{Q} \rho_{iq}\mu_{i,AA,q}\right) &= \boldsymbol{X}_i^{\mathrm{T}}\boldsymbol{\beta} + \log\left(\sum_{q=1}^{Q} \rho_{iq}\mu_{AA,q}\right) \\ &= \log\left(2\mu_{A,1}\right) + \boldsymbol{X}_i^{\mathrm{T}}\boldsymbol{\beta} + \log\left(\sum_{q=1}^{Q} \rho_{iq}\kappa_q\right)\end{aligned}$$

With similar derivations for genotypes $AB$, $BA$, and $BB$, we have

$$\log\left(\mu_i\right) = \begin{cases} \log\left(\mu_{i,AA}\right) & Z_i = AA \\ \log\left(\mu_{i,AA}\right) + \log\left(1 + \xi_i^{(\mathrm{T})}\right) - \log(2) & Z_i = AB, BA \\ \log\left(\mu_{i,AA}\right) + \log\left(\xi_i^{(\mathrm{T})}\right) & Z_i = BB \end{cases} \quad \text{where } \xi_i^{(\mathrm{T})} = \frac{\sum_{q=1}^{Q} \rho_{iq}\eta_q^{(\mathrm{T})}\kappa_q}{\sum_{q=1}^{Q} \rho_{iq}\kappa_q}.$$

It is crucial to notice that $\xi_i^{(\mathrm{T})}$ represents the bulk eQTL effect size for the $i$th sample. If eQTL effect is the same across cell types ($\eta_1^{(\mathrm{T})} = \cdots = \eta_Q^{(\mathrm{T})} = \eta^{(\mathrm{T})}$), $\xi_i^{(\mathrm{T})} = \eta^{(\mathrm{T})}$ simplifying CSeQTL's TReC model to the bulk TReC model presented by Sun (2012) [1]. For $Q = 2$, CSeQTL's TReC model would correspond to pTReCASE's TReC model [3]. After centering continuous covariates among $\boldsymbol{X}_i$ and setting categorical covariates among $\boldsymbol{X}_i$ to their reference level, the intercept term of the above model represents the log-transformed expected TReC of the reference cell type with genotype $AA$. This straightforward interpretation of CSeQTL is a crucial feature for model optimization and parameter estimate interpretation.

#### 1.4 ASReC Model

Let $\eta_q^{(\mathrm{A})}$ be the eQTL effect size associated with the ASReC model. Let $P_{BB}\left(N_1 | N; \pi; \psi\right)$ be the beta-binomial density for observing $N_1$ successes among $N$ trials with success probability $\pi$ and overdispersion parameter $\psi$. For a gene-SNP pair, the ASReC likelihood is defined as

$$P\left(N_{i2} | N_i, Z_i\right) = \begin{cases} 1 & \text{if } N_i = 0 \\ P_{BB}\left(N_{i2} \middle| N_i; \pi_i = \xi_i^{(\mathrm{A})} / \left(1 + \xi_i^{(\mathrm{A})}\right); \psi\right) & \text{if } N_i > 0, Z_i = AB \\ P_{BB}\left(N_i - N_{i2} \middle| N_i; \pi_i = \xi_i^{(\mathrm{A})} / \left(1 + \xi_i^{(\mathrm{A})}\right); \psi\right) & \text{if } N_i > 0, Z_i = BA \\ P_{BB}\left(N_{i2} | N_i; \pi_i = 0.5; \psi\right) & \text{if } N_i > 0, Z_i = AA, BB. \end{cases}$$

Similar to the TReC model,

$$\xi_i^{(\mathrm{A})} = \frac{\sum_q \rho_{iq}\eta_q^{(\mathrm{A})}\kappa_q}{\sum_q \rho_{iq}\kappa_q},$$

which is the bulk eQTL effect size estimated from ASReC. If $N_i = 0$, the ASReC likelihood factors out of the joint model. Furthermore, while samples with genotypes $AA$ and $BB$ do not add information when estimating $\eta_q^{(A)}$ and $\kappa_q$, they contribute toward estimating $\psi$.

##### 1.5 *cis*/*trans* eQTL Testing and eQTL Testing

Following Sun (2012) [1], the model-specific eQTL parameters $\eta_q^{(T)}$ and $\eta_q^{(A)}$ are used to formally characterize *cis* and *trans* eQTLs. By defining $\eta_q^{(A)} = \eta_q^{(T)}\alpha_q$, the $q$th cell type-specific eQTL being *cis* corresponds to $\alpha_q = 1$ and *trans* otherwise. For *cis*-eQTLs, we use the joint model that combines TReC and ASReC/ASE models with shared cell type-specific parameter $\eta_q = \eta_q^{(A)} = \eta_q^{(T)}$. We conduct *cis*/*trans* testing per gene-SNP pair and per cell type with $H_0 : \alpha_q = 1$ vs. $H_A : \alpha_q \neq 1$. Let $\boldsymbol{\alpha} = (\alpha_1, \ldots, \alpha_Q)^{\mathrm{T}}$.

EQTL significance testing is conducted for each gene, SNP, and cell type using either the TReC model for *trans*-eQTL with $H_0 : \eta_q^{(T)} = 1$ vs. $H_A : \eta_q^{(T)} \neq 1$ or the joint model for *cis*-eQTL with $H_0 : \eta_q = 1$ vs. $H_A : \eta_q \neq 1$. Thus our model formulation is flexible enough to allow subsets of cell type-specific eQTLs to be *cis*- or *trans*-eQTLs.

#### 2 Optimization Scheme and Parameter Assessment

Given cell type proportions, our optimization scheme for TReC and ASReC model fitting and hypothesis testing is based on the following procedure for a gene-SNP pair. This scheme helps to avoid local optima since parameter estimation can be sensitive to initialization and influential counts. Let $\boldsymbol{\theta} = (\mu_{A,1}, \boldsymbol{\beta}^{\mathrm{T}}, \phi, \boldsymbol{\kappa}^{\mathrm{T}}, \boldsymbol{\eta}^{\mathrm{T}}, \psi, \boldsymbol{\alpha}^{\mathrm{T}})^{\mathrm{T}}$ denote the pre-established set of unconstrained parameters to optimize over. First, we condition $T_i$ on $\boldsymbol{X}_i$ to obtain $\widehat{\boldsymbol{\theta}}_1 = (\widehat{\mu}_{A,1}, \widehat{\boldsymbol{\beta}}^{\mathrm{T}})^{\mathrm{T}}$ by fitting a Poisson model with Newton-Raphson. Second, we can fit a negative binomial model with initialization $\widehat{\phi} = 1$ to obtain $\widehat{\boldsymbol{\theta}}_2 = (\widehat{\mu}_{A,1}, \widehat{\boldsymbol{\beta}}^{\mathrm{T}}, \widehat{\phi})^{\mathrm{T}}$, also with Newton-Raphson. Third, we incorporate $\boldsymbol{\rho}_i$, initialize $\widehat{\kappa}_q = 1$, and use Broyden–Fletcher—Goldfarb–Shanno (BFGS) to obtain $\widehat{\boldsymbol{\theta}}_3 = (\widehat{\mu}_{A,1}, \widehat{\boldsymbol{\beta}}^{\mathrm{T}}, \widehat{\phi}, \widehat{\boldsymbol{\kappa}}^{\mathrm{T}})^{\mathrm{T}}$. Fourth, we incorporate $Z_i$, initialize $\widehat{\eta}_q = 1$, and use BFGS to obtain $\widehat{\boldsymbol{\theta}}_4 = (\widehat{\mu}_{A,1}, \widehat{\boldsymbol{\beta}}^{\mathrm{T}}, \widehat{\phi}, \widehat{\boldsymbol{\kappa}}^{\mathrm{T}}, \widehat{\boldsymbol{\eta}}^{\mathrm{T}})^{\mathrm{T}}$. Fifth, we incorporate $(N_i, N_{i2})$, initialize $\widehat{\psi} = 1$, fix $\widehat{\alpha}_q = 1$, and run BFGS to obtain $\widehat{\boldsymbol{\theta}}_5 = (\widehat{\mu}_{A,1}, \widehat{\boldsymbol{\beta}}^{\mathrm{T}}, \widehat{\phi}, \widehat{\boldsymbol{\kappa}}^{\mathrm{T}}, \widehat{\boldsymbol{\eta}}^{\mathrm{T}}, \widehat{\psi})^{\mathrm{T}}$. Lastly, we optimize over the full parameter set to obtain $\widehat{\boldsymbol{\theta}} = (\widehat{\mu}_{A,1}, \widehat{\boldsymbol{\beta}}^{\mathrm{T}}, \widehat{\phi}, \widehat{\boldsymbol{\kappa}}^{\mathrm{T}}, \widehat{\boldsymbol{\eta}}^{\mathrm{T}}, \widehat{\psi}, \widehat{\boldsymbol{\alpha}}^{\mathrm{T}})^{\mathrm{T}}$, also with BFGS.

This optimization scheme does have inherent challenges to achieve stable convergence. One key regularity condition is that the estimated parameters are not on the boundary of the parameter space, in our case, $\mu_{A,q} > 0$ and $\mu_{B,q} > 0$ corresponding to non-zero expression for each allele and cell type. A second requirement is sufficient variability in $\rho_{iq}$ across samples to estimate $\kappa_q$, $\eta_q$, and $\alpha_q$. This is comparable to an identifiability condition for a linear regression where each covariate has non-zero variance. It is likely that these two requirements are not satisfied for some genes or cell types. Therefore the full model or set of estimable parameters needs to be adjusted. Let $l_n(\boldsymbol{\theta})$, $\dot{l}_n(\boldsymbol{\theta})$, and $\ddot{l}_n(\boldsymbol{\theta})$ denote the log-likelihood, score, and (negative) observed information, respectively. Let $|\cdot|_2$ denote the $L_2$ norm. Convergence is defined when $\left|\dot{l}_n(\widehat{\boldsymbol{\theta}})\right|_2 < \epsilon_1$, $\ddot{l}_n(\boldsymbol{\theta})$ is invertible, no negative variances, and $\left|\ddot{l}_n(\widehat{\boldsymbol{\theta}})^{-1}\dot{l}_n(\widehat{\boldsymbol{\theta}})\right|_2 < \epsilon_2$ for pre-defined thresholds $\epsilon_1$ and $\epsilon_2$. By default, $\epsilon_1 = 10^{-3}$ and $\epsilon_2 = 10^{-6}$. To determine which cell type-specific parameters to constrain to their null values ($\kappa_q = 0, \eta_q = 1, \alpha_q = 1$), we run the above optimization procedure and set the unidentifiable parameters to their null values and re-run the optimization procedure, and iterate this procedure until all the remaining parameters are estimable. More specifically, we initialize our parameters with $\widehat{\boldsymbol{\theta}}_2$ and perform the following operations.

- First, we estimate $\widehat{\boldsymbol{\theta}}_2 = (\widehat{\mu}_{A,1}, \widehat{\boldsymbol{\beta}}^{\mathrm{T}}, \widehat{\phi})^{\mathrm{T}}$ by maximum likelihood estimate (MLE), while ignoring eQTL and cell type composition.

- Next we estimate $\widehat{\boldsymbol{\theta}}_3 = (\widehat{\mu}_{A,1}, \widehat{\boldsymbol{\beta}}^{\mathrm{T}}, \widehat{\phi}, \widehat{\boldsymbol{\kappa}}^{\mathrm{T}})^{\mathrm{T}}$. The $\kappa_q$ parameters are estimated relative to the reference cell type ($q = 1$) and therefore we must ensure the reference cell type has non-zero TReC ($\widehat{\mu}_{A,1} > 0$). By default we set the reference cell type to be the one with highest average proportion across samples. After estimating $\widehat{\boldsymbol{\kappa}}$, we can determine which cell type has highest TReC and swap that cell type to be the reference cell type. This choice of reference cell type can vary from gene to gene. It is an internal choice for the computation purpose and in the final output, all the parameters are transformed using the cell type with highest average proportion as reference. If $\boldsymbol{\theta}_3$ cannot be reliably estimated, it indicates that some $\kappa_q$'s are close to 0. We calculate $\widehat{\mu}_{AA,q} \equiv 2\widehat{\mu}_{A,1}\widehat{\kappa}_q$. If $\min_q(\widehat{\mu}_{AA,q}) < 2$ (each haplotype expresses at least one TReC), set $\widehat{\kappa}_q = 0$ and $\widehat{\eta}_q = \widehat{\alpha}_q = 1$, and then re-optimize.
- Next we estimate eQTL effects $\boldsymbol{\eta}$ in $\boldsymbol{\theta}_4$ and $\boldsymbol{\theta}_5$. If convergence is achieved, move on to the next step. Otherwise, the ASReCs of one or more cell types are near zero. Then we calculate $\widehat{\mu}_{Aq} \equiv \widehat{\mu}_{A,1}\widehat{\kappa}_q$, $\widehat{\mu}_{Bq} \equiv \widehat{\mu}_{Aq}\widehat{\eta}_q$, and $\widehat{\mu}_{zq} \equiv \min(\widehat{\mu}_{Aq}, \widehat{\mu}_{Bq})$, and variance estimate for $\log(\widehat{\eta}_q)$ (optimizing over unconstrained parameters). If $0 < \widehat{\mu}_{zq} < 1$ or $\eta_q$ variance estimate is negative, set $\widehat{\eta}_q = 1$ and re-optimize and repeat until convergence. For each cell type where $\widehat{\eta}_q = 1$, set $\widehat{\alpha}_q = 1$ for subsequent steps.
- Next we estimate $\boldsymbol{\alpha}$ in $\boldsymbol{\theta}_6$. If convergence is achieved, we have established the full model. Otherwise, check for variance estimates for $\log(\widehat{\alpha}_q)$ that are negative and set $\widehat{\alpha}_q = 1$ and re-optimize. If none of the cell types $\alpha_q$ variances are negative and convergence is not achieved, identify the cell type with largest $\alpha_q$ variance and set $\widehat{\alpha}_q = 1$.

In general, whenever we need to re-optimize, we simply start off at the step prior to the current step, there is no need to return to $\widehat{\boldsymbol{\theta}}_2$ since the procedure has established the "submodel" or nested set of estimated parameters that achieved convergence. In addition, our procedure is strictly designed to assess convergence first at each step before attempting to constrain parameter estimates or looking to the variance estimates to avoid unnecessary matrix inversions until all other criteria are met.

### 3 Trimming Influential Counts

Data trimming and quality control are a crucial issue associated with regression analyses [6]. Modeling observed outcomes directly risks highly influential or potential outlier data points that contribute to biased parameter estimates and inflated type I error. For eQTL analysis, if the same subset of samples were consistently identified as outlier, we could exclude them. But among post quality control samples, an analysis could involve potentially excluding different subsets of samples per gene, risking a power loss and difficulty to interpret the results. In the case of differential expression, DESeq2 [7] systematically trims outlier observations whose Cook's distance is beyond a predefined cutoff based on the F-distribution. We have adopted a similar trimming approach in our eQTL analyses.

For a given gene, we characterize the influence of a sample through our definition of Cook's Distance for the $i$th sample with

$$C_i = \frac{1}{m}\sum_{j=1}^{n}\frac{(\widehat{\mu}_{j(i)} - \widehat{\mu}_j)^2}{\widehat{v}_j}$$

where $m = p + Q - 1$, $\widehat{\mu}_j$ is the estimated mean TReC for the $j$th sample, $\widehat{\mu}_{j(i)}$ is the estimated mean TReC for the $j$th sample after excluding the $i$th sample, and $\widehat{v}_j = \widehat{\mu}_j + \widehat{\phi}\widehat{\mu}_j^2$, the estimated TReC variance for the $j$th sample. Since our TReC model is not the traditional GLM due to the sample-specific offset term $\left(\log\left(\sum_{q=1}^{Q}\rho_{iq}\kappa_q\right)\right)$, we cannot directly characterize leverage. We then calculate normalized Cook's distance to put Cook's distances on the same scale across genes, denoted $\tilde{C}_i$ and characterized as

$$\tilde{C}_i = \frac{C_i - \mathtt{med}(C_1, \ldots, C_n)}{\mathtt{mad}(C_1, \ldots, C_n)},$$

where $\mathtt{med}(\ldots)$ and $\mathtt{mad}(\ldots)$ denote median and median absolute deviation, respectively. We propose trimming the original TReC ($T_i$) if $\tilde{C}_i > c$ where $c$ is some predefined threshold. To calculate Cook's distance, we fit CSeQTL's TReC model without adjusting for SNP since a gene can have multiple SNPs and a gene's TReC can be influential regardless of genotype. We explored the possibility of using $C_i > 4/n$ and $C_i > F(q = 0.99, m, n-m)$ as a trimming criteria however it failed to detect clear visual outliers. We decided on an appropriate threshold on $\tilde{C}_i$ by running CSeQTL on chr1 genes with permuted SNP genotypes. We tried cutoff thresholds 40, 20, 10, and 5. The largest threshold that controls the type I error was selected. Unlike the trimmed means used by DESeq2 to impute the TReC value, we impute the TReC value with the estimated TReC for a sample from CSeQTL's TReC model without SNP adjustment.

### 4 Simulation Setup

We describe how the cell type proportions are simulated. In the first scenario, let $X \sim U(a, b)$ denote a random variable $X$ sampled from a continuous uniform distribution ranging from $a$ to $b$. Specifically $\rho_{iq} = \exp\{U_{iq}\} / \sum_{s=1}^{Q} \exp\{U_{is}\}$ and $U_{iq} \sim U(-4, 4)$. In the second scenario, to allow cell types to reflect observed proportions with wide and narrow ranges of proportions, we simulated $\rho_{i1}$ from a beta distribution with shape parameters 10 and 24 (values derived based on maximum likelihood estimates from fitting a beta distribution to CMC's astrocyte cell type proportions), $\rho_{i2} = \left|0.85 - 0.76\rho_{i1} - 0.03\rho_{i1}^2 + \epsilon_i\right|$, where $\epsilon_i$ was sampled from a centered normal distribution with standard deviation 0.02, and $\rho_{i3} = 1 - \rho_{i1} - \rho_{i2}$. If $\rho_{i3} < 0$, we set it to zero and normalize the proportions across cell types. For the third scenario, proportions are first simulated under the second scenario. Next, for each cell type, the initial proportions greater than the 99% quantile were replaced by values sampled from $U(0.7, 0.9)$ while initial proportions less than the 1% quantile were replaced by values sampled from $U(0, 0.1)$. These final values are re-normalized across cell types to sum to one.

For $n = 300$, we simulate $p = 4$ baseline covariates. The first covariate is $X_{i1}$, which represents read-depth, is simulated by a gamma distribution with shape parameter set to 600 and rate parameter set to 100, based on empirical MLE estimates from CMC samples. $X_{i2}$, which represent sex, is generated by a Bernoulli distribution with success probability of 0.5. $X_{i3}$ is generated by a continuous uniform distribution ranging from $-1$ to 1. $X_{i4}$ is simulated by a standard normal distribution. These latter two variables represent arbitrarily distributed continuous covariates. Continuous covariates $X_{i1}$, $X_{i3}$, and $X_{i4}$ are centered and scaled with zero mean and unit variance. Assuming Hardy Weinberg equilibrium, genotypes were generated using a categorical distribution with probabilities $(1 - m_A)^2$, $m_A(1 - m_A)$, $m_A(1 - m_A)$, $m_A^2$ for $AA$, $AB$, $BA$, $BB$, respectively, where $m_A$ denotes the minor allele frequency. We set $m_A = 0.2$.

### 5 Grouping 22 blood cell types to 7 cell types

The "CD4T" cell type is defined by pooling CD4 naive, CD4 memory resting, CD4 memory activated, follicular helper, regulatory T cells (Tregs) and gamma delta cells. The gamma delta T cells is indeed a different type of T cells while all other type of T cells are alpha beta T cells. However, its proportion is very low (Supplementary Figure 8) and thus adding it to any other cell type does not lead to any noticeable change of cell type composition. Here we combine it into the CD4T category just for the convenience of implementation. The "B_Cell" cell type is the result of combining B cell naive, B cell memory and Plasma cells. CD8 T cells were not collapsed with other cell types and simply denoted "CD8T". The "Mast_Eosinophil" cell type is composed of mast cells resting, mast cells activated, dendritic cells resting, dendritic cells activated and eosinophils. The "NK" cell type comprises of natural killer cells resting and natural killer cells activated. The "Monocytes" cell type is made up of monocytes, macrophages M0, macrophages M1 and macrophages M2. The proportion of microphage cells are very low (Supplementary Figure 8) and thus adding them to monocytes does not make substantial changes to monocyte proportions. We further discuss the algebraic interpretation and implications of combining cell types in Supplementary Materials 2.5.
