## Supplementary Materials for "A computational method for Cell type-specific Expression Quantitative Trait Loci mapping using bulk RNA-seq data"

**3 Multiple Testing and FDR** **12**

**4 Data Preprocessing** **13**

**5 Quality Control, Filtering, ASReC adjustment** **14**

**6 Real Data Analyses** **17**

### 1 Extended Simulations

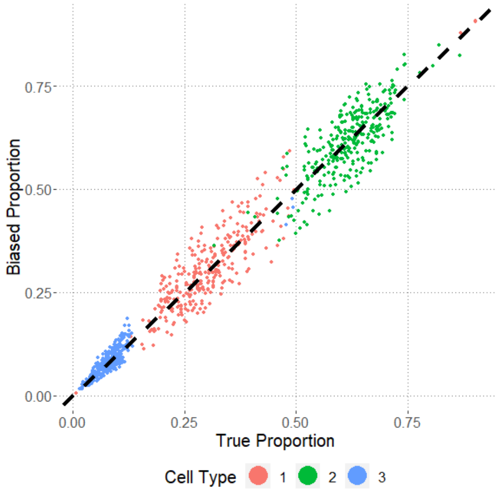

Figure 1: Simulated cell type proportions under Scenario 3 with cell type proportions after adding noise on the y-axis and true proportions on the x-axis.

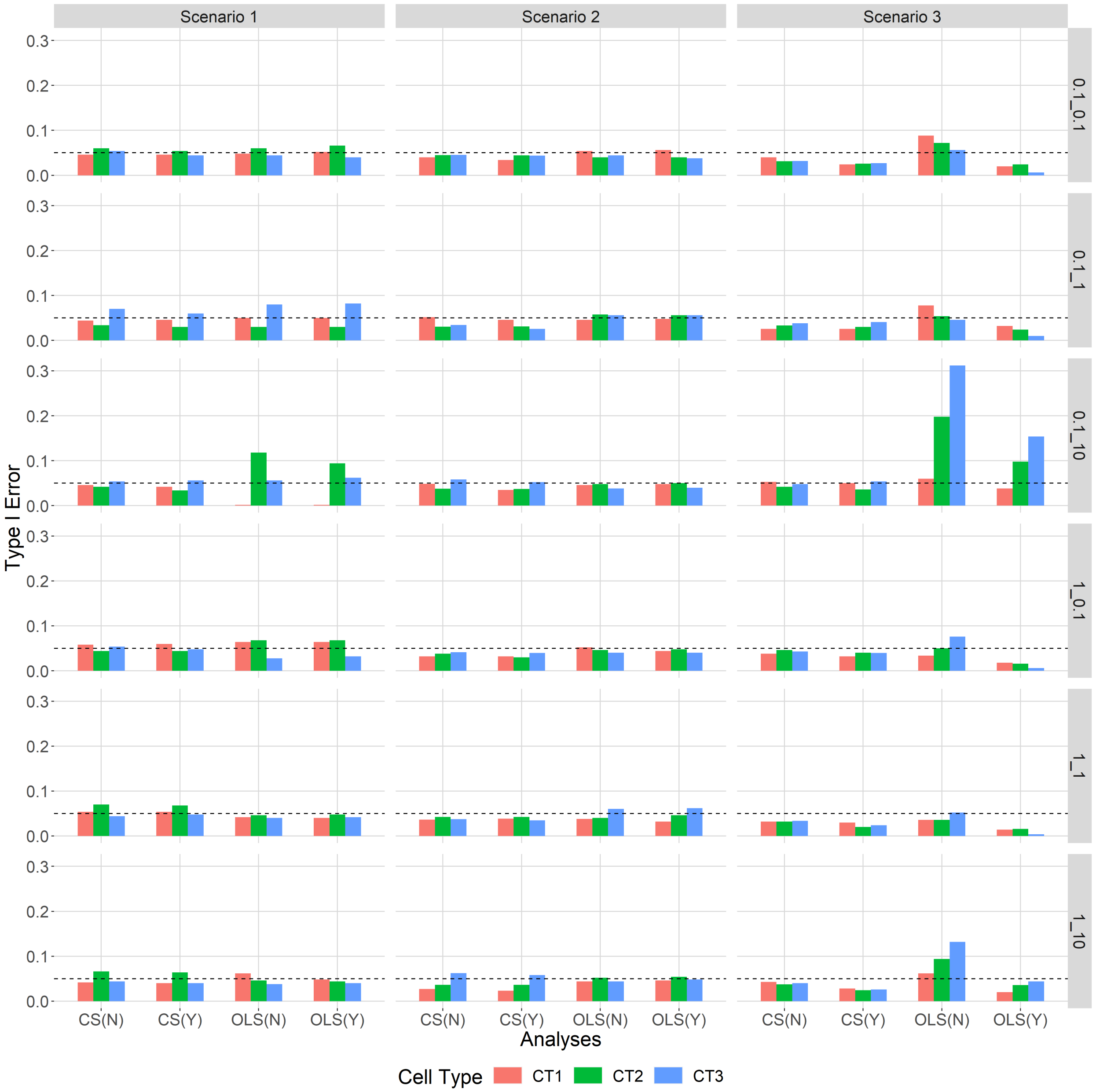

Figure 2: Type I error for the data simulated under complete null, with **true** cell type proportions. X-axis labels CS(N), CS(Y), OLS(N), and OLS(Y) correspond to CSeQTL w/o trimming, CSeQTL w/ trimming, OLS w/o trimming, and OLS w/ trimming, respectively.

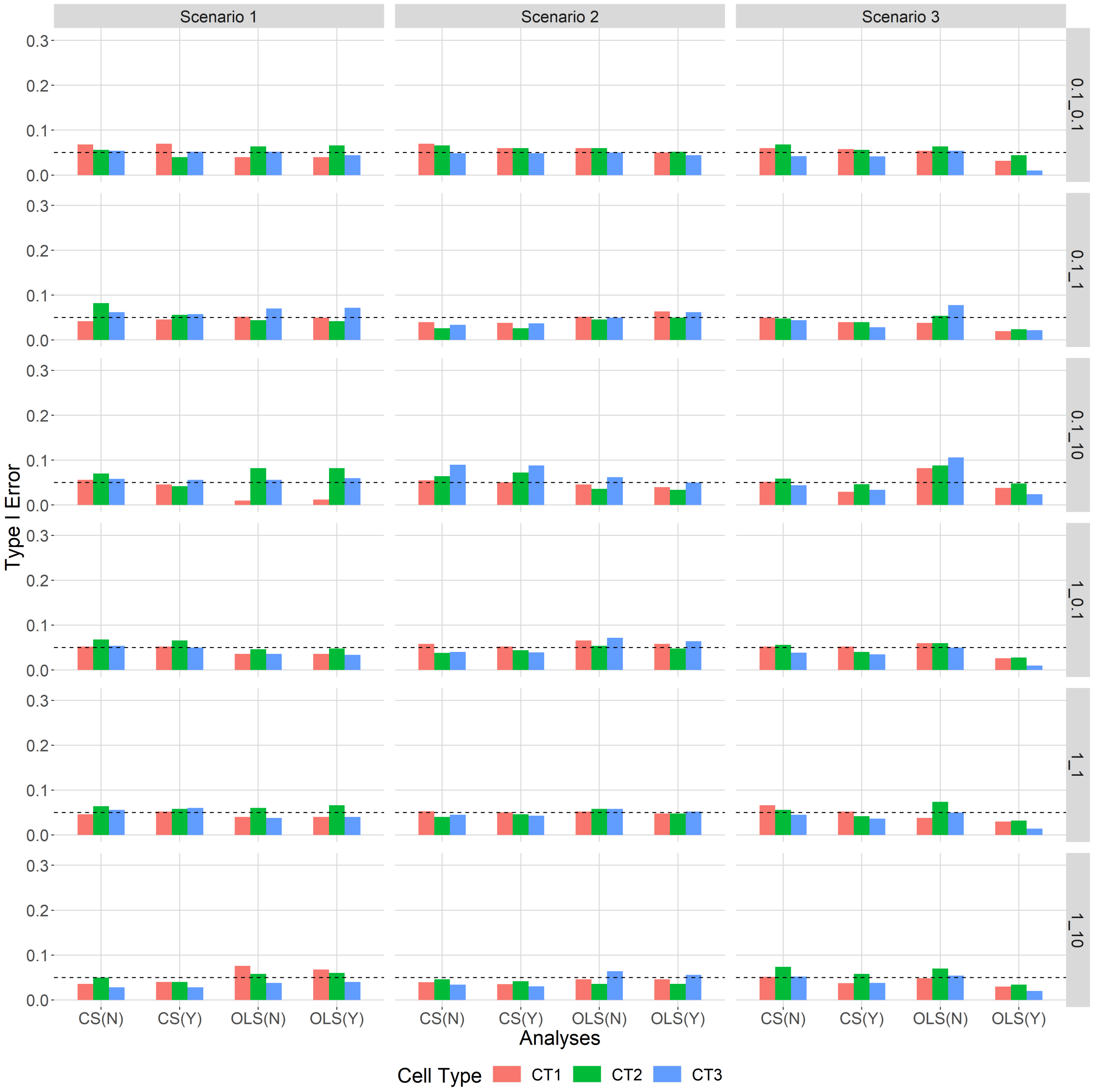

Figure 3: Type I error for the data simulated under complete null, with **noisy** cell type proportions. X-axis labels CS(N), CS(Y), OLS(N), and OLS(Y) correspond to CSeQTL w/o trimming, CSeQTL w/ trimming, OLS w/o trimming, and OLS w/ trimming, respectively.

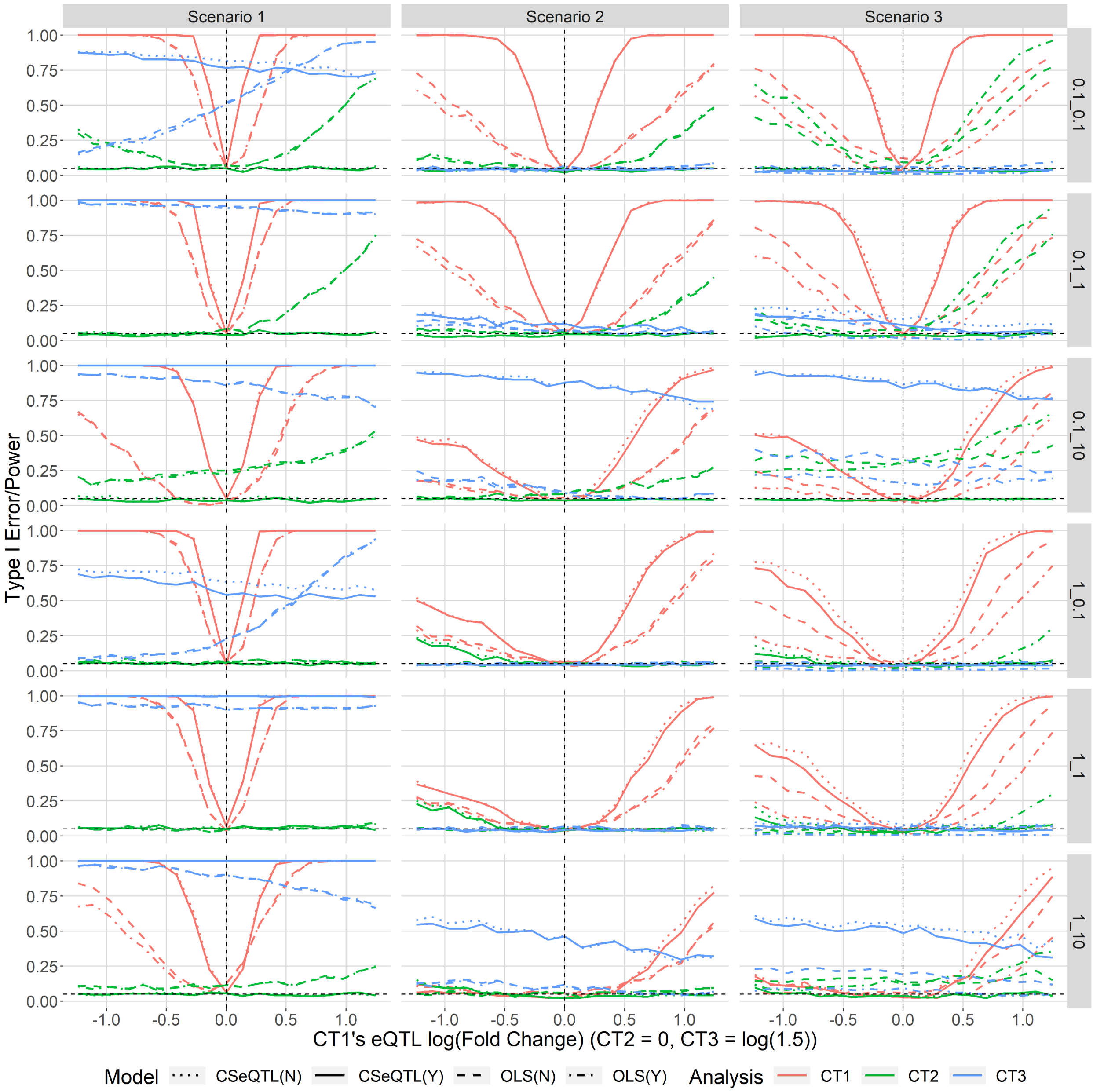

Figure 4: Power/Type I error for the data simulated under mixed null/alternative with **true** cell type proportions. X-axis labels CSeQTL(N), CSeQTL(Y), OLS(N), and OLS(Y) correspond to CSeQTL w/o trimming, CSeQTL w/ trimming, OLS w/o trimming, and OLS w/ trimming, respectively.

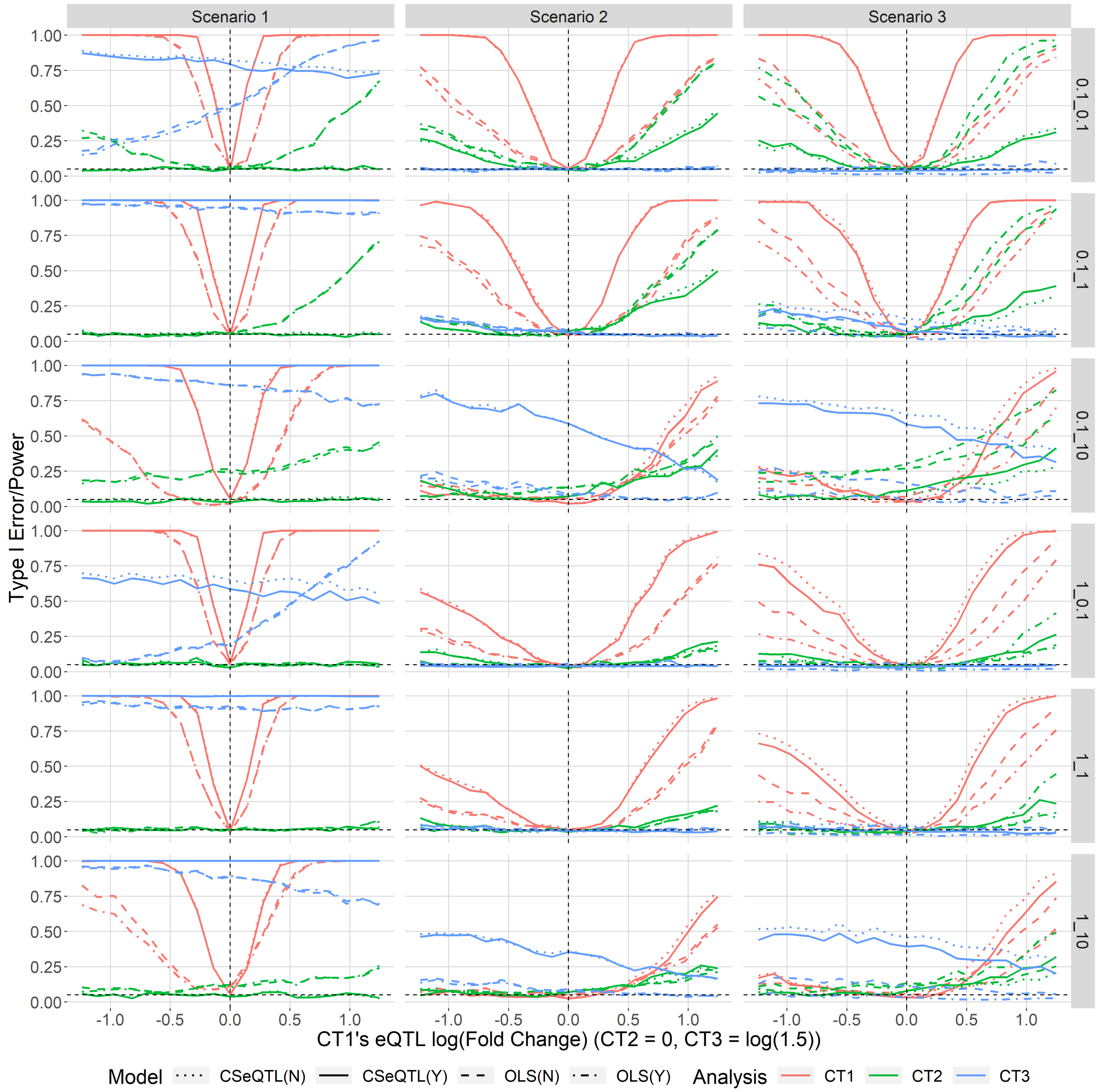

Figure 5: Power/Type I error for the data simulated under mixed null/alternative with **noisy** cell type proportions. X-axis labels CSeQTL(N), CSeQTL(Y), OLS(N), and OLS(Y) correspond to CSeQTL w/o trimming, CSeQTL w/ trimming, OLS w/o trimming, and OLS w/ trimming, respectively.

### 2 Deconvolution

#### 2.1 Input/output for deconvolution

To perform deconvolution, total read count data from the $i$-th bulk RNA-seq sample ($T_{ig}$) and reference scRNA-seq data for the $q$th cell type, denoted by $R_{qg}$, across $G$ shared genes are converted to transcripts per million (TPM), denoted by $t_{ig}$ and $r_{qg}$:

$$t_{ig} = 10^6 \frac{T_{ig}/l_g}{\sum_{h=1}^{G} T_{ih}/l_h} \text{ and } r_{qg} = 10^6 \frac{R_{qg}/l_g}{\sum_{h=1}^{G} R_{qh}/l_h},$$

where $l_g$ denotes the $g$th gene's exonic region length. For reference data from scRNA-seq, we can calculate cell size factors for the $q$th cell type, denoted $s_q$ and it is proportional to $\sum_{g=1}^{G}(R_{qg}/l_g)$. For ease of comparison across other studies, we calculate $s_q = \sum_{g=1}^{G}(R_{qg}/l_g)$ and then normalize it by the largest cell type's cell size ($s_q = s_q/\max_r(s_r)$). For reference data not derived from scRNA-seq, we use cell size factors reported by other studies. We estimate cell type proportions using ICeDT [1], with TPM data as input, setting tumor purity to 0 for all samples, and specifying default values for the remaining arguments. Cell type proportion estimates per sample, denoted by $\tilde{\rho}_{iq}$, are considered proportions of transcripts per cell type that are then corrected using cell size factors, such that the final cell type proportion estimates are given by

$$\rho_{iq} = \frac{\tilde{\rho}_{iq}/s_q}{\sum_{r=1}^{Q} \tilde{\rho}_{ir}/s_r}.$$

For CMC and GTEx brain cohorts, we corrected for cell size factors from MTG (astrocyte = 0.885, excitatory = 0.893, inhibitory = 1.00, microglia = 0.856, oligodendrocyte = 0.927, oligodendrocyte precursor cells = 0.726). For whole blood samples we used cell sizes reported by EPIC [2], which have been utilized in our earlier work [1].

#### 2.2 Considered Methods

Many methods have been developed to estimate cell type compositions and most of them rely on cell type-specific expression from external sources, such as scRNA-seq or purified samples of individual cell types [3, 4]. These methods differ in their models and strategies to improve their robustness and accuracy. For example, CIBERSORT [4] employs support-vector regression and its relu loss function is robust to outliers. TIMER [5] uses a linear regression with least squared loss function, and thus is sensitive to outliers. Their strategy is to remove the genes with very high expression due to their strong influence on model fitting. EPIC [2] uses weighted linear regression to give the genes with lower expression variation higher weights. ICeDT [1], which shares some similar spirit with an earlier method DeMix [6], models the gene expression by a log-normal distribution to reduce the impact of genes with high expression and conducts deconvolution in the linear scale. Importantly, ICeDT acknowledges the fact that for some genes, cell type-specific expression from external sources may not be consistent with their expression in bulk tissue samples. ICeDT uses a mixture model to down-weigh the contribution of such genes in cell type composition estimation.

#### 2.3 LM22 transformation

For whole blood, the existing LM22 dataset, derived from microarray data would be utilized and transformed for deconvolution with RNA-seq based expression. LM22 was obtained through the CIBERSORT website and it was a matrix of gene expression for 547 marker genes and 22 cell types. CIBERSORT performs deconvolution on centered and scaled bulk and signature gene expression, this eliminates concern on units or scaling between bulk and signature expression. However, for ICeDT deconvolution, we need to ensure that the bulk and signature expression were on the same scale. To achieve this, we transformed LM22 signatures based on an assumption that bulk TPM from samples with very high proportions of neutrophils should strongly correlate with neutrophil expression in the LM22 matrix. We first estimate cell type proportions by CIBERSORT, with TPM-transformed bulk expression and untransformed LM22 as input. Using these estimates, we subsetted samples with estimated neutrophil proportions greater than 85%, which resulted in 80 samples. We identified a subset of neutrophil-specific marker genes who have highest expression in neutrophil and their expression in the next cell type is less than half of their expression in neutrophil. For each of the 80 samples with high neutrophil proportion, we fitted a linear model per sample (log bulk TPM vs. log neutrophil expression from LM22) using those neutrophil-specific marker genes, and retained sample-specific slopes and intercepts (Supplementary Figure 6). The average slope (1.031) and intercept (-2.32) were used to transform all the genes in LM22 matrix by multiplying the log-transformed LM22 matrix by 1.031, subtracting 2.32, and exponentiating. We assumed the transformation may leave some gene expression incompatible between bulk and signature. We then compared the updated LM22 signature with the bulk TPM for each gene. More specifically, for each gene, we calculated the proportion of samples whose bulk expression was less than the minimum cell type-specific expression, as well as the proportion of samples whose bulk expression was greater than the maximum cell type-specific expression. Using cutoffs of 15% for both quantities, we subsetted 228 genes for ICeDT deconvolution (Supplementary Figure 7).

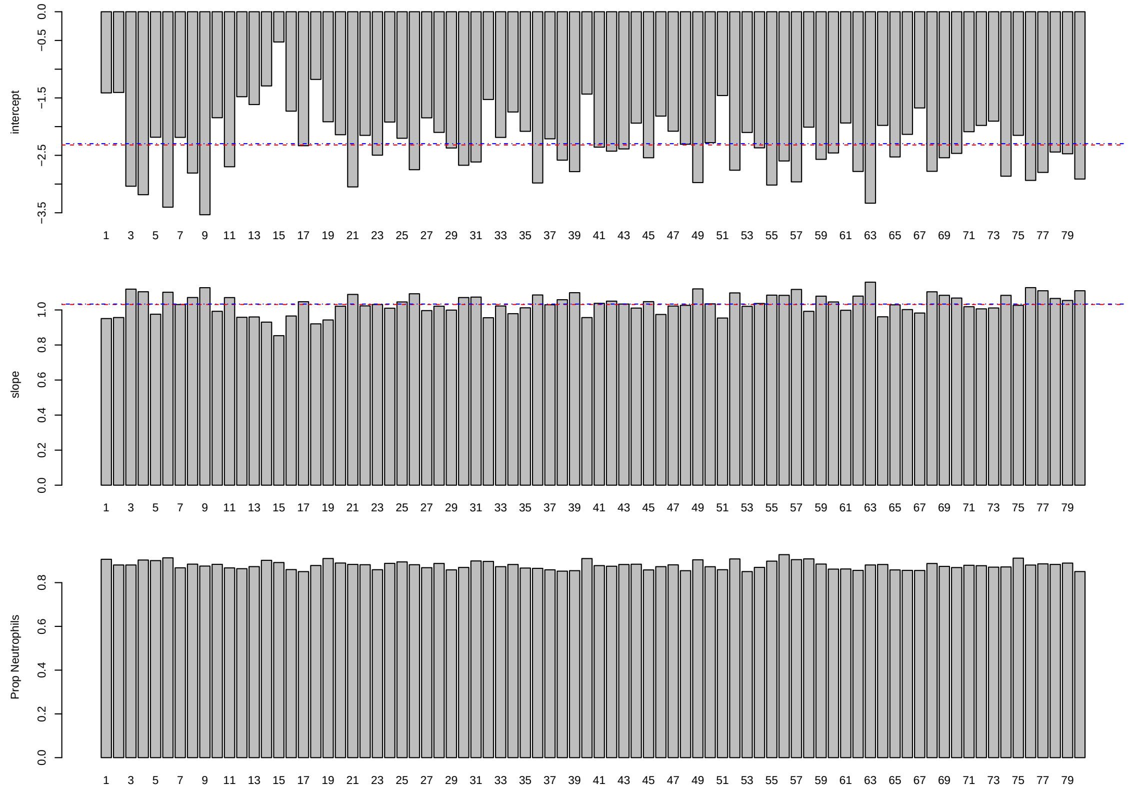

Figure 6: **GTEx whole blood samples with greater than 85% neutrophil composition.** The first, second, and third barplots correspond to linear fit intercepts and slopes and neutrophil proportions, respectively.

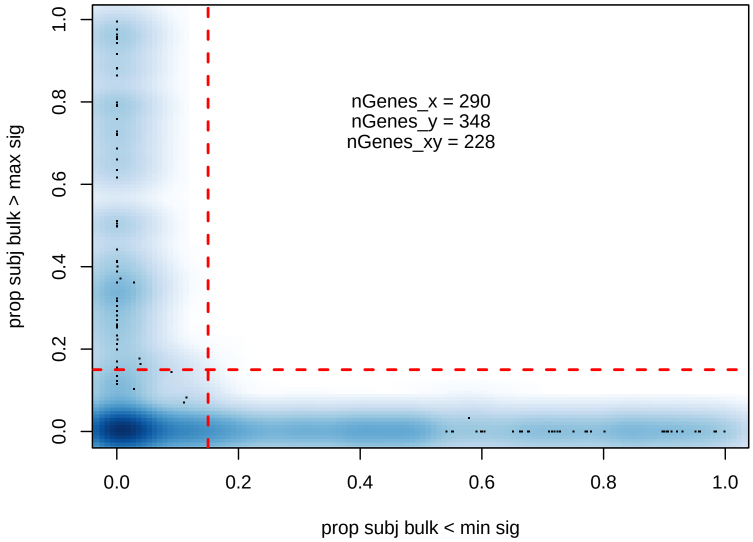

Figure 7: **Selection of marker genes for cell types in blood.** Each point is a marker gene. The x-axis is the proportion of bulk samples where this gene's expression is smaller than the minimum expression of this gene across all cell types. The y-axis is the proportion of bulk samples where this gene's expression is larger than the maximum expression of this gene across all cell types.

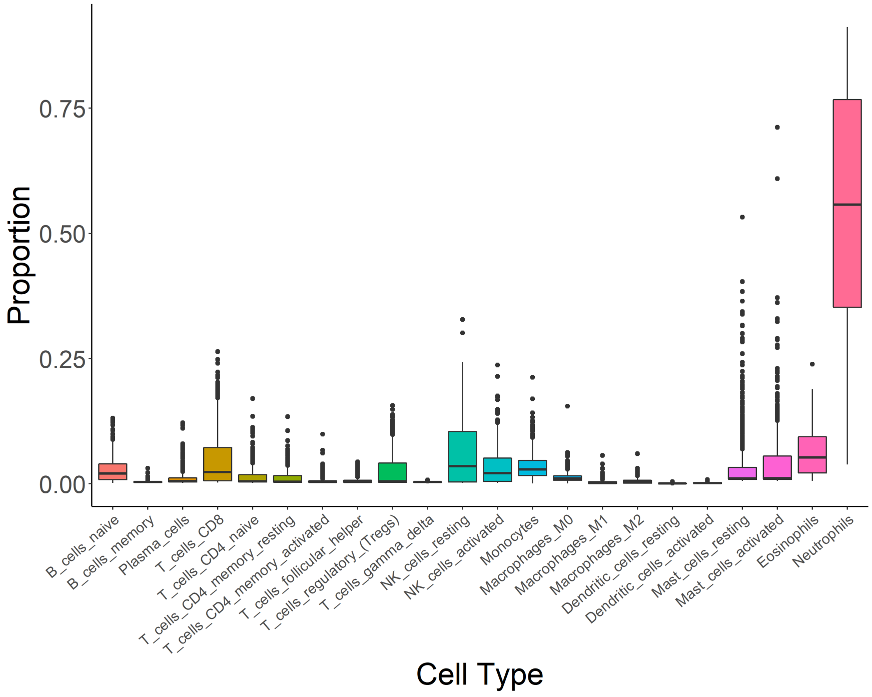

Figure 8: **ICeDT deconvolution estimates of cell type proportion for GTEx whole blood samples.**

#### 2.4 Signature Expression from scRNA-seq

To perform deconvolution on GTEx's brain frontal cortex and CMC's dorsolateral prefrontal cortex (LDPFC) samples, we studied three single nucleus RNA-seq (snRNA-seq) datasets: DroNc-seq [7] (19,550 nuclei from human brain), the Allen Brain Atlas's middle temporal gyrus (MTG) datasets [8] (15,928 nuclei by SMART-seq platform), and a very recent dataset from The Seattle Alzheimer's Disease Brain Cell Atlas (SEA-AD), which included more than 1 million nuclei from human middle temporal gyrus (MTG) of 89 donors, using 10x 3' platform. After running a workflow of quality control and clustering, we concluded not to use the DroNc-seq dataset since its clustering results were unstable, likely due to high sparsity and limited number of nuclei. In contrast, clustering results by MTG data were more robust. Different methods led to similar clusters that matched pre-labeled cell types. This was likely because MTG used the SMART-seq protocol, which generated less sparse and full-length gene expression data.

We processed MTG data using our workflow `www.github.com/Sun-lab/scRNAseq_pipelines/MTG`, which utilized R packages scater [9], scran [10], limma [11], and Rtsne [12]. When processing MTG data, 15,929 nuclei underwent quality control (QC) and filtering. We first filtered out nuclei with less than 50% unique reads, and 15,929 nuclei were retained. Next, genes were subjected to QC and filtering by retaining genes that were detected (at least 1 gene count) in at least 30 nuclei, which left us 37,657 genes in the following analysis. Normalization was conducted by calculating size factors using functions quickCluster() and computeSumFactors(). We identified a strong positive correlation between computed size factors and total counts. We used makeTechTrend() to decompose the variance of gene expression into biological and technical noise. A total of 4,745 highly variable genes were selected based on an FDR threshold of $10^{-20}$ and a bio threshold of 0.1, and they were used as inputs for PCA. We selected the top 50 PCs for tSNE visualization and kMeans clustering. We ran kMeans clustering with fifty initializations with specified number of clusters ranging from 10 to 20, each time comparing the clustering results to the pre-labeled cell type clusters generated by MTG. At 15 clusters, cell types and clusters had sufficient one-to-one match with each other. Among the pre-labeled cell types, only nine nuclei were labeled as endothelial and our clusters contained endothelial nuclei that overlapped strongly with OPC cell types. Thus we excluded endothelial cell type from our final reference expression calculation. We next used MAST [13] to identify approximately 120 top marker genes per cell type based on fold change and q-value. Once marker genes were identified, scRNA-seq counts were averaged within cell type to construct our signature expression matrix for subsequent bulk RNA-seq deconvolution.

We also compared gene expression similarity as well as cell type composition estimation when using MTG vs. SEA-AD data. Our results showed that the expression of signature genes were highly correlated between the two datasets and they led to very consistent cell type composition estimation (Figure 9). Therefore, the large number of nuclei available from the SEA-AD study likely overcame the sparsity limitation.

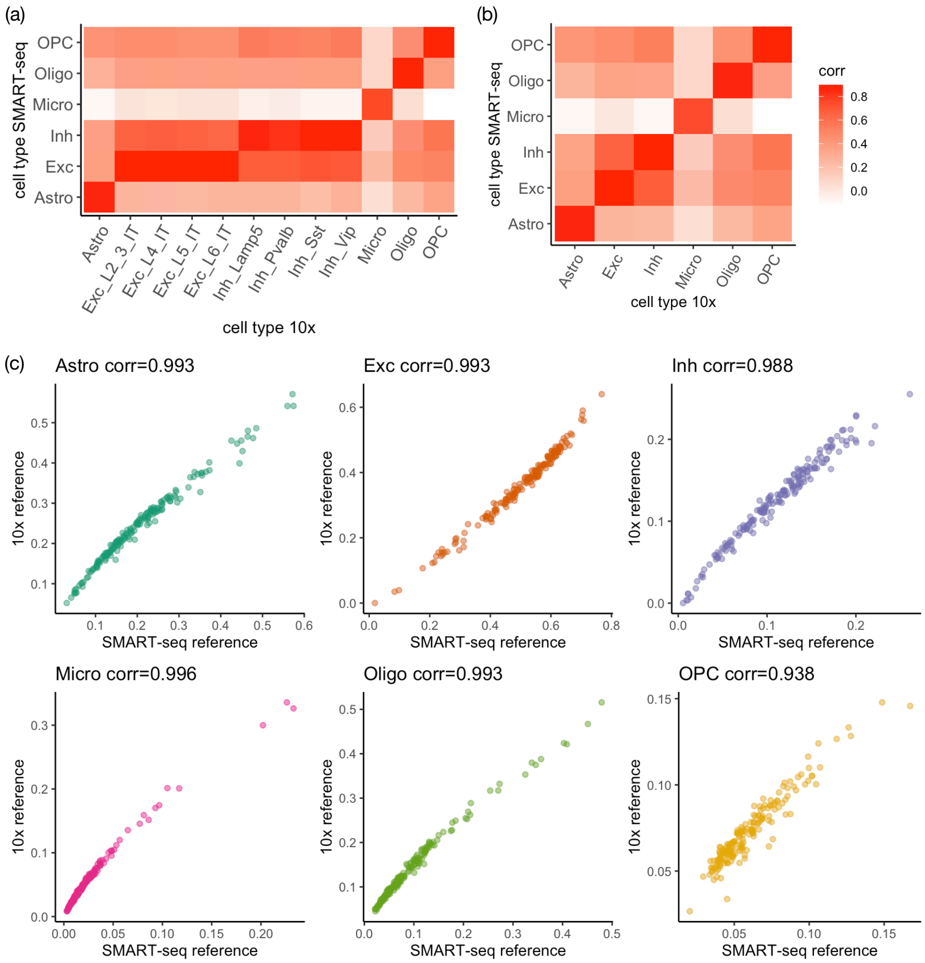

Figure 9: **Compare cell type reference by SMART-seq vs. 10x 3' platform.** (a) Correlation of gene expression of 6 cell types from SMART-seq data versus gene expression of 12 cell types (4 sub-types of excitatory neurons labeled by Exc_L*, 4 subtypes of inhibitory neurons labeled by Inh_*, and 4 type of glia cells. (b) Similar correlation matrix as (a) except that excitatory/inhibitory neuron subtypes are combined (by adding up expression from all the cells) to excitatory/inhibitory neurons. (c). Scatterplots of cell type proportion estimates based on SMART-seq or 10x references.

#### 2.5 Collapsing cell types

We provide an algebraic derivation of the loss of explained variability in TReC resulting from collapsing cell types. Suppose we have three cell types ($Q = 3$) and we would like to collapse cell types 1 and 2 together. We would like to interpret the parameters of the collapsed model with regard to the original parameters. Starting with the definition of the $i$th sample's bulk eQTL, we have

$$\xi_i = \frac{\mu_{iB}}{\mu_{iA}} = \frac{\sum_q \rho_{iq}\kappa_q\eta_q}{\sum_q \rho_{iq}\kappa_q} = \frac{\rho_{i1}\eta_1 + \rho_{i2}\kappa_2\eta_2 + \rho_{i3}\kappa_3\eta_3}{\rho_{i1} + \rho_{i2}\kappa_2 + \rho_{i3}\kappa_3} = \frac{(\rho_{i1} + \rho_{i2})\eta_{i12} + \rho_{i3}\kappa_{i3}\eta_3}{(\rho_{i1} + \rho_{i2}) + \rho_{i3}\kappa_{i3}}$$

where

$$\eta_{i12} = \frac{\rho_{i1}\eta_1 + \rho_{i2}\kappa_2\eta_2}{\rho_{i1} + \rho_{i2}\kappa_2} = \frac{\rho_{i1}}{\rho_{i1} + \kappa_2\rho_{i2}}\eta_1 + \left(1 - \frac{\rho_{i1}}{\rho_{i1} + \kappa_2\rho_{i2}}\right)\eta_2 \text{ and } \kappa_{i3} = \frac{\rho_{i1} + \rho_{i2}}{\rho_{i1} + \kappa_2\rho_{i2}}\kappa_3.$$

Thus the combined cell type eQTL parameter is now weighted between the first two cell type eQTLs where the weights are functions of cell type proportion and $\kappa_2$. Also, for the TReC model's offset term, we have

$$\begin{aligned}
\log\left(\sum_q \rho_{iq}\kappa_q\right) &= \log\left(\rho_{i1} + \rho_{i2}\kappa_2 + \rho_{i3}\kappa_3\right) = \log\left(\frac{\rho_{i1} + \rho_{i2}\kappa_2}{\rho_{i1} + \rho_{i2}}(\rho_{i1} + \rho_{i2}) + \rho_{i3}\kappa_3\right) \\
&= \log\left(\frac{\rho_{i1} + \rho_{i2}\kappa_2}{\rho_{i1} + \rho_{i2}}\left[\rho_{i1} + \rho_{i2} + \frac{\rho_{i1} + \rho_{i2}}{\rho_{i1} + \rho_{i2}\kappa_2}\rho_{i3}\kappa_3\right]\right) \\
&= \log\left(\frac{\rho_{i1} + \rho_{i2}\kappa_2}{\rho_{i1} + \rho_{i2}}\right) + \log\left(\rho_{i1} + \rho_{i2} + \rho_{i3}\kappa_{i3}\right)
\end{aligned}$$

Thus, if we collapse a subset of cell types and proceed with optimization, this assumes that these sample-specific terms $\eta_{i12}$, $\kappa_{i3}$, and $\log\left(\frac{\rho_{i1} + \rho_{i2}\kappa_2}{\rho_{i1} + \rho_{i2}}\right)$ are constant parameters for all $i$. These sample-specific terms can potentially explain a source of variability in TReC that is lost through collapsing cell types. Given our framework, this may lead to Type I Error inflation which we can observe through SNP permutation analyses. However, if a subset of cell types, denoted $Q_s$, present at very similar proportions across all samples (i.e. $\rho_{iq} \approx \rho_{jq}$ for samples $i, j$ and $q \in Q_s$), then collapsing cell types can be justified and recommended since cell types with narrow spread in proportion can lead to unstable $\kappa_q$, $\eta_q$, and $\alpha_q$ estimates.

### 3 Multiple Testing and FDR

Just as in existing eQTL mapping methods, the issue of multiple testing arises for a gene's tested SNPs. The linkage disequilibrium (LD) of nearby SNPs leads to redundancy and so, similar to Sun (2012) [14] and Hu et al. [15], we propose selecting the most significant SNP per gene and cell type after accounting for the multiple testing. But unlike the aforementioned approaches, CSeQTL does not scale well with using SNP permutation to approximate a gene's minimum p-value null distribution from the added computation of cell type-specific eQTLs and reference expression fold changes. For the $g$th gene, suppose there are $S_g$ corresponding SNPs. If $S_g$ tests along a gene were independent one could simply apply Bonferroni adjustment to the eQTL significance testing p-values. However due to LD, we risk a loss in power. Instead, we chose to employ the geoP approach [16] for approximating the number of independent tests among $S_g$ SNPs to approximate the permutation p-value. Let $\tilde{S}_g$ denote the number of independent tests from geoP and let $p_{gs}$ denote the $s$th SNP's nominal p-value tested for a specific analysis (bulk or cell type-specific). Let $p_g$ denote the $g$th gene's minimum p-value SNP approximated permutation p-value and defined as $p_g = \min\left(1, \min_s(p_{gs})\tilde{S}_g\right)$.

With regard to assessing the number of genes with eQTLs, we control for the false discovery rate (FDR) using the approach by Storey and Tibshirani [17] with a small modification. This is due to the observation that many permutation p-values equal 1.0. First, FDR is defined as

$$\text{FDR}(p_c) = \frac{\pi_0 p_c N_G}{\sum_{g=1}^{G} 1\left\{p_g \leq p_c\right\}},$$

where $\pi_0$ is the expected proportion of true null hypotheses, $p_c$ is the significance p-value cutoff, $1\{A\} = 1$ if $A$ is true and $1\{A\} = 0$ otherwise, and $N_G$ is the number of genes tested. Indexed by $\lambda$, the parameter $\pi_0$ is typically estimated as

$$\widehat{\pi}_0(\lambda) = \frac{1}{N_G} \sum_{g=1}^{G} \frac{1\{p_g \geq \lambda\}}{1-\lambda},$$

where $0 < \lambda < 1$. Our modification assumes the observed $p_g$'s equal to 1.0 are actually continuous uniformly distributed between 0.0 and 1.0 and thus we have

$$\widehat{\pi}_0(\lambda) = \frac{1}{N_G} \sum_{g=1}^{G} \left[ \frac{1\{1 > p_g \geq \lambda\}}{1-\lambda} + 1\{p_g = 1\} \right],$$

where we chose to set $\lambda = 0.5$. We can search over a grid of values $p_c$ to achieve the desired FDR (such as 0.01, 0.05, 0.10). The q-value is then calculated as

$$\text{q-value}(p_g) = \min_{p_c \geq p_g} \text{FDR}(p_c).$$

### 4 Data Preprocessing

We assessed data collected from the CommonMind Consortium (CMC) [18, 19], BLUEPRINT [20] for its purified blood cell types, Genotype Tissue Expression's (GTEx) Whole Blood (WB) and Brain Frontal Cortex (BFC).

Among CMC samples, we obtained 606 RNA-seq bam files and corresponding genotypes, from SNP6 microarray, on control and schizophrenic (SCZ) samples using the Synapse platform. For CMC, starting with 752,269 genotyped SNPs by Illumina Infinium HumanOmniExpressExome 8v1.1b chip, we arrived at 5,792,709 imputed and phased SNPs across autosomes using 1,000 Genomes Phase 3 reference panels after phasing with SHAPEIT v2.837 [21], imputation of variants in 5 Mb segments with IMPUTE2 v2.3.2 [22], retaining loci with info metric >0.3, minor allele frequency >0.05, and SNPs with allelic dosages >0.8, and lifting coordinates from GRCh37 to GRCh38.

For the BLUEPRINT cohort, we used the European Genome Archive's pyega3 client to download phased SNPs derived from whole genome sequencing (EGAD00001002663) and three purified cell types of RNA-seq bam files (EGAD00001002671, EGAD00001002674, EGAD00001002675) corresponding to CD4+ alpha-beta T cell (n=212), CD14+/CD16- monocytes (n=197), and mature neutrophil (n=205), respectively. BLUEPRINT's VCF contained phased GRCh37 loci from whole genome sequencing that we lifted over to GRCh38 resulting in 4,851,911 successfully mapped and phased SNPs.

For GTEx WB and BFC cohorts, preprocessed phased variants from whole genome sequencing were downloaded from `https://anvil.terra.bio/#workspaces/anvil-datastorage/AnVIL_GTEx_V8_hg38`. GTEx's VCF provided 5,938,874 phased SNPs mapped to GRCh38. Preprocessed GRCh38-aligned RNA-seq bam files were obtained and processed on Google Cloud to obtain TReC and ASReC per gene through the AnVIL platform, using the pipeline from the GitHub repository, gtex_AnVIL: `https://github.com/Sun-lab/gtex_AnVIL` [16].

We have reprocessed the RNA-seq bam files from CMC and BLUEPRINT following GTEx's pipeline, available at `https://github.com/broadinstitute/gtex-pipeline/tree/master/rnaseq`, and contained in Figure 10. RNA-seq bam files were converted from bam to fastq with Picard v2.21.7 [23]. Samples were aligned using reference genome GRCh38 excluding ALT, HLA, and Decoy contigs but including ERCC spike-ins. Gene models were based on GENCODE's v26 GRCh38 with ERCC spike-in. The STAR index was constructed using the aforementioned reference genome and gene models. Reads were aligned using STAR v2.5.3a with input parameters provided by GTEx's template. Bam files were subsequently sorted by coordinate and indexed with Samtools v1.8 and duplicated reads were marked by Picard. With CMC's imputed SNPs and WGS SNPs for BLUEPRINT, WB, and BFC, reads were next sorted by read name. Then, bam files were passed into asSeq [24] v0.99.501 to obtain TReC and ASReC for haplotype 1 or haplotype 2. In addition, some RNA-seq reads that harbor more than one heterozygous SNPs are not compatible with either haplotype. We refer to those as haplotype-ambiguous reads.

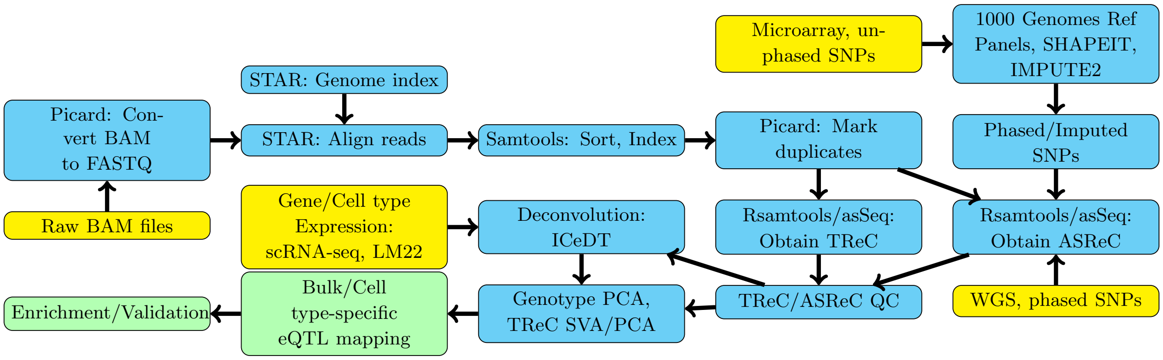

Figure 10: **Analysis Workflow.** Yellow nodes correspond to initial input data files. Blue nodes correspond to intermediate workflow steps. Green nodes correspond to our contributed analyses.

### 5 Quality Control, Filtering, ASReC adjustment

We started by plotting samples' total TReC versus total ASReC summed across genes to identify potential outliers. The exact threshold varied across studies as described next. In addition, we also examined the proportion of allele-specific reads per sample (i.e., total ASReC divided by total TReC) and filtered out samples with extreme proportions with study-specific thresholds.

Next we performed QC for allele-specific expression. If a gene had more than 10 allele-specific reads in a sample, we calculated the fraction of allele-specific reads mapped to the first haplotype, denoted by $h_{gi}$. If $|h_{gi} - 0.5| > 0.4$, we considered it as an extreme ASReC fraction. At the gene level, we examined the total ASReC across all samples as well as the total number of haplotype-ambiguous reads, if the fraction of haplotype-ambiguous reads was greater than 5%, we set a gene's ASReC to zero for all samples. Also, if the proportion of samples with extreme ASReC fractions was greater than 20% and mean proportion of haplotype-ambiguous reads was greater than 1%, we set that gene's ASReC to zero in all samples. Finally, for one gene and one sample, if the proportion of haplotype-ambiguous reads was greater than 10% and the number of allele-specific reads was greater than 10, we set its ASReC to zero.

From CMC, we selected the samples with both RNA-seq and genotype data, labeled as control or SCZ, and were not labeled as outlier by Fromer et al. [18]. Among the selected samples, we calculated the proportion of allele-specific reads per sample and excluded one sample with a low proportion less than 0.04. Next, we removed one sample with the proportion of haplotype-ambiguous reads being larger than 0.4%. These filtering criteria resulted in 537 samples (283 controls, 254 SCZ).

For BLUEPRINT, there were 548 samples with both RNA-seq and genotype data: 176 monocyte, 211 CD4+ T cell, and 161 neutrophil. With each purified cell type cohort having a mixture of paired and single-end read sequencing, we selected single-end sequencing samples for monocyte (n=173) and neutrophil (n=154) cohorts and paired-end sequencing samples for CD4+ T cell (n=208). None of the remaining samples appeared to deviate in terms of ASReC vs. TReC. When we inspected the proportion of haplotype-ambiguous reads, four T cell samples were excluded after applying a 0.25% cutoff. These criteria resulted in 531 samples (173 monocytes, 204 CD4+ T cells, 154 neutrophils).

For GTEx's BFC, there were 175 samples with both RNA-seq and genotype data. One sample appeared to have an extremely high TReC across genes and was excluded. When we compared ASReC vs TReC per sample, there were 17 samples with the proportion of allele-specific reads larger than 5.5%. Unlike the above filtering procedure, we retained these samples but set their ASReC to zero to maintain a similar sample size with the recent GTEx eQTL analysis [25] resulting in 174 samples.

For WB, GTEx contained 670 samples with available RNA-seq and genotype data. None of the samples were considered outliers based on TReC across genes. When we compared ASReC versus TReC, in 68 samples the proportions of allele-specific reads were greater than 7%. Similar to BFC, rather than excluding these samples, we retained them but set their ASReC to zero to maintain the sample sizes in the GTEx analysis [25].

| Cohort | Counts | | | | SNPs per Gene | | |
|---|---|---|---|---|---|---|---|
| | Samples | Genes | SNPs | Gene+SNP pairs | Q25 | Q50 | Q75 |
| CommonMind Consortium - Control | 275 | 15,334 | 5,792,709 | 32,523,198 | 1,661 | 2,045 | 2,462 |
| CommonMind Consortium - SCZ | 250 | 15,275 | 5,792,709 | 32,186,224 | 1,645 | 2,036 | 2,450 |
| GTEx Brain | 174 | 15,400 | 5,938,874 | 33,326,549 | 1,665 | 2,093 | 2,546 |
| BLUEPRINT - CD4T | 167 | 13,339 | 4,851,911 | 22,532,356 | 1,304 | 1,672 | 2,035 |
| BLUEPRINT - Monocyte | 173 | 13,624 | 4,851,911 | 23,101,424 | 1,315 | 1,680 | 2,034 |
| BLUEPRINT - Neutrophil | 151 | 10,769 | 4,851,911 | 18,044,122 | 1,298 | 1,662 | 2,016 |
| GTEx Blood | 670 | 12,022 | 5,938,874 | 26,426,763 | 1,697 | 2,124 | 2,581 |

Table 1: **Summary of sample size, the number of genes, SNPs, local gene-SNP pairs, and the number of local SNPs per gene.** Q25, Q50, and Q75 indicate 1st quartile, median and 3rd quartile.

### 6 Real Data Analyses

#### 6.1 Covariates

For bulk and cell type-specific models with CMC, covariates included log transformed 75th percentile of TReC (library size), sex, institution, binned library cluster obtained from the supplemental materials of Fromer et al. [18], age at death, postmortem interval, RNA integrity number, RNA integrity number squared, and the top five genotype principal components.

For the BLUEPRINT dataset, covariates included library size, sex, donor age, submission year, % unique reads, % coding bases, % UTR bases, the top five genotype principal components, and the top sixteen residual TReC principal components per purified cell type.

For the GTEx WB dataset, covariates included library size, sex, sequencing protocol, sequencing platform, sequencing center, RNA integrity number, RNA integrity number squared, donor age, base mismatch rate, % reads mapped within exons, number of mapped read pairs, mean fragment length, % intronic reads, % rRNA reads, number of genes detected with $\geq 5$ exon reads, end 1 mismatch rate (fraction of end 1 bases not matching the reference), fragment length standard deviation, intergenic rate, number of rRNA reads, number of reads labeled failed by sequencer, ratio of exon reads to total reads, end 2 mismatch rate (fraction of end 2 bases not matching the reference), end 2 % sense (percentage of intragenic end 2 reads sequenced in the sense direction), and the top ten genotype principal components.

For the GTEx brain dataset, covariates included library size, sex, sequencing protocol, % rRNA reads, end 2 mapping rate, intergenic rate, end 2 % sense, binned year of nucleic acid isolation batch, RNA integrity number, fragment length standard deviation, total filtered reads, number of alternate alignments, number of reads spanning an exon-exon boundary, end 1 % sense, and the top three genotype principal components. GTEx dataset variables and data dictionary were obtained at **https://www.gtexportal.org/home/datasets**.

For CMC and GTEx datasets, we also calculated PCs based on log-transformed TReC data. We first fit a linear model of log transformed TReC data on all the aforementioned covariates, and then calculated PCs (or surrogate variables) using the residuals of the linear model. In addition, we also considered another linear model that included aforementioned covariates as well as cell type proportions, and then calculated PCs (or surrogate variables) using the residuals of this larger model. Therefore, these two sets of PCs capture the latent batch effects that were ignorant or orthogonal to cell type proportion. For CMC, we used the top twelve surrogate variables calculated independent of disease status with SVA [26]. For GTEx WB and brain samples, latent effects were captured by the top 20 PCs and top 16 PCs from residualized TReC, respectively.

#### 6.2 Commonmind Consortium

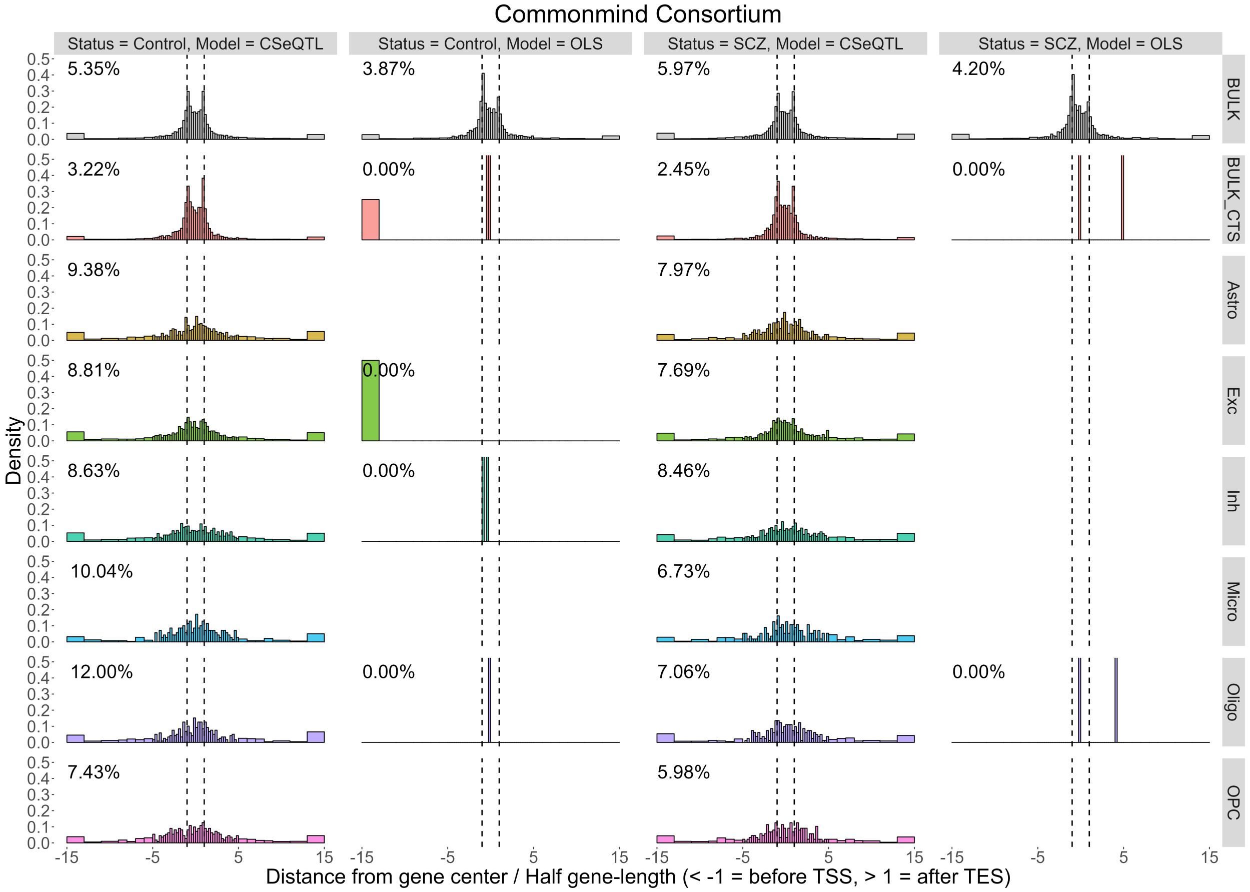

Figure 11: **The distribution of eQTLs from the CMC study.** The distribution of normalized distance between each eGene (the center of its gene body) and its minimum p-value SNP. The raw distance (base pairs) were normalized by dividing by half of the gene size. Therefore $[-1, 1]$ was the range of gene body. Percentages of eGene for which such distances were beyond 15 units were provided per facet plot. BULK refers to the eGenes detected by bulk eQTL mapping while BULK_CTS refers to all bulk eGenes identified as cell type-specific eGenes in at least one cell type.

| TRIM | COHORT | MODEL | CELLTYPE | Perm PVAL | ERROR | eGene | OL_eGene | eQTL | OL_eQTL |
|---|---|---|---|---|---|---|---|---|---|
| FALSE | Control | CSeQTL | Bulk | 2.20e-02 | 0.07 | 8,116 | - | 835,196 | - |
| | | | Astro | 3.22e-04 | 0.05 | 716 | 70.17% | 7,606 | 73.49% |
| | | | Exc | 1.76e-03 | 0.05 | 2,511 | 75.23% | 65,321 | 81.91% |
| | | | Inh | 5.12e-04 | 0.05 | 1,133 | 61.61% | 11,125 | 34.88% |
| | | | Micro | 1.66e-04 | 0.05 | 264 | 68.94% | 9,460 | 93.50% |
| | | | Oligo | 3.15e-04 | 0.05 | 495 | 71.92% | 5,525 | 78.75% |
| | | | OPC | 2.01e-04 | 0.04 | 392 | 60.20% | 3,310 | 60.00% |
| | | OLS | Bulk | 3.28e-03 | 0.05 | 3,812 | - | 374,302 | - |
| | | | Astro | 1.58e-06 | 0.06 | 0 | - | 0 | - |
| | | | Exc | 7.05e-08 | 0.05 | 1 | 100.00% | 1 | 100.00% |
| | | | Inh | 5.85e-07 | 0.05 | 2 | 100.00% | 9 | 100.00% |
| | | | Micro | 1.20e-06 | 0.06 | 0 | - | 0 | - |
| | | | Oligo | 1.16e-06 | 0.05 | 2 | 100.00% | 45 | 100.00% |
| | | | OPC | 7.83e-07 | 0.06 | 0 | - | 0 | - |
| | SCZ | CSeQTL | Bulk | 1.78e-02 | 0.07 | 7,228 | - | 704,913 | - |
| | | | Astro | 3.16e-04 | 0.05 | 626 | 61.50% | 6,373 | 78.94% |
| | | | Exc | 1.48e-03 | 0.06 | 1,888 | 72.62% | 51,864 | 89.80% |
| | | | Inh | 4.44e-04 | 0.05 | 979 | 57.46% | 8,059 | 18.33% |
| | | | Micro | 1.25e-04 | 0.06 | 183 | 60.66% | 1,894 | 74.76% |
| | | | Oligo | 2.02e-04 | 0.05 | 300 | 62.33% | 4,533 | 83.76% |
| | | | OPC | 1.60e-04 | 0.04 | 334 | 57.66% | 2,157 | 64.44% |
| | | OLS | Bulk | 2.17e-03 | 0.05 | 2,919 | - | 285,391 | - |
| | | | Astro | 5.20e-06 | 0.07 | 0 | - | 0 | - |
| | | | Exc | 2.57e-06 | 0.06 | 2 | 100.00% | 63 | 100.00% |
| | | | Inh | 5.77e-06 | 0.06 | 0 | - | 0 | - |
| | | | Micro | 3.04e-06 | 0.07 | 2 | 0.00% | 16 | 0.00% |
| | | | Oligo | 1.11e-06 | 0.06 | 3 | 66.67% | 20 | 85.00% |
| | | | OPC | 2.46e-05 | 0.06 | 0 | - | 0 | - |
| TRUE | Control | CSeQTL | Bulk | 1.90e-02 | 0.07 | 7,780 | - | 807,352 | - |
| | | | Astro | 3.20e-04 | 0.04 | 725 | 65.10% | 7,281 | 69.32% |
| | | | Exc | 1.50e-03 | 0.05 | 2,337 | 74.27% | 57,499 | 80.70% |
| | | | Inh | 5.48e-04 | 0.05 | 1,217 | 58.34% | 10,893 | 24.22% |
| | | | Micro | 1.65e-04 | 0.05 | 279 | 62.72% | 9,241 | 93.15% |
| | | | Oligo | 2.90e-04 | 0.05 | 475 | 70.11% | 5,872 | 79.00% |
| | | | OPC | 2.29e-04 | 0.04 | 444 | 57.88% | 2,849 | 57.11% |
| | | OLS | Bulk | 3.14e-03 | 0.05 | 3,774 | - | 374,638 | - |
| | | | Astro | 1.73e-05 | 0.05 | 0 | - | 0 | - |
| | | | Exc | 7.05e-08 | 0.05 | 1 | 100.00% | 1 | 100.00% |
| | | | Inh | 4.10e-07 | 0.05 | 2 | 100.00% | 7 | 100.00% |
| | | | Micro | 4.71e-05 | 0.06 | 0 | - | 0 | - |
| | | | Oligo | 7.90e-07 | 0.05 | 1 | 100.00% | 23 | 100.00% |
| | | | OPC | 2.72e-05 | 0.06 | 0 | - | 0 | - |
| | SCZ | CSeQTL | Bulk | 1.53e-02 | 0.07 | 6,885 | - | 680,924 | - |
| | | | Astro | 2.56e-04 | 0.05 | 552 | 58.33% | 6,197 | 78.59% |
| | | | Exc | 1.21e-03 | 0.06 | 1,716 | 69.80% | 49,403 | 84.32% |
| | | | Inh | 4.49e-04 | 0.05 | 1,016 | 49.61% | 9,423 | 13.76% |
| | | | Micro | 1.37e-04 | 0.05 | 223 | 47.09% | 3,007 | 84.87% |
| | | | Oligo | 2.08e-04 | 0.05 | 326 | 61.04% | 5,203 | 83.05% |
| | | | OPC | 1.75e-04 | 0.04 | 351 | 53.85% | 2,070 | 61.11% |
| | | OLS | Bulk | 2.12e-03 | 0.05 | 2,902 | - | 286,245 | - |
| | | | Astro | 1.12e-04 | 0.06 | 0 | - | 0 | - |
| | | | Exc | 4.82e-07 | 0.06 | 0 | - | 0 | - |
| | | | Inh | 8.75e-06 | 0.06 | 0 | - | 0 | - |
| | | | Micro | 1.67e-05 | 0.06 | 0 | - | 0 | - |
| | | | Oligo | 9.05e-08 | 0.05 | 2 | 100.00% | 10 | 100.00% |
| | | | OPC | 2.18e-05 | 0.06 | 0 | - | 0 | - |

Table 2: **Commonmind Consortium analysis.** Perm PVAL is the permutation p-value cutoff for q-value 0.005. Type I Error (ERROR) corresponds to proportion of eQTLs identified at p-value 0.05 after permuting SNP genotype data. OL_eGene and OL_eQTL are the percent of ct-eGenes and ct-eQTLs that are discovered by bulk eQTL mapping.

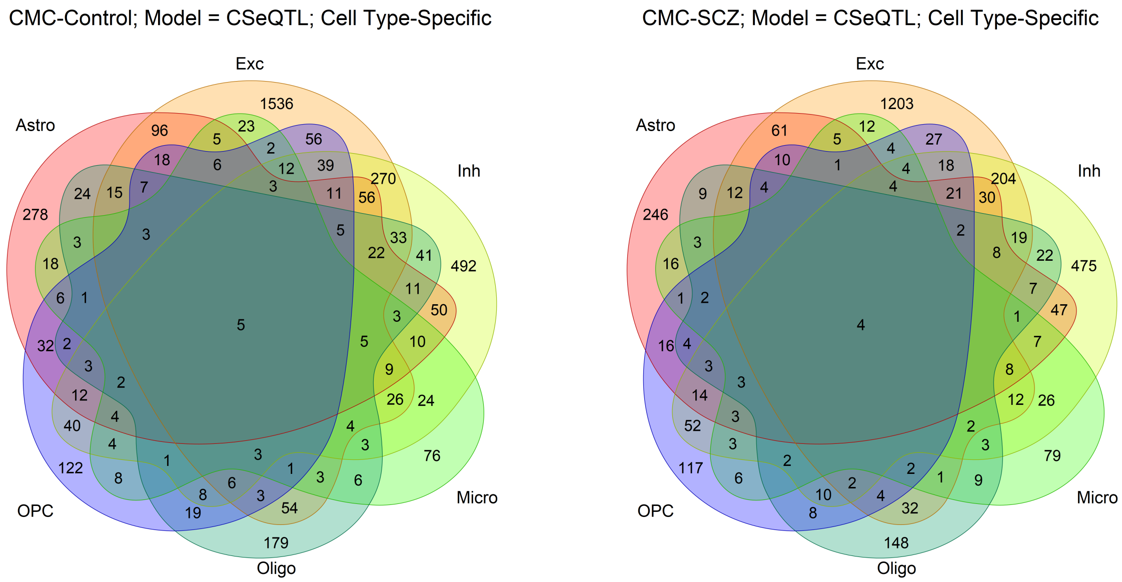

Figure 12: **Overlapping eGenes among Commonmind Consortium control samples (left panel) and schizophrenia samples (right panel).**

##### 6.3 GTEx Brain

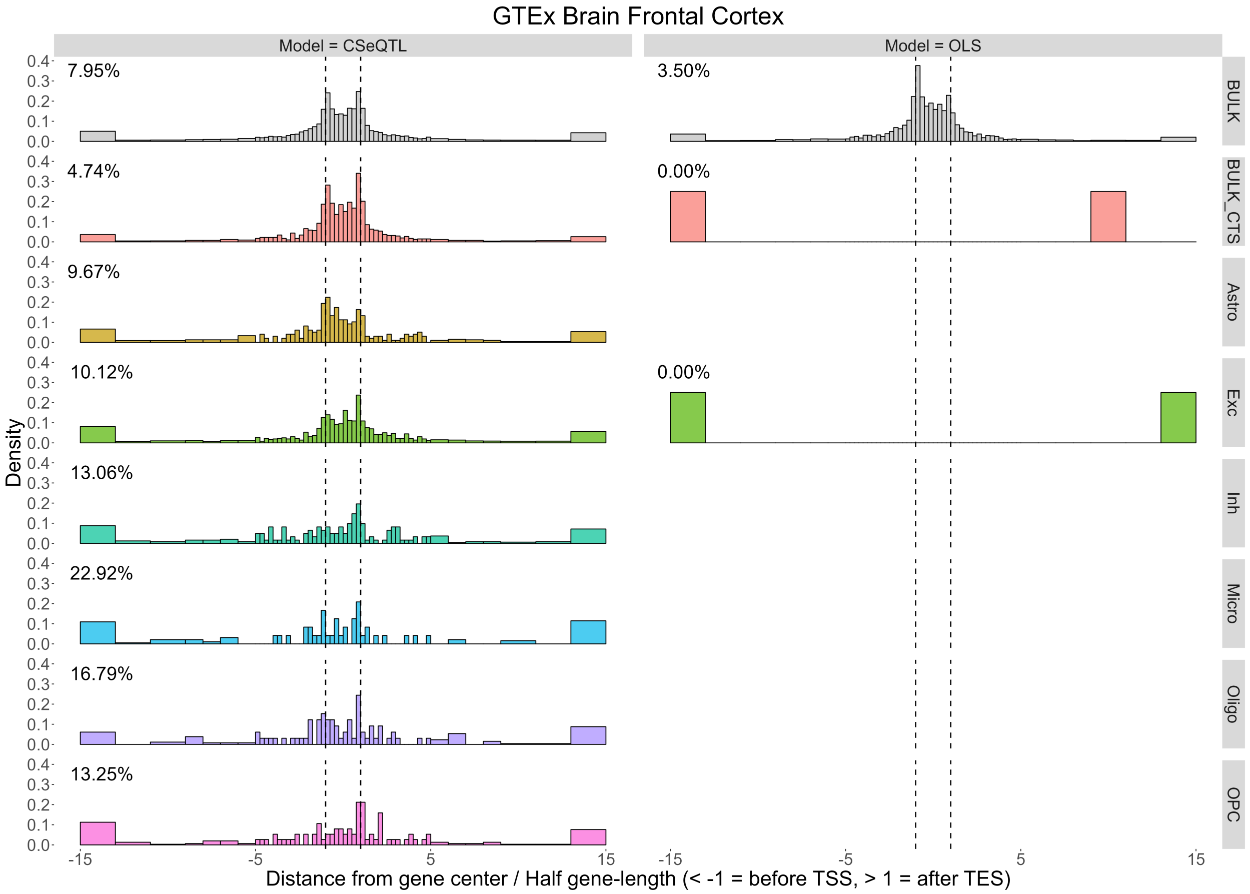

Figure 13: **The distribution of eQTLs from the GTEx brain study.** The distribution of normalized distance between each eGene (the center of its gene body) and its minimum p-value SNP. The raw distance (base pairs) were normalized by dividing by half of the gene size. Therefore $[-1, 1]$ was the range of gene body. Percentages of eGene for which such distances were beyond 15 units were provided per facet plot. BULK refers to the eGenes detected by bulk eQTL mapping while BULK_CTS refers to all bulk eGenes identified as cell type-specific eGenes in at least one cell type.

| TRIM | MODEL | CELLTYPE | Perm PVAL | ERROR | eGene | OL_eGene | eQTL | OL_eQTL |
|---|---|---|---|---|---|---|---|---|
| FALSE | CSeQTL | Bulk | 3.24e-02 | 0.09 | 8,769 | - | 718,012 | - |
| | | Astro | 6.40e-04 | 0.08 | 748 | 76.78% | 13,551 | 76.84% |
| | | Exc | 2.06e-03 | 0.08 | 1,780 | 77.25% | 44,562 | 84.39% |
| | | Inh | 1.69e-04 | 0.06 | 265 | 71.70% | 1,981 | 58.35% |
| | | Micro | 7.76e-04 | 0.08 | 885 | 50.62% | 15,318 | 10.77% |
| | | Oligo | 3.90e-04 | 0.09 | 576 | 60.87% | 6,585 | 44.60% |
| | | OPC | 1.06e-04 | 0.06 | 200 | 69.00% | 2,655 | 80.38% |
| | OLS | Bulk | 1.80e-03 | 0.05 | 2,683 | - | 215,762 | - |
| | | Astro | 2.20e-05 | 0.07 | 0 | - | 0 | - |
| | | Exc | 4.86e-06 | 0.06 | 12 | 83.33% | 101 | 97.03% |
| | | Inh | 9.04e-06 | 0.05 | 0 | - | 0 | - |
| | | Micro | 1.09e-04 | 0.11 | 184 | 3.30% | 1,332 | 0.00% |
| | | Oligo | 9.39e-07 | 0.07 | 3 | - | 0 | - |
| | | OPC | 7.31e-07 | 0.07 | 0 | - | 0 | - |
| TRUE | CSeQTL | Bulk | 2.97e-02 | 0.08 | 8,362 | - | 667,554 | - |
| | | Astro | 2.42e-04 | 0.06 | 393 | 79.90% | 6,910 | 83.44% |
| | | Exc | 1.35e-03 | 0.07 | 1,433 | 77.60% | 37,734 | 84.46% |
| | | Inh | 1.40e-04 | 0.06 | 245 | 70.49% | 2,144 | 63.34% |
| | | Micro | 5.15e-05 | 0.06 | 96 | 66.32% | 1,650 | 84.61% |
| | | Oligo | 6.55e-05 | 0.06 | 131 | 67.69% | 1,698 | 78.62% |
| | | OPC | 7.29e-05 | 0.05 | 151 | 60.67% | 2,090 | 82.73% |
| | OLS | Bulk | 1.75e-03 | 0.05 | 2,626 | - | 207,562 | - |
| | | Astro | 1.15e-04 | 0.04 | 0 | - | 0 | - |
| | | Exc | 1.62e-06 | 0.05 | 2 | 100.00% | 10 | 100.00% |
| | | Inh | 4.58e-05 | 0.05 | 0 | - | 0 | - |
| | | Micro | 1.28e-05 | 0.04 | 0 | - | 0 | - |
| | | Oligo | 1.25e-05 | 0.05 | 0 | - | 0 | - |
| | | OPC | 6.78e-05 | 0.04 | 0 | - | 0 | - |

Table 3: **GTEx Brain Frontal Cortex (BA9) analysis.** Perm PVAL is the permutation p-value cutoff for q-value 0.005. Type I Error (ERROR) corresponds to proportion of eQTLs identified at p-value 0.05 after permuting SNP genotype data. OL_eGene and OL_eQTL are the percent of ct-eGenes and ct-eQTLs that are discovered by bulk eQTL mapping.

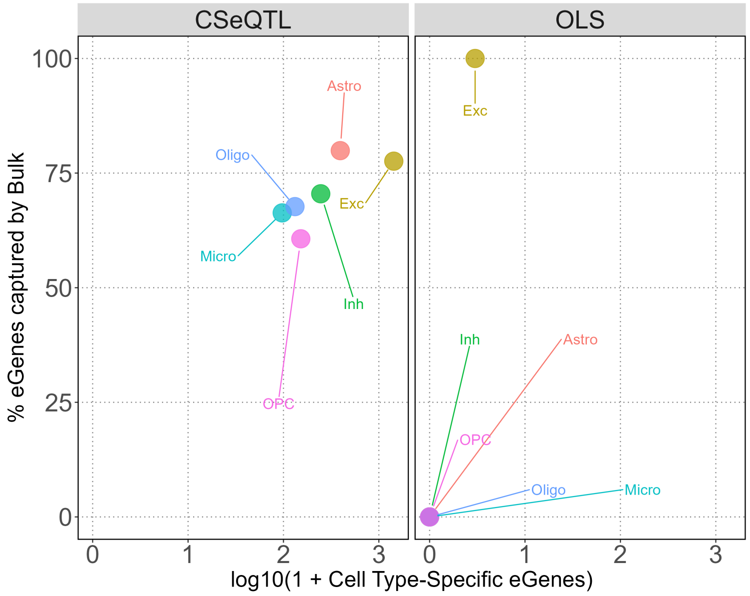

Figure 14: **The number of eGenes from the GTEx brain study.** Comparison of the number of eGenes identified by CSeQTL or OLS method (X-axis) and what percentage overlap with the eGenes from bulk eQTL mapping.

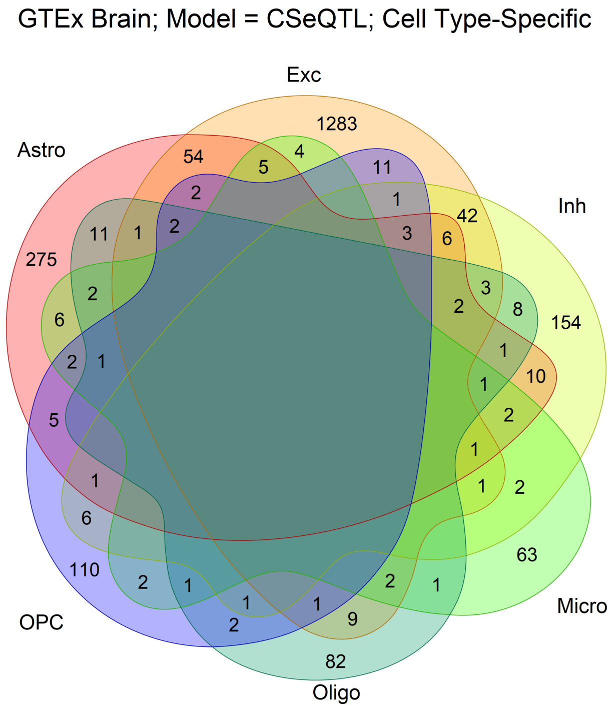

Figure 15: **Overlapping eGenes among GTEx Brain samples.**

#### 6.4 BLUEPRINT

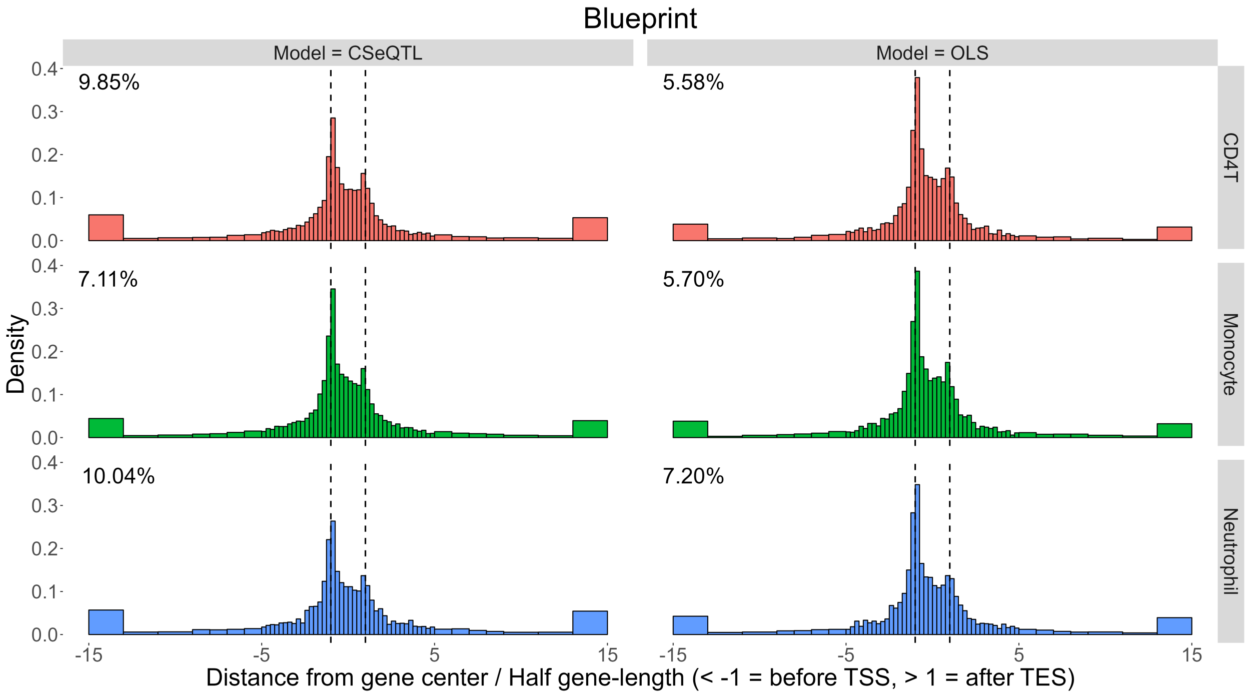

Figure 16: **The distribution of eQTLs from the BLUEPRINT study.** The distribution of normalized distance between each eGene (the center of its gene body) and its minimum p-value SNP. The raw distance (base pairs) were normalized by dividing by half of the gene size. Therefore $[-1, 1]$ was the range of gene body. Percentages of eGene for which such distances were beyond 15 units were provided per facet plot.

| TRIM | MODEL | CELLTYPE | Perm PVAL | ERROR | eGene | eQTL |
|---|---|---|---|---|---|---|
| FALSE | CSeQTL | CD4T | 5.38e-02 | 0.08 | 9,450 | 983,572 |
| | OLS | | 6.28e-03 | 0.05 | 5,268 | 459,610 |
| | CSeQTL | Monocyte | 1.36e-02 | 0.08 | 7,485 | 762,871 |
| | OLS | | 6.57e-03 | 0.05 | 5,521 | 486,071 |
| | CSeQTL | Neutrophil | 2.29e-02 | 0.08 | 6,567 | 642,900 |
| | OLS | | 5.62e-03 | 0.05 | 3,997 | 335,550 |
| TRUE | CSeQTL | CD4T | 5.10e-02 | 0.08 | 9,342 | 969,848 |
| | OLS | | 6.21e-03 | 0.05 | 5,214 | 455,354 |
| | CSeQTL | Monocyte | 1.24e-02 | 0.07 | 7,304 | 750,037 |
| | OLS | | 6.43e-03 | 0.05 | 5,470 | 481,877 |
| | CSeQTL | Neutrophil | 2.06e-02 | 0.08 | 6,402 | 628,963 |
| | OLS | | 5.56e-03 | 0.05 | 3,972 | 332,108 |

Table 4: BLUEPRINT eQTL mapping results. Perm PVAL is the permutation p-value cutoff for q-value 0.005. Type I Error (ERROR) corresponds to proportion of eQTLs identified at p-value 0.05 after permuting SNP genotype data.

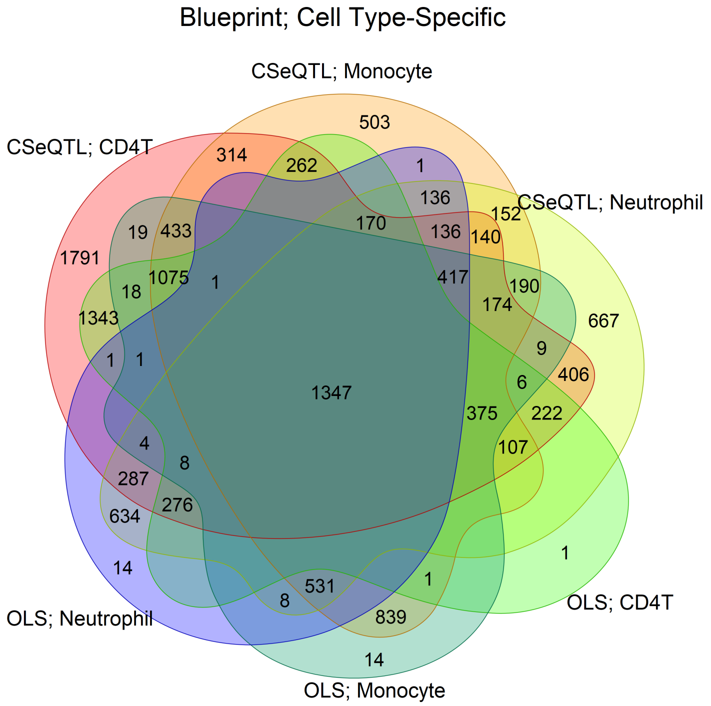

Figure 17: **Overlapping eGenes among BLUEPRINT samples between cell types and models.**

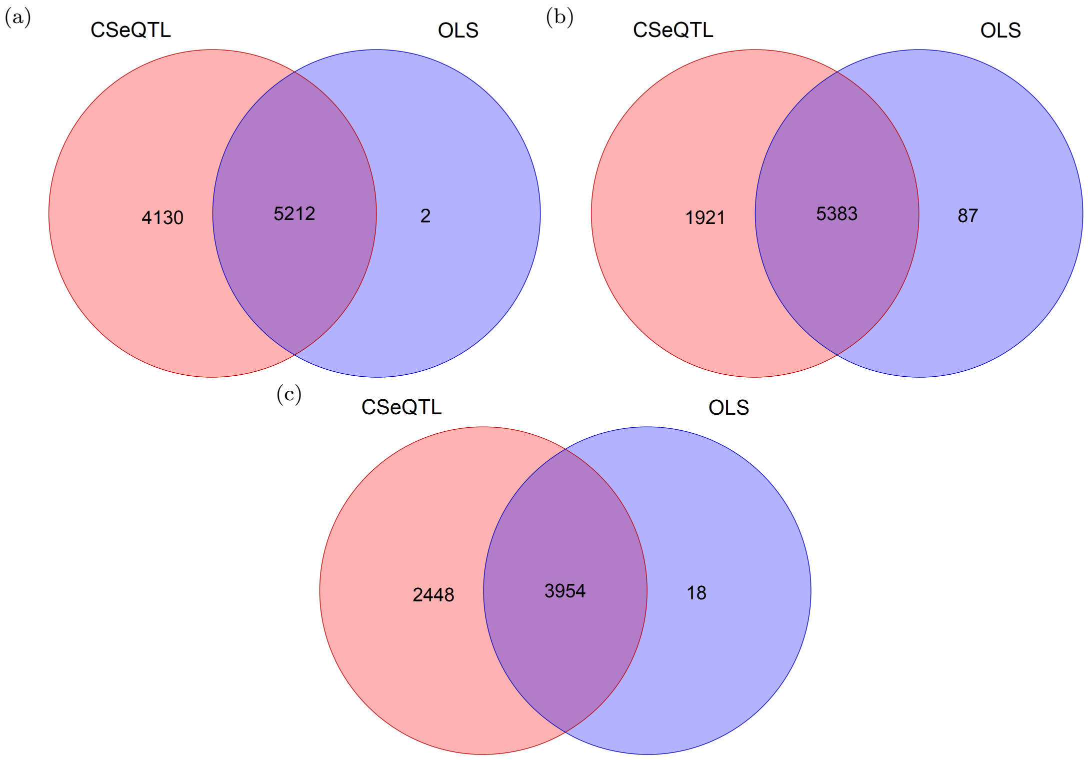

Figure 18: **Shared and exclusive BLUEPRINT eGenes per cell type and by model.** (a) CD4+ T cells. (b) Monocytes. (c) Neutrophils.

#### 6.5 GTEx Whole Blood

| TRIM | MODEL | CELLTYPE | Perm PVAL | ERROR | eGene | OL_eGene | eQTL | OL_eQTL |
|---|---|---|---|---|---|---|---|---|
| FALSE | CSeQTL | Bulk | 4.75e-02 | 0.07 | 9,079 | - | 1,422,635 | - |
| | | CD4T | 1.26e-04 | 0.05 | 255 | 86.27% | 4,229 | 75.05% |
| | | CD8T | 1.06e-04 | 0.05 | 172 | 90.70% | 2,041 | 79.42% |
| | | B_Cell | 1.04e-04 | 0.05 | 130 | 85.38% | 3,182 | 71.84% |
| | | Monocyte | 4.05e-04 | 0.06 | 438 | 89.73% | 15,138 | 70.93% |
| | | NK | 3.72e-04 | 0.06 | 486 | 89.92% | 14,193 | 82.58% |
| | | Neutrophil | 7.50e-03 | 0.07 | 4,371 | 95.29% | 391,262 | 86.68% |
| | | Mast_Eosinophil | 1.88e-03 | 0.10 | 1,405 | 85.05% | 37,853 | 64.58% |
| | OLS | Bulk | 8.37e-03 | 0.05 | 5,521 | - | 699,125 | - |
| | | CD4T | 5.05e-05 | 0.08 | 91 | 46.07% | 875 | 8.57% |
| | | CD8T | 1.43e-06 | 0.05 | 0 | - | 0 | - |
| | | B_Cell | 4.90e-07 | 0.07 | 1 | - | 0 | - |
| | | Monocyte | 4.93e-05 | 0.08 | 59 | 38.98% | 553 | 34.00% |
| | | NK | 3.65e-06 | 0.06 | 0 | - | 0 | - |
| | | Neutrophil | 1.29e-03 | 0.07 | 1,125 | 93.07% | 51,090 | 97.00% |
| | | Mast_Eosinophil | 1.93e-04 | 0.09 | 160 | 48.75% | 1,143 | 26.07% |
| TRUE | CSeQTL | Bulk | 4.01e-02 | 0.06 | 8,822 | - | 1,364,160 | - |
| | | CD4T | 1.15e-04 | 0.04 | 223 | 85.65% | 3,809 | 76.53% |
| | | CD8T | 1.22e-04 | 0.04 | 202 | 83.66% | 2,394 | 72.64% |
| | | B_Cell | 1.00e-04 | 0.04 | 145 | 90.28% | 2,899 | 84.37% |
| | | Monocyte | 3.74e-04 | 0.06 | 437 | 90.85% | 17,928 | 71.64% |
| | | NK | 2.72e-04 | 0.05 | 436 | 92.20% | 14,173 | 83.03% |
| | | Neutrophil | 6.09e-03 | 0.07 | 4,097 | 95.46% | 362,627 | 86.65% |
| | | Mast_Eosinophil | 5.20e-04 | 0.07 | 579 | 90.33% | 17,228 | 82.57% |
| | OLS | Bulk | 8.39e-03 | 0.05 | 5,586 | - | 702,162 | - |
| | | CD4T | 2.97e-05 | 0.04 | 0 | - | 0 | - |
| | | CD8T | 4.24e-05 | 0.05 | 0 | - | 0 | - |
| | | B_Cell | 1.65e-05 | 0.05 | 0 | - | 0 | - |
| | | Monocyte | 9.81e-10 | 0.06 | 1 | - | 0 | - |
| | | NK | 3.00e-05 | 0.05 | 0 | - | 0 | - |
| | | Neutrophil | 8.75e-04 | 0.06 | 1,014 | 96.94% | 47,168 | 97.84% |
| | | Mast_Eosinophil | 2.61e-07 | 0.07 | 2 | 100.00% | 1 | 100.00% |

Table 5: **GTEx Whole Blood analysis.** Perm PVAL is the permutation p-value cutoff for q-value 0.005. Type I Error (ERROR) corresponds to proportion of eQTLs identified at p-value 0.05 after permuting SNP genotype data. OL_eGene and OL_eQTL are the percent of ct-eGenes and ct-eQTLs that are discovered by bulk eQTL mapping.

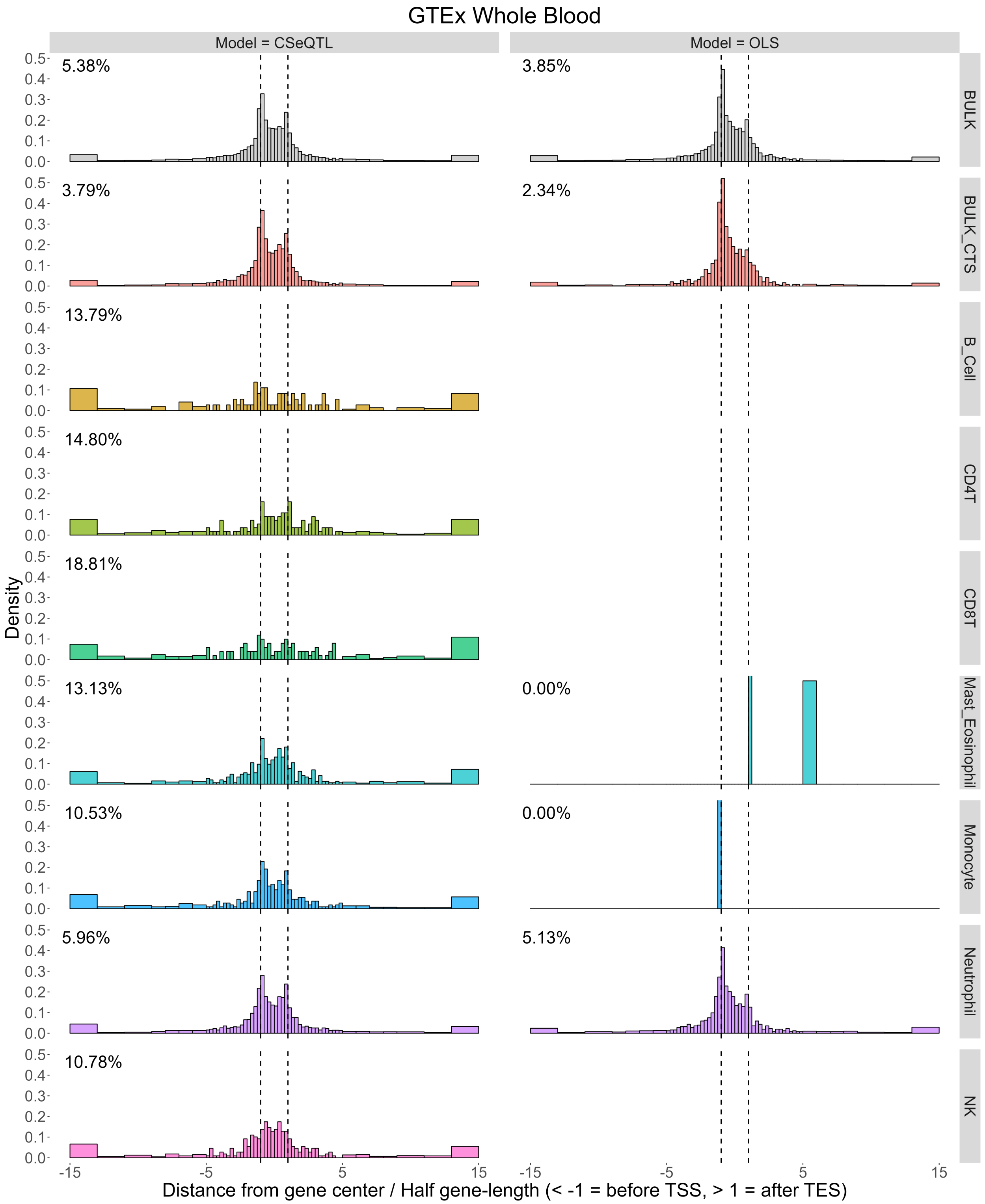

Figure 19: **The distribution of eQTLs from the GTEx whole blood study.** The distribution of normalized distance between each eGene (the center of its gene body) and its minimum p-value SNP. The raw distance (base pairs) were normalized by dividing by half of the gene size. Therefore $[-1, 1]$ was the range of gene body. Percentages of eGene for which such distances were beyond 15 units were provided per facet plot. BULK refers to the eGenes detected by bulk eQTL mapping while BULK_CTS refers to all bulk eGenes identified as cell type-specific eGenes in at least one cell type.

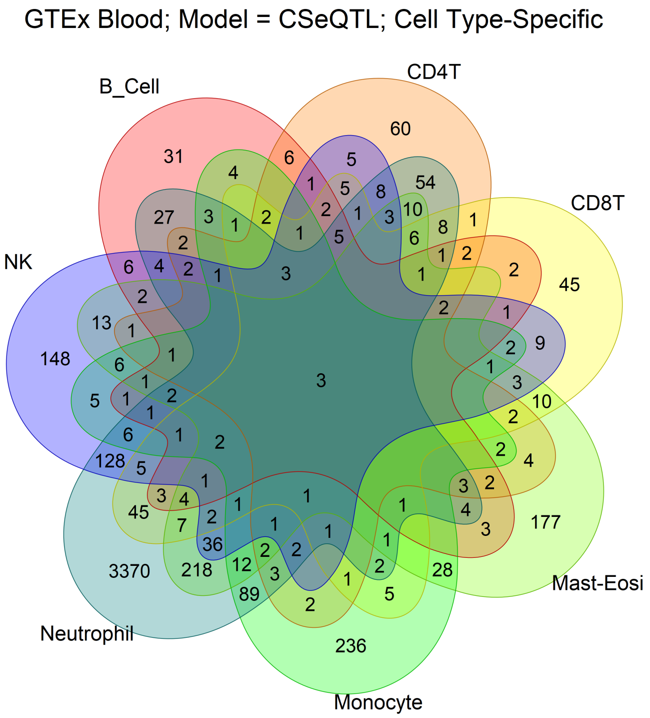

Figure 20: **Overlapping eGenes among GTEx Blood samples.**

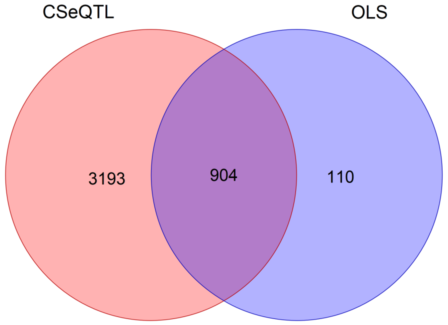

Figure 21: **Overlapping eGenes among GTEx Blood samples for neutrophils between models.**

#### 6.6 Validation

In addition to purified bulk RNA-seq data from BLUEPRINT, we also validated our method using sc-eQTLs from a scRNA-seq dataset of PBMC samples [27] and a snRNA-seq dataset of brain samples [28].

We obtained ct-eQTLs in 14 immune cell types reported by Yazar et al. [27]. They collected scRNA-seq data for almost 1,000 individuals (982 donors in total) with an average of 1,291 cells per donor, and named their cohort as OneK1K. Genotype data were collected from more than 5.3 million SNPs. ScRNA-seq data were generated using a pooled multiplexing strategy where each pool included cells from 12-14 donors. Cells were assigned to each donor using their genotype data. Gene expression data were normalized by SCTransform method [29] to remove the effect of read-depth, sequencing pool, as well as percentage of mitochondrial expression. The average expression of each gene and per person was calculated using the normalized counts. The covariates included were age, sex, six genotype PCs, and two PEER factors. They searched for local eQTLs within 1Mb of the gene, including gene body, by Spearman's rank correlation using residual expression. The FDR was evaluated for each chromosome separately, instead of a genome-wide search.

The other set of ct-eQTL results were derived by snRNA-seq data from brain samples. Bryois et al. [28] collected gene expression data of 6,940–14,595 genes and genotypes for 5.3 million SNPs in 144–192 individuals for eight major brain cell types: Excitatory neurons, Inhibitory neurons, Astrocytes, Microglia, Oligodendrocytes, Oligodendrocyte precursor cells (OPCs), Endothelial, and Pericytes. They searched for local eQTLs within 1Mb of the transcription start site (TSS). For each cell type and each individual, Bryois et al. [28] generated pseudo-bulk gene expression data by summing all counts for each gene. eQTL mapping was conducted by the fastQTL method that estimated the distribution of the minimum p-value per gene under the null (no eQTL), and then the authors calculated the permutation p-value for the minimum p-value per gene. Genes were filtered based on their average expression level ($\geq 1$ count per million) and proportion of ambient RNA ($< 10\%$). They used the following covariates: three first genotyping PCs, disease status (MS, AD, control or other), study (Roche_MS, Roche_AD, Columbia_AD, Mathys_AD and Zhou_AD) and 70 first expression PCs.

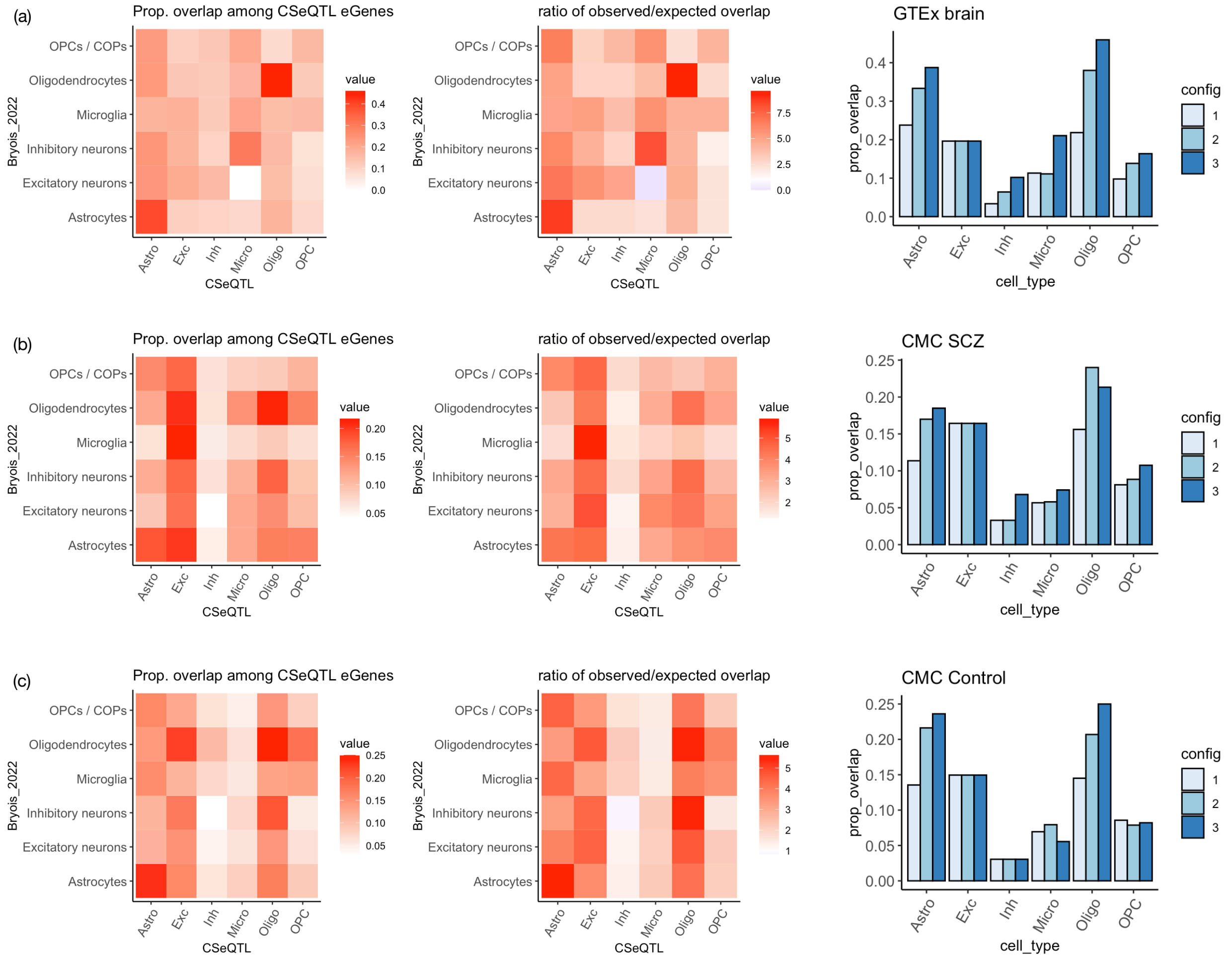

Figure 22: **Validation of CSeQTL results brain bulk RNA-seq data** We compare the results by CSeQTL versus snRNA-seq data [28] in terms of proportion of overlap (among CSeQTL eGenes), the ratio of observed versus expected overlap and the trend of overlap with three configurations to define CSeQTL eGenes: (1) q-value $< 0.005$; (2) q-value $< 0.005$ and fold change $\geq 1.5$; and (3) q-value $< 0.001$ and fold change $\geq 1.5$. There are CSeQTL results for three brain bulk RNA-seq datasets: (a) GTEx brain, (b) CMC schizophrenia (SCZ), and (c) CMC controls.

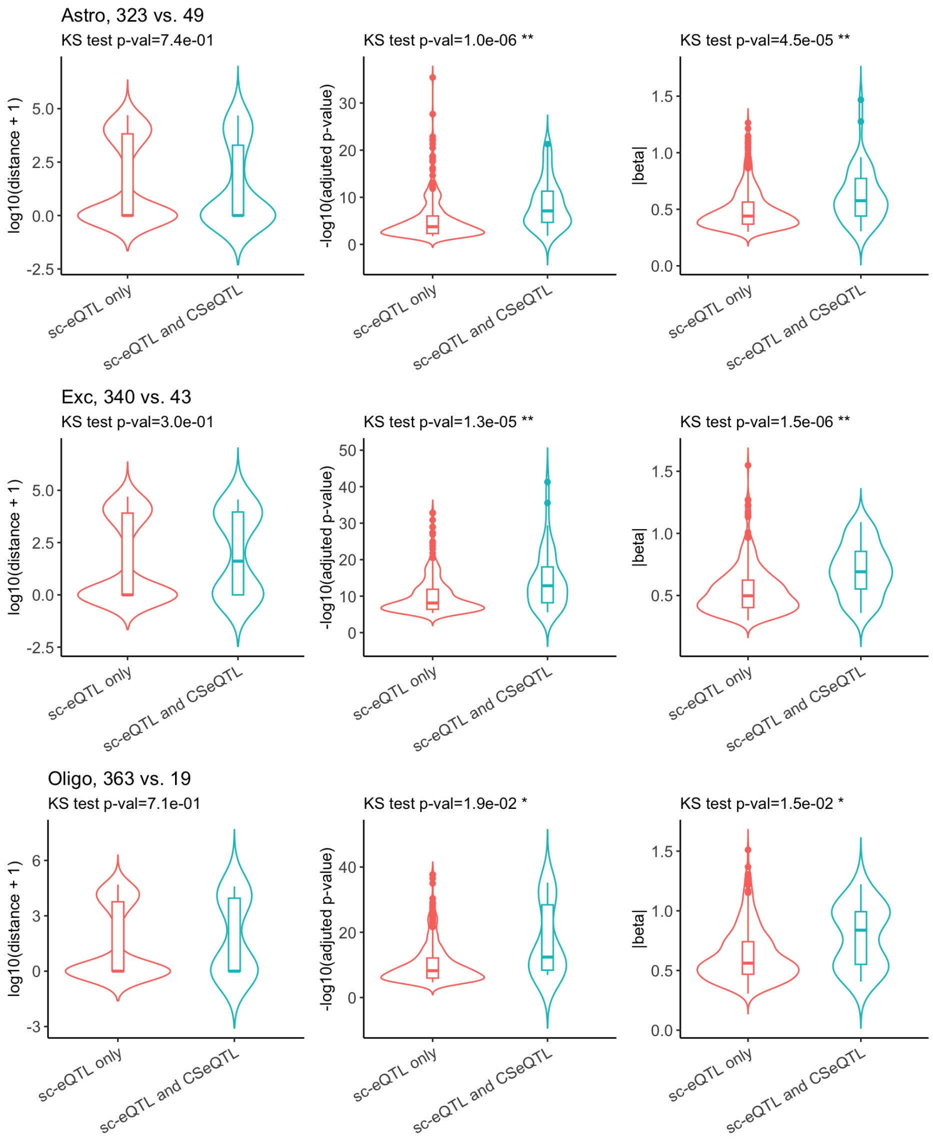

Figure 23: **Characteristics of eGenes identified in sc-eQTL analysis.** Among the eGenes identified by scRNA-seq data [28], compare those identified by sc-eQTL only versus those identified by both sc-eQTL and CSeQTL using GTEx brain data. Each row corresponds to a cell type. For example, in the first row, the title "Astro 323 vs. 49" indicates comparison of 323 eGenes identified by scRNA-seq only versus 49 eGenes identified by both scRNA-seq and CSeQTL. Three columns correspond to three comparisons: distance between the eSNP with the smallest p-value to the corresponding eGene, which is 0 if the eSNP is within the eGene, adjusted p-value, and absolute value of effect size. For each comparison, we conducted Kolmogorov–Smirnov test and significant results were labeled by ** (p-value $< 0.01$) or * ($0.01 \leq$ p-value $< 0.05$).

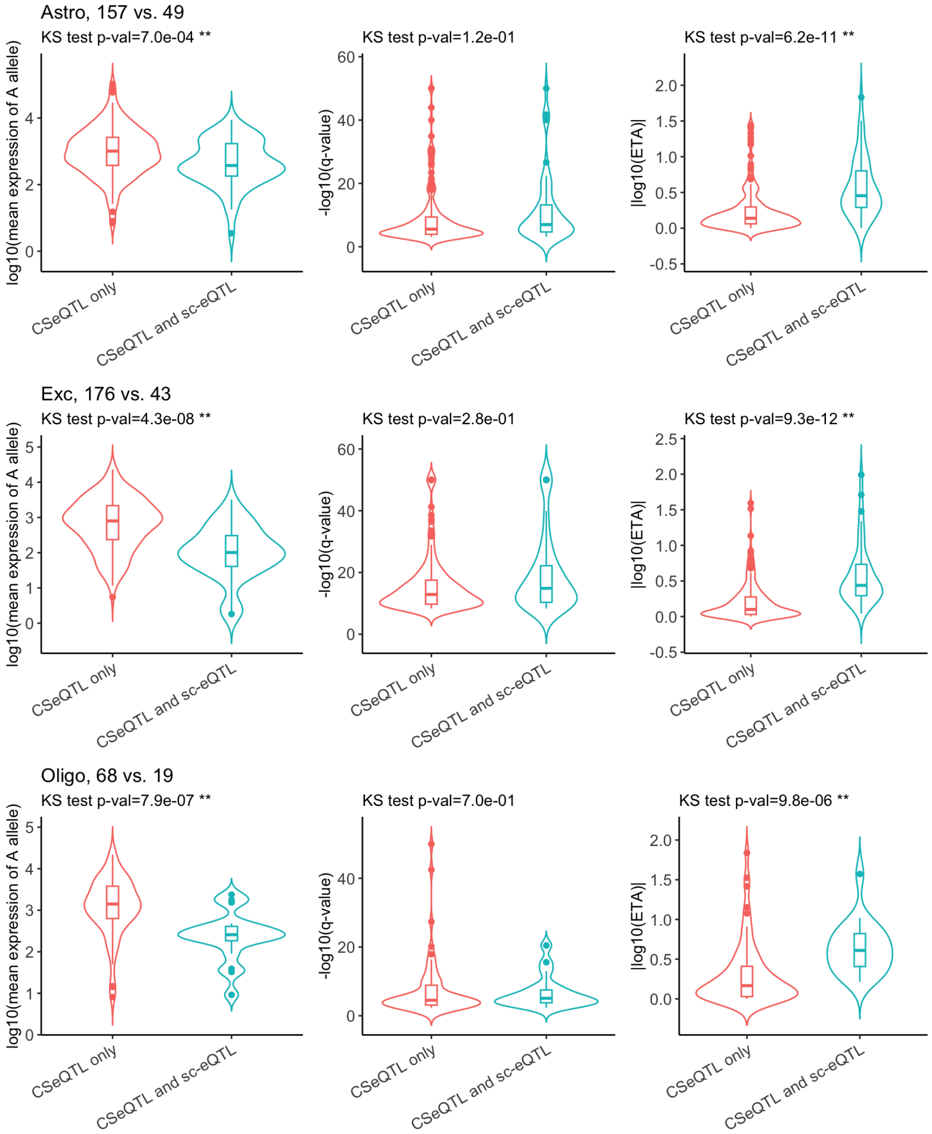

Figure 24: **Characteristics of eGenes identified in CSeQTL analysis.** Among the eGenes identified by CSeQTL method using GTEx brain dat, compare those identified by CSeQTL only versus those identified by both CSeQTL and sc-eQTL. Each row corresponds to a cell type. For example, in the first row, the title "Astro 157 vs. 49" indicates comparison of 157 eGenes identified by CSeQTL only versus 49 eGenes identified by both CSeQTL and sc-eQTL. Three columns correspond to three comparisons: base-line expression of reference allele (A allele), q-value, and absolute value of effect size. Here effect size is quantified by ETA which is fold change. For each comparison, we conducted Kolmogorov–Smirnov test and significant results were labeled by ** (p-value $< 0.01$) or * ($0.01 \leq$ p-value $< 0.05$).

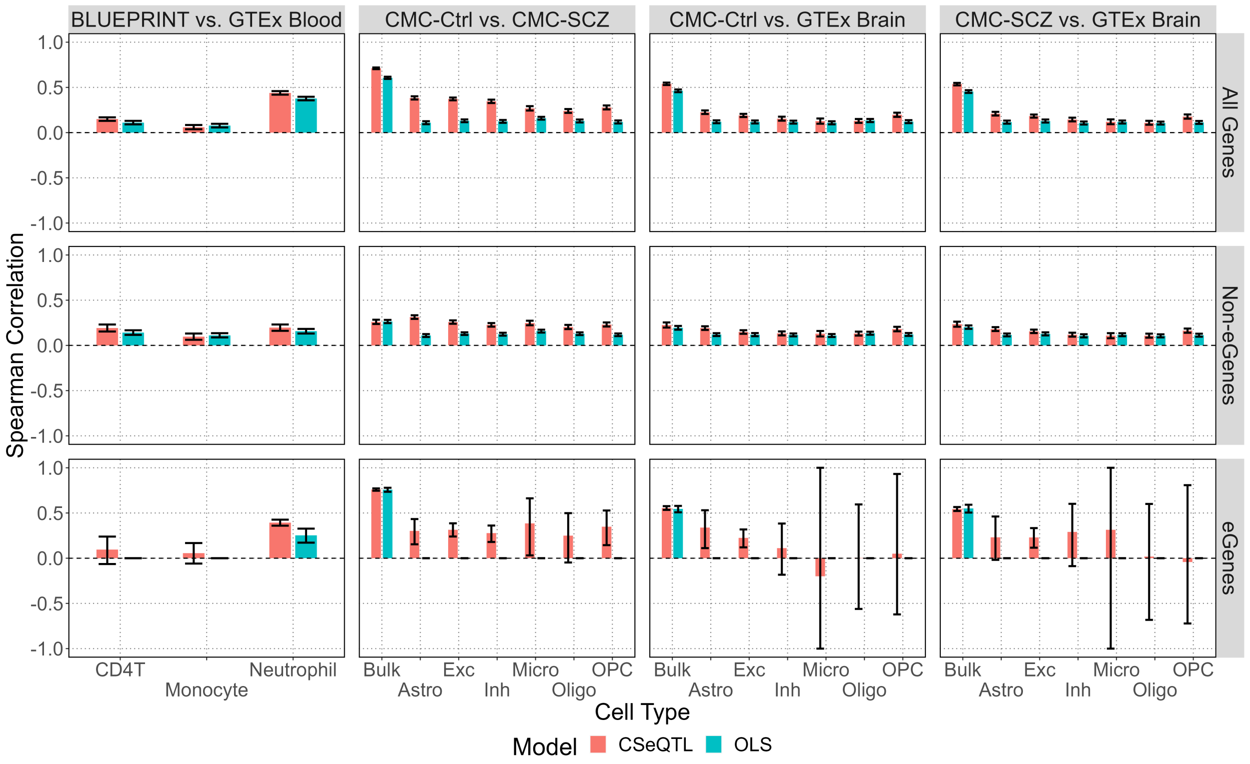

Figure 25: **Comparing findings between data cohorts for shared cell types and between models.** The y-axis corresponds to the correlation of minimum eQTL p-value per gene between a pair of cohorts. 95% confidence intervals are also plotted. For BLUEPRINT and GTEx Whole Blood, CD4 T cells, monocytes, and neutrophils are presented. For Commonmind Consortium and GTEx Brain Frontal Cortex, bulk and the six cell types are presented. Spearman correlation was calculated among all shared genes analyzed ("All Genes"), among shared eGenes ("eGenes"), and among all shared non-eGenes ("Non-eGenes").

#### 6.7 Consistency and enrichment

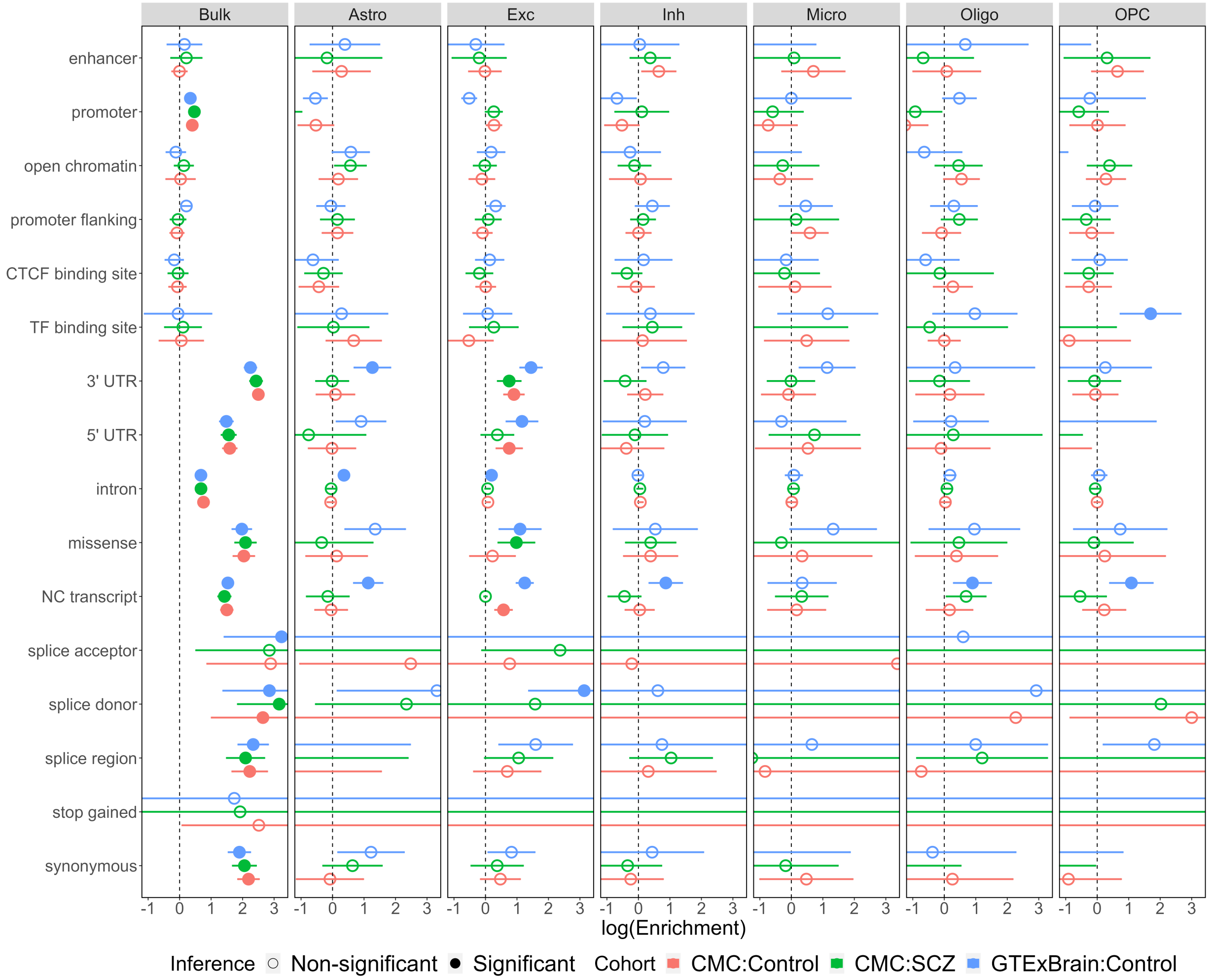

Figure 26: Functional enrichment for the eQTL results from the CMC cohorts and GTEx Brain samples. Each circle is a point estimate and the lines are 95% confidence interval. Nominal p-values are Bonferroni corrected. Filled circles correspond to the ones with lower bound of confidence intervals larger than zero and adjusted p-values $< 0.05$.

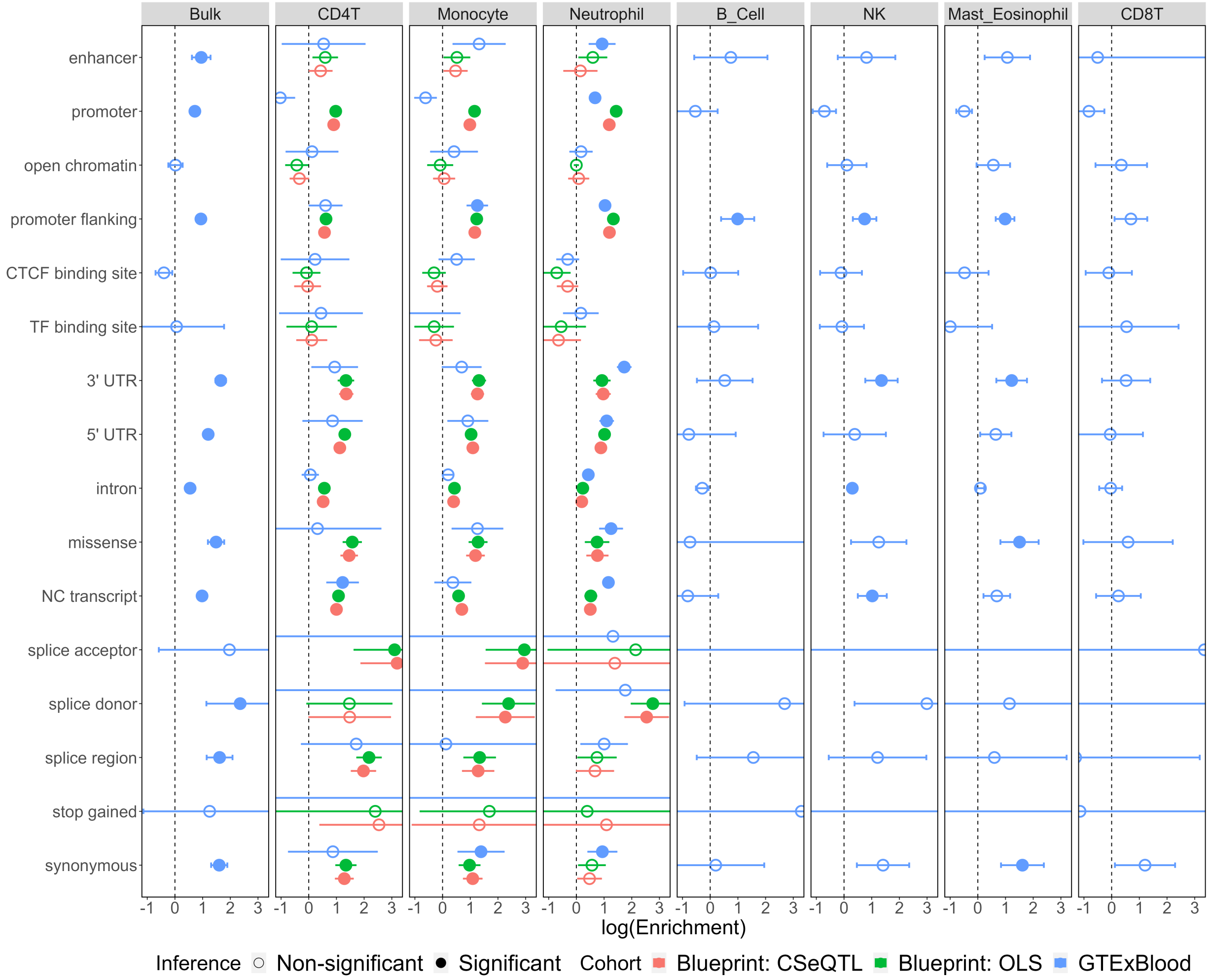

Figure 27: Functional enrichment for the eQTL results from from BLUEPRINT study and GTEx whole blood samples. Each circle is a point estimate and the lines are 95% confidence interval. Nominal p-values are Bonferroni corrected. Filled circles correspond to the ones with lower bound of confidence intervals larger than zero and adjusted p-values $< 0.05$. Note that for purified RNA-seq data that only contains one cell type, CSeQTL becomes equivalent to TReCASE.

| Comparison | MODEL | CELL TYPE | eGene1; eGene2; eGene intersection | P-value | Correlation | | |
|---|---|---|---|---|---|---|---|
| | | | | | All genes | eGenes | non-eGenes |
| CMC-Ctrl vs. CMC-SCZ | CSeQTL | Bulk | 7676; 6828; 5458 | 0.00e+00 | 0.71 (0.70, 0.72) | 0.76 (0.75, 0.77) | 0.26 (0.24, 0.28) |
| | | Astro | 644; 502; 177 | 1.90e-147 | 0.38 (0.37, 0.40) | 0.30 (0.15, 0.43) | 0.31 (0.29, 0.33) |
| | | Exc | 2246; 1640; 715 | 7.40e-213 | 0.37 (0.36, 0.39) | 0.31 (0.24, 0.39) | 0.26 (0.24, 0.28) |
| | | Inh | 1084; 889; 437 | 7.01e-315 | 0.35 (0.33, 0.36) | 0.28 (0.18, 0.36) | 0.23 (0.21, 0.25) |
| | | Micro | 197; 186; 36 | 3.92e-35 | 0.27 (0.24, 0.29) | 0.39 (0.03, 0.66) | 0.25 (0.22, 0.27) |
| | | Oligo | 381; 256; 66 | 6.96e-55 | 0.24 (0.22, 0.26) | 0.25 (-0.05, 0.50) | 0.20 (0.18, 0.23) |
| | | OPC | 375; 265; 68 | 4.34e-56 | 0.28 (0.26, 0.30) | 0.35 (0.14, 0.53) | 0.23 (0.21, 0.25) |
| | OLS | Bulk | 3707; 2872; 2370 | 0.00e+00 | 0.61 (0.59, 0.62) | 0.76 (0.73, 0.78) | 0.26 (0.25, 0.28) |
| | | Astro | 0; 0; 0 | | 0.11 (0.09, 0.13) | | 0.11 (0.09, 0.13) |
| | | Exc | 1; 0; 0 | | 0.13 (0.12, 0.15) | | 0.13 (0.12, 0.14) |
| | | Inh | 2; 0; 0 | | 0.12 (0.11, 0.14) | | 0.12 (0.11, 0.14) |
| | | Micro | 0; 0; 0 | | 0.16 (0.14, 0.17) | | 0.16 (0.14, 0.17) |
| | | Oligo | 1; 2; 1 | 1.32e-04 | 0.13 (0.11, 0.14) | | 0.13 (0.11, 0.14) |
| | | OPC | 0; 0; 0 | | 0.12 (0.10, 0.13) | | 0.12 (0.10, 0.13) |
| CMC-Ctrl vs. GTEx Brain | CSeQTL | Bulk | 7194; 7544; 5089 | 0.00e+00 | 0.54 (0.53, 0.55) | 0.55 (0.53, 0.58) | 0.23 (0.20, 0.25) |
| | | Astro | 613; 315; 82 | 1.43e-45 | 0.23 (0.21, 0.25) | 0.34 (0.11, 0.53) | 0.19 (0.17, 0.21) |
| | | Exc | 2086; 1223; 385 | 4.33e-45 | 0.19 (0.17, 0.21) | 0.22 (0.12, 0.32) | 0.15 (0.13, 0.17) |
| | | Inh | 912; 171; 51 | 2.11e-14 | 0.15 (0.13, 0.18) | 0.11 (-0.18, 0.38) | 0.13 (0.11, 0.15) |
| | | Micro | 124; 52; 4 | 5.16e-02 | 0.13 (0.10, 0.16) | -0.20 (-1.00, 1.00) | 0.13 (0.10, 0.16) |
| | | Oligo | 373; 78; 15 | 2.02e-06 | 0.13 (0.11, 0.15) | 0.00 (-0.56, 0.59) | 0.13 (0.11, 0.15) |
| | | OPC | 310; 106; 9 | 9.61e-02 | 0.20 (0.17, 0.22) | 0.05 (-0.62, 0.93) | 0.18 (0.16, 0.21) |
| | OLS | Bulk | 3447; 2260; 1668 | 0.00e+00 | 0.46 (0.45, 0.48) | 0.55 (0.51, 0.58) | 0.19 (0.17, 0.22) |
| | | Astro | 0; 0; 0 | | 0.12 (0.10, 0.14) | | 0.12 (0.10, 0.14) |
| | | Exc | 0; 2; 0 | | 0.12 (0.10, 0.13) | | 0.12 (0.10, 0.14) |
| | | Inh | 2; 0; 0 | | 0.12 (0.10, 0.13) | | 0.12 (0.10, 0.13) |
| | | Micro | 0; 0; 0 | | 0.11 (0.09, 0.13) | | 0.11 (0.09, 0.13) |
| | | Oligo | 0; 0; 0 | | 0.13 (0.12, 0.15) | | 0.13 (0.12, 0.15) |
| | | OPC | 0; 0; 0 | | 0.12 (0.10, 0.14) | | 0.12 (0.10, 0.14) |
| CMC-SCZ vs. GTEx Brain | CSeQTL | Bulk | 6324; 7472; 4657 | 0.00e+00 | 0.54 (0.52, 0.55) | 0.54 (0.52, 0.57) | 0.23 (0.21, 0.26) |
| | | Astro | 460; 310; 66 | 9.32e-40 | 0.21 (0.19, 0.23) | 0.23 (-0.02, 0.46) | 0.18 (0.16, 0.20) |
| | | Exc | 1519; 1212; 293 | 1.22e-38 | 0.18 (0.17, 0.20) | 0.23 (0.12, 0.33) | 0.16 (0.14, 0.17) |
| | | Inh | 754; 169; 39 | 3.46e-10 | 0.14 (0.12, 0.17) | 0.29 (-0.09, 0.60) | 0.12 (0.10, 0.14) |
| | | Micro | 136; 55; 6 | 3.59e-03 | 0.12 (0.09, 0.15) | 0.31 (-1.00, 1.00) | 0.11 (0.08, 0.14) |
| | | Oligo | 247; 77; 10 | 1.33e-04 | 0.11 (0.09, 0.13) | 0.02 (-0.68, 0.60) | 0.11 (0.09, 0.13) |
| | | OPC | 245; 113; 10 | 6.06e-03 | 0.18 (0.15, 0.20) | -0.04 (-0.72, 0.81) | 0.16 (0.14, 0.19) |
| | OLS | Bulk | 2596; 2227; 1469 | 0.00e+00 | 0.45 (0.44, 0.47) | 0.55 (0.50, 0.59) | 0.20 (0.18, 0.22) |
| | | Astro | 0; 0; 0 | | 0.12 (0.10, 0.13) | | 0.12 (0.10, 0.13) |
| | | Exc | 0; 2; 0 | | 0.13 (0.11, 0.15) | | 0.13 (0.11, 0.14) |
| | | Inh | 0; 0; 0 | | 0.11 (0.09, 0.12) | | 0.11 (0.09, 0.12) |
| | | Micro | 0; 0; 0 | | 0.12 (0.10, 0.13) | | 0.12 (0.10, 0.13) |
| | | Oligo | 1; 0; 0 | | 0.11 (0.09, 0.12) | | 0.11 (0.09, 0.12) |
| | | OPC | 0; 0; 0 | | 0.11 (0.10, 0.13) | | 0.11 (0.10, 0.13) |
| BLUEPRINT vs. GTEx Blood | CSeQTL | CD4T | 6520; 207; 164 | 2.35e-02 | 0.15 (0.13, 0.17) | 0.10 (-0.06, 0.24) | 0.19 (0.15, 0.23) |
| | | Monocyte | 3782; 409; 275 | 2.71e-09 | 0.06 (0.04, 0.08) | 0.06 (-0.06, 0.17) | 0.10 (0.06, 0.13) |
| | | Neutrophil | 5322; 3336; 2636 | 1.99e-160 | 0.44 (0.42, 0.46) | 0.39 (0.36, 0.43) | 0.20 (0.16, 0.23) |
| | OLS | CD4T | 3934; 0; 0 | | 0.11 (0.09, 0.13) | | 0.14 (0.12, 0.17) |
| | | Monocyte | 4229; 1; 1 | 4.02e-01 | 0.08 (0.06, 0.10) | | 0.11 (0.09, 0.14) |
| | | Neutrophil | 3154; 752; 628 | 3.54e-170 | 0.38 (0.36, 0.40) | 0.25 (0.17, 0.33) | 0.16 (0.13, 0.18) |

Table 6: **Consistency results between cohorts by model, analysis, and cell type.** Column 4 contains the number of eGenes detected per cohort and shared between cohorts. The fifth column corresponds to the Pearson $\chi^2$ test associating eGene classification between a pair of cohorts. The last three columns contain the Spearman correlation with 95% confidence intervals of minimum p-value per gene across all genes, all shared eGenes, and all shared non-eGenes, respectively.

##### 6.8 Summary of number of eQTL findings with alternative q-value cutoff

| DISEASE | MODEL | CELLTYPE | Perm. PVAL | eGenes (%all, %CIS) | overlap_Bulk | ERROR |
|---|---|---|---|---|---|---|
| Control | CSeQTL | Bulk | 3.2e-01 | 13008 (84.8%, 75%) | | 0.067 |
| | | Astro | 4.4e-03 | 1008 (6.6%, 95%) | 93% | 0.043 |
| | | Exc | 2.5e-02 | 3843 (25.3%, 90%) | 92% | 0.051 |
| | | Inh | 7.8e-03 | 1722 (11.5%, 93%) | 90% | 0.046 |
| | | Micro | 2.4e-03 | 411 (2.7%, 90%) | 93% | 0.050 |
| | | Oligo | 4.7e-03 | 764 (5%, 91%) | 92% | 0.051 |
| | | OPC | 3.1e-03 | 606 (4.1%, 95%) | 89% | 0.043 |
| | OLS | Bulk | 4.9e-02 | 5916 (38.6%, -) | | 0.050 |
| | | Astro | 1.7e-05 | 0 (-, -) | - | 0.052 |
| | | Exc | 7.1e-08 | 1 (0%, -) | 100% | 0.053 |
| | | Inh | 4.1e-07 | 2 (0%, -) | 100% | 0.052 |
| | | Micro | 4.7e-05 | 0 (-, -) | - | 0.056 |
| | | Oligo | 2.1e-05 | 4 (0%, -) | 50% | 0.052 |
| | | OPC | 2.7e-05 | 0 (-, -) | - | 0.056 |
| SCZ | CSeQTL | Bulk | 2.7e-01 | 12285 (80.4%, 75%) | | 0.070 |
| | | Astro | 3.7e-03 | 809 (5.4%, 93%) | 89% | 0.046 |
| | | Exc | 2.4e-02 | 3381 (22.3%, 89%) | 89% | 0.055 |
| | | Inh | 6.2e-03 | 1411 (9.5%, 94%) | 87% | 0.049 |
| | | Micro | 2.2e-03 | 356 (2.4%, 91%) | 84% | 0.054 |
| | | Oligo | 3.2e-03 | 506 (3.3%, 91%) | 89% | 0.054 |
| | | OPC | 2.4e-03 | 505 (3.4%, 94%) | 87% | 0.043 |
| | OLS | Bulk | 3.4e-02 | 4675 (30.6%, -) | | 0.050 |
| | | Astro | 1.1e-04 | 0 (-, -) | - | 0.058 |
| | | Exc | 3.0e-05 | 4 (0%, -) | 100% | 0.058 |
| | | Inh | 8.8e-06 | 0 (-, -) | - | 0.056 |
| | | Micro | 1.7e-05 | 0 (-, -) | - | 0.061 |
| | | Oligo | 1.5e-05 | 6 (0%, -) | 50% | 0.053 |
| | | OPC | 2.2e-05 | 0 (-, -) | - | 0.056 |

Table 7: CMC eQTL results given q-value cutoff 0.05. “Perm. PVAL” is the permutation p-value cutoff for q-value 0.05. eGenes (%all, %CIS) is the number of eGenes, the corresponding percentage of all the genes being eGenes, and the percentage of genes whose most significant eQTL is a cis-eQTLs (defined as those with cis-trans test p-value $> 0.01$). overlap_Bulk is the percent of eGenes that are also eGenes from bulk eQTL analysis. ERROR is the proportion of eQTLs with p-value smaller than 0.05 using permuted data. All the results are after trimming.

| MODEL | CELLTYPE | Perm. PVAL | eGenes (%all, %CIS) | overlap_Bulk | ERROR |
|---|---|---|---|---|---|
| CSeQTL | Bulk | 5.3e-01 | 14812 (96.2%, 72%) | | 0.082 |
| | Astro | 6.2e-03 | 1006 (6.6%, 85%) | 99% | 0.063 |
| | Exc | 3.4e-02 | 3590 (23.5%, 82%) | 97% | 0.068 |
| | Inh | 2.6e-03 | 440 (3%, 83%) | 98% | 0.058 |
| | Micro | 1.1e-03 | 193 (1.3%, 69%) | 96% | 0.056 |
| | Oligo | 1.7e-03 | 353 (2.4%, 72%) | 98% | 0.065 |
| | OPC | 1.2e-03 | 257 (1.8%, 81%) | 96% | 0.050 |
| OLS | Bulk | 2.8e-02 | 4159 (27%, -) | | 0.051 |
| | Astro | 1.1e-04 | 0 (-, -) | - | 0.043 |
| | Exc | 1.5e-05 | 7 (0%, -) | 100% | 0.050 |
| | Inh | 4.6e-05 | 0 (-, -) | - | 0.049 |
| | Micro | 1.3e-05 | 0 (-, -) | - | 0.042 |
| | Oligo | 1.2e-05 | 0 (-, -) | - | 0.046 |
| | OPC | 6.8e-05 | 0 (-, -) | - | 0.045 |

Table 8: GTEx Brain Frontal Cortex (BA9) eQTL results for q-value cutoff 0.05. The meaning of the columns are the same as the columns of Table 7.

| MODEL | CELLTYPE | Perm. PVAL | eGenes (%all, %CIS) | ERROR |
|---|---|---|---|---|
| CSeQTL | | 7.2e-01 | 13256 (99.4%, 70%) | 0.084 |
| | | 1.7e-01 | 10192 (74.8%, 67%) | 0.074 |
| | | 3.0e-01 | 9310 (86.5%, 64%) | 0.084 |
| OLS | | 8.6e-02 | 7256 (54.4%, -) | 0.050 |
| | | 9.0e-02 | 7611 (55.9%, -) | 0.051 |
| | | 7.8e-02 | 5608 (52.1%, -) | 0.051 |

Table 9: BLUEPRINT eQTL results for q-value cutoff 0.05. The meaning of the columns are the same as the columns of Table 7.

| MODEL | CELLTYPE | Perm. PVAL | eGenes (%all, %CIS) | overlap_Bulk | ERROR |
|---|---|---|---|---|---|
| CSeQTL | Bulk | 5.2e-01 | 11517 (95.8%, 80%, -) | | 0.063 |
| | CD4T | 1.7e-03 | 331 (2.9%, 95%, 0%) | 98% | 0.040 |
| | CD8T | 1.9e-03 | 317 (2.7%, 97%, 0%) | 98% | 0.044 |
| | B_Cell | 1.4e-03 | 203 (1.8%, 92%, 0.1%) | 98% | 0.045 |
| | Monocyte | 5.6e-03 | 678 (5.8%, 86%, 0.4%) | 99% | 0.056 |
| | NK | 4.4e-03 | 710 (6.1%, 88%, 0.2%) | 99% | 0.053 |
| | Neutrophil | 9.7e-02 | 6545 (54.6%, 85%, 0.6%) | 98% | 0.066 |
| | Mast_Eosinophil | 1.4e-02 | 1546 (13.1%, 75%, 1.2%) | 97% | 0.073 |
| OLS | Bulk | 1.1e-01 | 7426 (61.9%, -, -) | | 0.050 |
| | CD4T | 3.0e-05 | 0 (-, -, -) | - | 0.044 |
| | CD8T | 4.2e-05 | 0 (-, -, -) | - | 0.049 |
| | B_Cell | 1.6e-05 | 0 (-, -, -) | - | 0.052 |
| | Monocyte | 1.3e-05 | 1 (0%, -, -) | 100% | 0.059 |
| | NK | 3.0e-05 | 0 (-, -, -) | - | 0.055 |
| | Neutrophil | 1.6e-02 | 1882 (16%, -, -) | 92% | 0.060 |
| | Mast_Eosinophil | 1.9e-04 | 28 (0.2%, -, 0%) | 75% | 0.067 |

Table 10: GTEx Whole Blood eQTL results for q-value cutoff 0.05. The meaning of the columns are the same as the columns of Table 7.

#### 6.9 Comparison of eQTL effect directions

| Cell type | CSeQTL q-value cutoff | number of eQTLs | Consistency prop. expected | Consistency prop. observed | Chi-squared test pval |
|---|---|---|---|---|---|
| All cell types | none | 72,790 | 0.50 | 0.51 | |
| CD4T | 0.1 | 156 | 0.51 | 0.88 | 1.9e-20 |
| CD4T | 0.005 | 98 | 0.52 | 0.92 | 8.3e-16 |
| CD8T | 0.1 | 137 | 0.57 | 0.9 | 2.9e-18 |
| CD8T | 0.005 | 71 | 0.62 | 0.92 | 4.1e-10 |
| B | 0.1 | 51 | 0.55 | 0.88 | 4.9e-07 |
| B | 0.005 | 31 | 0.53 | 0.9 | 3.6e-05 |
| NK | 0.1 | 212 | 0.5 | 0.94 | 1.4e-36 |
| NK | 0.005 | 145 | 0.51 | 0.96 | 1.9e-27 |
| Monocyte | 0.1 | 159 | 0.5 | 0.89 | 1.3e-22 |
| Monocyte | 0.005 | 121 | 0.5 | 0.91 | 1.3e-18 |
| CD4NC | 0.1 | 90 | 0.51 | 0.87 | 1.3e-11 |
| CD4NC | 0.005 | 54 | 0.51 | 0.91 | 1.4e-08 |
| CD4ET | 0.1 | 66 | 0.52 | 0.89 | 1.2e-09 |
| CD4ET | 0.005 | 44 | 0.52 | 0.93 | 5.5e-08 |
| CD8NC | 0.1 | 43 | 0.58 | 0.86 | 4.9e-05 |
| CD8NC | 0.005 | 21 | 0.59 | 0.9 | 0.0029 |
| CD8ET | 0.1 | 61 | 0.56 | 0.89 | 4.8e-08 |
| CD8ET | 0.005 | 31 | 0.63 | 0.9 | 0.00029 |
| CD8S100B | 0.1 | 33 | 0.59 | 0.97 | 9.1e-07 |
| CD8S100B | 0.005 | 19 | 0.64 | 0.95 | 0.0018 |
| BIN | 0.1 | 26 | 0.55 | 0.88 | 0.00054 |
| BIN | 0.005 | 16 | 0.56 | 0.88 | 0.017 |
| BMem | 0.1 | 25 | 0.55 | 0.88 | 0.0012 |
| BMem | 0.005 | 15 | 0.51 | 0.93 | 0.0043 |
| NKR | 0.1 | 42 | 0.51 | 1 | 7e-10 |
| NKR | 0.005 | 33 | 0.51 | 1 | 7.1e-08 |
| NK | 0.1 | 170 | 0.51 | 0.92 | 1.7e-27 |
| NK | 0.005 | 112 | 0.53 | 0.95 | 4.3e-20 |
| MonoC | 0.1 | 84 | 0.5 | 0.89 | 2e-12 |
| MonoC | 0.005 | 64 | 0.5 | 0.91 | 3.8e-10 |
| MonoNC | 0.1 | 75 | 0.53 | 0.89 | 9.1e-11 |
| MonoNC | 0.005 | 57 | 0.52 | 0.91 | 3.8e-09 |

Table 11: Comparison of eQTL effect directions between CSeQTL results from GTEx whole blood (q-value $< 0.1$ or 0.005 except the first row where no filtering was conducted) and scRNA-seq eQTL results from Yazar et al. [27] (p-value $< 0.01$ except for the first row). We considered either the cell types used in CSeQTL or the cell types used by Yazar et al. [27].

| Cell type | CSeQTL q-value cutoff | number of eQTLs | Consistency prop. expected | observed | Chi-squared test pval |
|---|---|---|---|---|---|
| All cell types | none | 39,940 | 0.50 | 0.53 | |
| Astro | 0.1 | 149 | 0.52 | 0.96 | 2.9e-28 |
| Astro | 0.005 | 89 | 0.55 | 0.97 | 1.6e-17 |
| Exc | 0.1 | 547 | 0.54 | 0.91 | 2.1e-80 |
| Exc | 0.005 | 284 | 0.53 | 0.93 | 1.6e-46 |
| Inh | 0.1 | 18 | 0.52 | 0.78 | 0.065 |
| Inh | 0.005 | 9 | 0.52 | 0.89 | 0.097 |
| Micro | 0.1 | 13 | 0.51 | 0.92 | 0.012 |
| Micro | 0.005 | 9 | 0.52 | 0.89 | 0.097 |
| Oligo | 0.1 | 62 | 0.51 | 0.85 | 1.3e-07 |
| Oligo | 0.005 | 27 | 0.52 | 0.96 | 1.1e-05 |
| OPC | 0.1 | 15 | 0.51 | 0.93 | 0.0043 |
| OPC | 0.005 | 9 | 0.59 | 0.89 | 0.16 |

Table 12: Comparison of eQTL effect directions between CSeQTL results from GTEx brain and scRNA-seq eQTL results from Bryois et al. [28]. The p-value/q-value cutoffs are the same as Table 11.

| Cell type | CSeQTL q-value cutoff | number of eQTLs | Consistency prop. expected | observed | Chi-squared test pval |
|---|---|---|---|---|---|
| All cell types | none | 35,161 | 0.50 | 0.52 | |
| Astro | 0.1 | 53 | 0.53 | 0.91 | 3.5e-08 |
| Astro | 0.005 | 41 | 0.53 | 0.93 | 3.7e-07 |
| Exc | 0.1 | 378 | 0.54 | 0.88 | 6.8e-47 |
| Exc | 0.005 | 229 | 0.55 | 0.93 | 1e-35 |
| Inh | 0.1 | 42 | 0.53 | 0.6 | 0.58 |
| Inh | 0.005 | 27 | 0.51 | 0.63 | 0.32 |
| Micro | 0.1 | 11 | 0.52 | 0.82 | 0.12 |
| Micro | 0.005 | 7 | 0.53 | 0.86 | 0.28 |
| Oligo | 0.1 | 61 | 0.5 | 0.89 | 9.2e-09 |
| Oligo | 0.005 | 28 | 0.54 | 0.93 | 3.9e-05 |
| OPC | 0.1 | 17 | 0.56 | 0.65 | 0.73 |
| OPC | 0.005 | 13 | 0.63 | 0.69 | 1.00 |

Table 13: Comparison of eQTL effect directions between CSeQTL results from CMC schizophrenia patients and scRNA-seq eQTL results from Bryois et al. [28]. The p-value/q-value cutoffs are the same as Table 11.

| Cell type | CSeQTL q-value cutoff | number of eQTLs | Consistency prop. expected | observed | Chi-squared test pval |
|---|---|---|---|---|---|
| All cell types | none | 34,321 | 0.50 | 0.52 | |
| Astro | 0.1 | 74 | 0.5 | 0.89 | 7.1e-11 |
| Astro | 0.005 | 55 | 0.5 | 0.93 | 1.2e-09 |
| Exc | 0.1 | 411 | 0.52 | 0.83 | 1.2e-39 |
| Exc | 0.005 | 264 | 0.51 | 0.85 | 4.3e-30 |
| Inh | 0.1 | 58 | 0.5 | 0.67 | 0.017 |
| Inh | 0.005 | 35 | 0.54 | 0.69 | 0.092 |
| Micro | 0.1 | 22 | 0.51 | 0.82 | 0.011 |
| Micro | 0.005 | 15 | 0.53 | 0.8 | 0.094 |
| Oligo | 0.1 | 69 | 0.57 | 0.84 | 4.9e-07 |
| Oligo | 0.005 | 41 | 0.55 | 0.83 | 0.00017 |
| OPC | 0.1 | 15 | 0.58 | 0.8 | 0.15 |
| OPC | 0.005 | 10 | 0.68 | 1 | 0.03 |

Table 14: Comparison of eQTL effect directions between CSeQTL results from CMC controls and scRNA-seq eQTL results from Bryois et al. [28]. The p-value/q-value cutoffs are the same as Table 11.

[12] Jesse Krijthe, Laurens van der Maaten, and Maintainer Jesse Krijthe. Package 'rtsne', 2018.

[13] Greg Finak, Andrew McDavid, Masanao Yajima, Jingyuan Deng, Vivian Gersuk, Alex K Shalek, Chloe K Slichter, Hannah W Miller, M Juliana McElrath, Martin Prlic, et al. MAST: a flexible statistical framework for assessing transcriptional changes and characterizing heterogeneity in single-cell RNA sequencing data. *Genome biology*, 16(1):278, 2015.

[14] Wei Sun. A statistical framework for eQTL mapping using RNA-seq data. *Biometrics*, 68(1):1–11, 2012.

[15] Yi-Juan Hu, Wei Sun, Jung-Ying Tzeng, and Charles M Perou. Proper use of allele-specific expression improves statistical power for cis-eQTL mapping with RNA-seq data. *Journal of the American Statistical Association*, 110(511):962–974, 2015.

[16] Vasyl Zhabotynsky, Licai Huang, Paul Little, Yi-Juan Hu, Fernando Pardo-Manuel de Villena, Fei Zou, and Wei Sun. eQTL mapping using allele-specific count data is computationally feasible, powerful, and provides individual-specific estimates of genetic effects. *PLOS Genetics*, in press, 2022.

[23] Picard toolkit, 2019.

[24] Wei Sun, Vasyl Zhabotynsky, and Chong Jin. *asSeq: allele specific sequencing study*, 2018. R package version 0.99.501.
